# Reverse regression increases power for detecting trans-eQTLs

**DOI:** 10.1101/2020.05.07.083386

**Authors:** Saikat Banerjee, Franco L. Simonetti, Kira E. Detrois, Anubhav Kaphle, Raktim Mitra, Rahul Nagial, Johannes Söding

**Affiliations:** Quantitative and Computational Biology, Max-Planck Institute for Biophysical Chemistry, 37077 Göttingen, Germany; Georg-August University, 37075 Göttingen, Germany; Indian Institute of Technology, Kanpur, India; Campus-Institut Data Science (CIDAS), University of Göttingen, Germany; Cluster of Excellence “Multiscale Bioimaging” (MBExC), University of Göttingen, Germany

## Abstract

Trans-acting expression quantitative trait loci (trans-eQTLs) are genetic variants affecting the expression of distant genes. They account for ≥70% expression heritability and could therefore facilitate uncovering mechansisms underlying the origination of complex diseases. However, unlike cis-eQTLs, identifying trans-eQTLs is challenging because of small effect sizes, tissue-specificity, and the severe multiple-testing burden. Trans-eQTLs affect multiple target genes, but aggregating evidence over individual SNP-gene associations is hampered by strong gene expression correlations resulting in correlated p-values. Our method Tejaas predicts trans-eQTLs by performing L_2_-regularized ‘reverse’ multiple regression of each SNP on all genes, aggregating evidence from many small trans-effects while being unaffected by the strong expression correlations. Combined with a novel non-linear, unsupervised k-nearest-neighbor method to remove confounders, Tejaas predicted 18851 unique trans-eQTLs across 49 tissues from GTEx. They are enriched in open chromatin, enhancers and other regulatory regions. Many overlap with disease-associated SNPs, pointing to tissue-specific transcriptional regulation mechanisms. Tejaas is available under GPL at https://github.com/soedinglab/tejaas.

## Introduction

The detection, prevention and therapeutics of complex diseases, such as atherosclerosis, Alzheimer’s disease or schizophrenia, can improve with better understanding of the genetic pathways underlying these diseases. Over the last decade, genome-wide association studies (GWASs) have identified tens of thousands of bona fide genetic loci associated with complex traits and diseases. However, it remains unclear how most of the disease-associated variants exert their effects and influence disease risk. Over 90% of the GWAS variants are single-nucleotide polymorphisms (SNPs) in noncoding regions [1], potentially regulating gene expression that influence disease risk. Indeed, eQTL mapping has identified many genetic variants that affect gene expression. These have been mostly limited to cis-eQTLs, which modulate the expression of proximal genes (usually within ±1 Mbp), while little is known about trans-eQTLs, which modulate distal genes or those residing on different chromosomes.

The discovery of trans-eQTLs is critical to advance our understanding of causative disease pathways because they account for a large proportion of the heritability of gene expression. Several recent studies converge on an estimate of 60%-90% genetic variance in gene expression contributed by trans-eQTLs and only 10%-40% by cis-eQTLs (see Table 1 in [2] for an overview).

However, in contrast to cis-eQTLs, trans-eQTLs are notoriously difficult to discover. The standard method involves simple regression of each gene on all SNPs. For cis-eQTLs, the number of association tests is limited to SNPs in the vicinity of each gene, while for trans-eQTLs, testing all genes against all SNPs imposes a hefty multiple testing burden. The major impediment, however, comes from the small effect sizes of trans-eQTLs on individual genes. Moreover, combining signals across multiple tissues is hindered by the tissue-specificity of trans-eQTLs.

Several studies searched for trans-eQTLs among restricted sets to reduce the multiple testing burden; for instance among trait-associated SNPs [3] or among SNPs with significant cis-associations [4]. A few methods have been developed to find trans-eQTLs using distinctive biological signatures. For example, GNetLMM [5] implicitly assumes that a trans-eQTL targets a trans-eGene via an intermediate cis-eGene. Their method tests for association between the SNP and the candidate gene using a linear mixed model, while conditioning on another set of genes that affect the candidate gene but are uncorrelated to the cis-eGene. Another method [6] used tensor decomposition to succinctly encode the behavior of coregulated gene networks with latent components that represent the major modes of variation in gene expression across patients and tissues, testing for association between SNPs and the latent components. A class of methods using mediation analysis try to identify the genetic control points or cis-mediators of gene co-expression networks [7–9]. These methods regress the candidate trans-eGene on the cis-eGene (not on the SNP) by adaptively selecting for potential confounding variables using the SNP as an “instrumental variable”. More recently, a method for imputing gene expression was used to learn and predict each gene’s expression from its cis-eQTLs, and then the observed gene expressions were tested for association with the predicted gene expressions to find trans-eGenes [10].

Trans-eQTLs are believed to affect the expression of a proximal diffusible factor such as a transcription, RNA-binding or signaling factor, chromatin modifier, or possibly a non-coding RNA, which in turn directly or indirectly affects the expression of the trans genes [11]. It is therefore expected that trans-eQTLs affect tens or hundreds of target genes in trans. Many examples in humans (see, e.g. [12, 13]) and strong evidence in yeast [14] support this hypothesis. If this information could be used effectively to predict trans-eQTLs, it might easily compensate their weaker effect sizes and multiple testing burden.

We expect the target genes to have more significant *p*-values for association with their trans-eQTL than expected by chance. Brynedal *et al.* [15] presented a method (CPMA) that tests whether the distribution of regression *p*-values for association of the candidate SNP with each gene expression level has an excess of low *p*-values arising from the association of the target genes with the SNP. However, the *p*-values inherit the strong correlation from their gene expressions. Therefore, if one gene has a *p*-value near zero by chance, many strongly correlated genes will also have very low *p*-values. This makes it difficult to decide if an enrichment of *p*-values near zero is due to trans genes or due to chance, diminishing the power of the method significantly.

Here, we circumvent the problem by reversing the direction of regression (Fig. 1). Instead of regressing each expression level on the SNP’s minor allele count, Tejaas performs multiple regression of the SNP on *all* genes jointly. In this way, no matter how strong the correlations, they do not negatively impact the test for association between gene expressions and SNP. This approach brings two decisive advantages: First, the information from each and every target gene is accumulated while automatically taking their redundancy through correlations into account. Therefore, the more target genes a SNP has, the more sensitive Tejaas will be, even when individual effect sizes are much below the significance level for individual gene-SNP association tests. Second, the multiple testing burden is reduced because association is tested for all genes at once. To correct for known and unknown confounder variables, we present a novel nonlinear, nonparametric K-nearest neighbor correction and demonstrate its effectiveness in simulations.

**Fig. 1.**
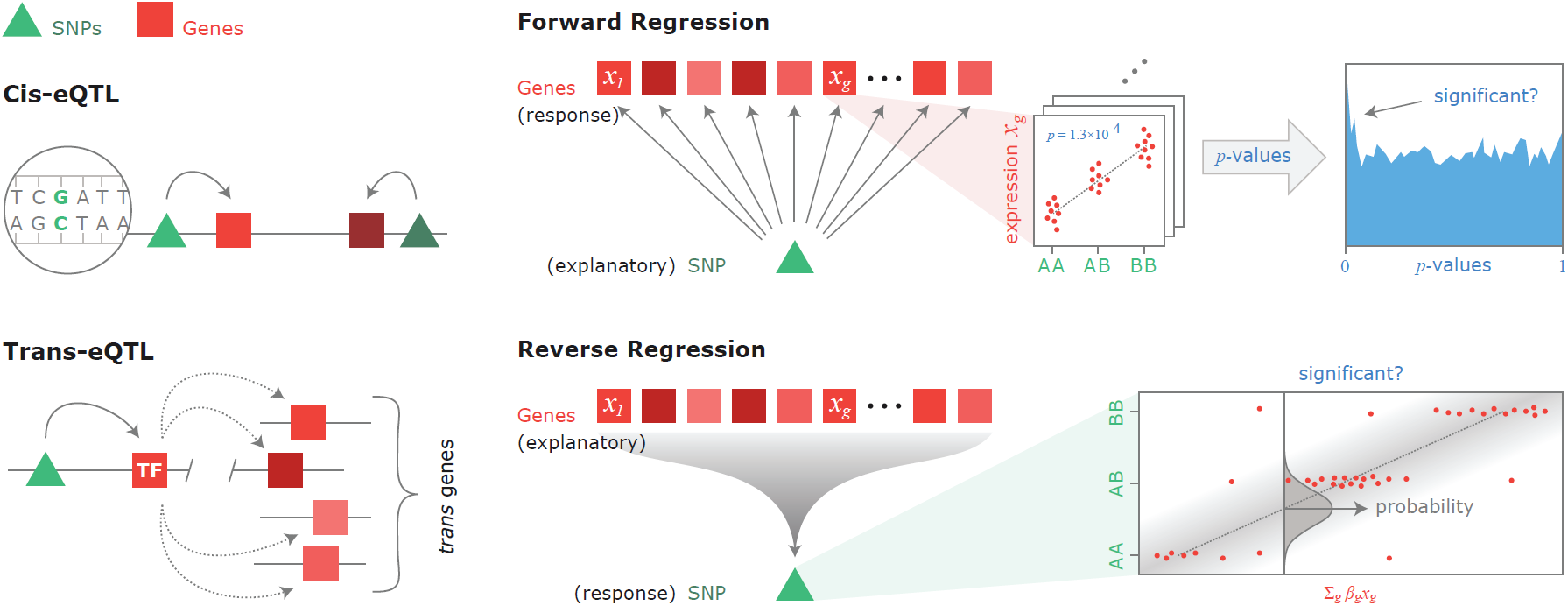
Forward and reverse regression for trans-eQTL discovery. Trans-eQTLs affect multiple genes simultaneously by exerting a cis-effect on a diffusible trans-acting factor such as a transcription factor (TF) (left). In forward regression (FR), we perform univariate regression of the expression level of each gene individually on the candidate SNP’s genotype (= centered minor allele frequency) and estimate whether the distribution of resulting association *p*-values is enriched near zero. In reverse regression (Tejaas), we perform L_2_-regularized multiple regression of the candidate SNP’s genotype jointly on all gene expression levels. Crucially, reverse regression is not negatively affected by correlations between gene expression levels.

We applied Tejaas to the Genotype Tissue Expression (GTEx) dataset and predicted 18851 trans-eQTLs in 49 tissues with a *p*-value threshold for genome-wide significance of *p* < 5 × 10^−8^, which corresponds to false discovery rates below 5%. These putative trans-eQTLs are significantly enriched in various functional genomic signatures such as chromatin accessibility, functional histone marks and reporter assay annotations, and are also enriched among GWAS SNPs associated to various complex traits.

## Results

### Methods overview

Tejaas (**T**rans-**E**QTLs by **J**oint **A**ssociation **A**nalysi**S**) computes the Reverse Regression **RR-score** *q*_rev_ to discover and rank trans-eQTLs, making use of the expectation that each trans-eQTL has multiple target genes. To our knowledge, only one other method makes use of it, the “forward” regression method CPMA by Brynedal *et al.* [15]. In order to compare Tejaas with CPMA, we implemented our own version of Forward Regression (FR) within Tejaas, as there is no publicly available software for CPMA. We used MatrixEQTL [16] as representative of all methods using single SNP-gene regression.

The FR-score *q*_fwd_ and the RR-score *q*_rev_ are summarized in Fig. 1. For details, see Online Methods, Supplementary Sec. 1 and 2. The FR score evaluates the distribution of the *p*-values for the pairwise linear association of a candidate SNP with each of the *G* gene expression levels. SNPs without trans-effect should have uniformly distributed *p*-values, while we expect trans-eQTLs to have a distribution that is enriched near zero, contributed by their target genes.

Reverse Regression (RR) performs a multiple linear regression using expression levels of all genes to explain the genotype of a candidate SNP. Let x denote the vector of centered minor allele counts of a SNP for *N* samples and Y be the *G* × *N* matrix of preprocessed expression levels for *G* genes. We model x with a normal distribution whose mean depends linearly on the gene expression through a vector of regression coefficients *β*:

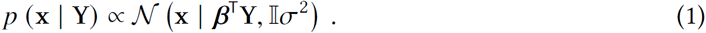

Generally, the number of explanatory variables (genes) is much larger than the number of samples (*G* ≫ *N*) in currently available eQTL data sets. To avoid overfitting, we introduce a normal prior on *β*, with mean 0 and variance *γ*^2^,

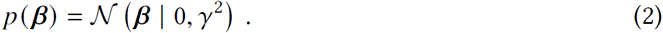

This *L*_2_ regularization pushes the effect size of non-target genes towards zero. We calculated the significance of the trans-eQTL model (*β* ≠ 0) compared to the null model (*β* = 0) using Bayes theorem to obtain

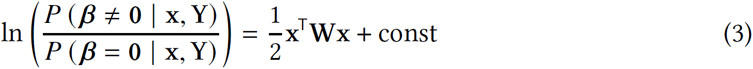

with

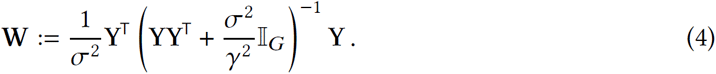

We therefore defined the RR-score as *q*_rev_ := x^T^Wx.

The null distribution of *q*_rev_ is different for every SNP and can be obtained by randomly permuting the sample labels of the genotype multiple times. Although it is computationally infeasible to obtain the null distribution empirically for each SNP independently, we were able to analytically calculate the expectation and variance of *q*_rev_ under this permuted null model (Supplementary Appendix 1). Assuming that the null distribution is Gaussian, which holds well in practice (Supplementary Sec. 2.6 and Fig. S1), we calculate a *p*-value to get the significance of any observed *q*_rev_.

The assumption of normality of the RR-score null distribution breaks down when standard confounder correction methods are used (Supplementary Fig. S2, Sec. 2.6 and Sec. 3.1). Therefore, we developed a novel, non-parametric, non-linear confounder correction using k-nearest neighbors, which we call KNN correction (Supplementary Sec. 3.2). The KNN correction does not require the confounders to be known but efficiently corrects for both hidden and known confounders (Supplementary Fig. S4, Sec. 5.4 and Fig. S9).

Tejaas is a fast and efficiently MPI-parallelized software (Supplementary Fig. S3) written in Python and C++. It is open-source and released under GNU General Public License v3 (Code Availability).

### Simulation studies

We applied Tejaas reverse regression, FR and MatrixEQTL on semi-synthetic datasets to compare their performance in well-defined settings. The simulations also allowed us to find optimum values for the number of nearest neighbors *K* and the effect size variance *γ*^2^.

For simulations, we followed the strategy of Hore *et al.* [6] (Online Methods and Supplementary Sec. 4). Briefly, for each simulation with 12 639 SNPs and 12 639 genes, we randomly selected 800 SNPs as cis-eQTLs, out of which 30 were also trans-eQTLs. The cis target genes of the trans-eQTLs were considered as transcription factors (TFs) and regulated multiple target genes downstream. Some strategies were different from the work of Hore *et al.*to make the simulations more realistic. First, we sampled the genotype directly from real data. Second, we used the covariance matrix of real gene expression as the background noise for the synthetic gene expression. Third, we included the first three genotype principal components as confounders to mimic population substructure. We measured the performance in predicting the planted trans-eQTLs by the partial area under the ROC curve (pAUC) up to a false positive rate (FPR) of 0.1.

Figure 2a shows how the pAUC is affected by three confounder correction methods: (1) without any confounder correction (None), (2) the de facto standard method using residuals after linear regression with known confounders (CCLM) and (3) the K-nearest-neighbor correction (KNN). For Tejaas, we set *γ* = 0.2 and *K* = 30 empirically (Supplementary Figs. S5–7). To avoid false discovery of cis-eQTLs as trans-eQTLs, we masked all cis genes within ±1Mb of each candidate SNP for Tejaas and Forward Regression (Supplementary Sec. 2.9).

**Fig. 2.**
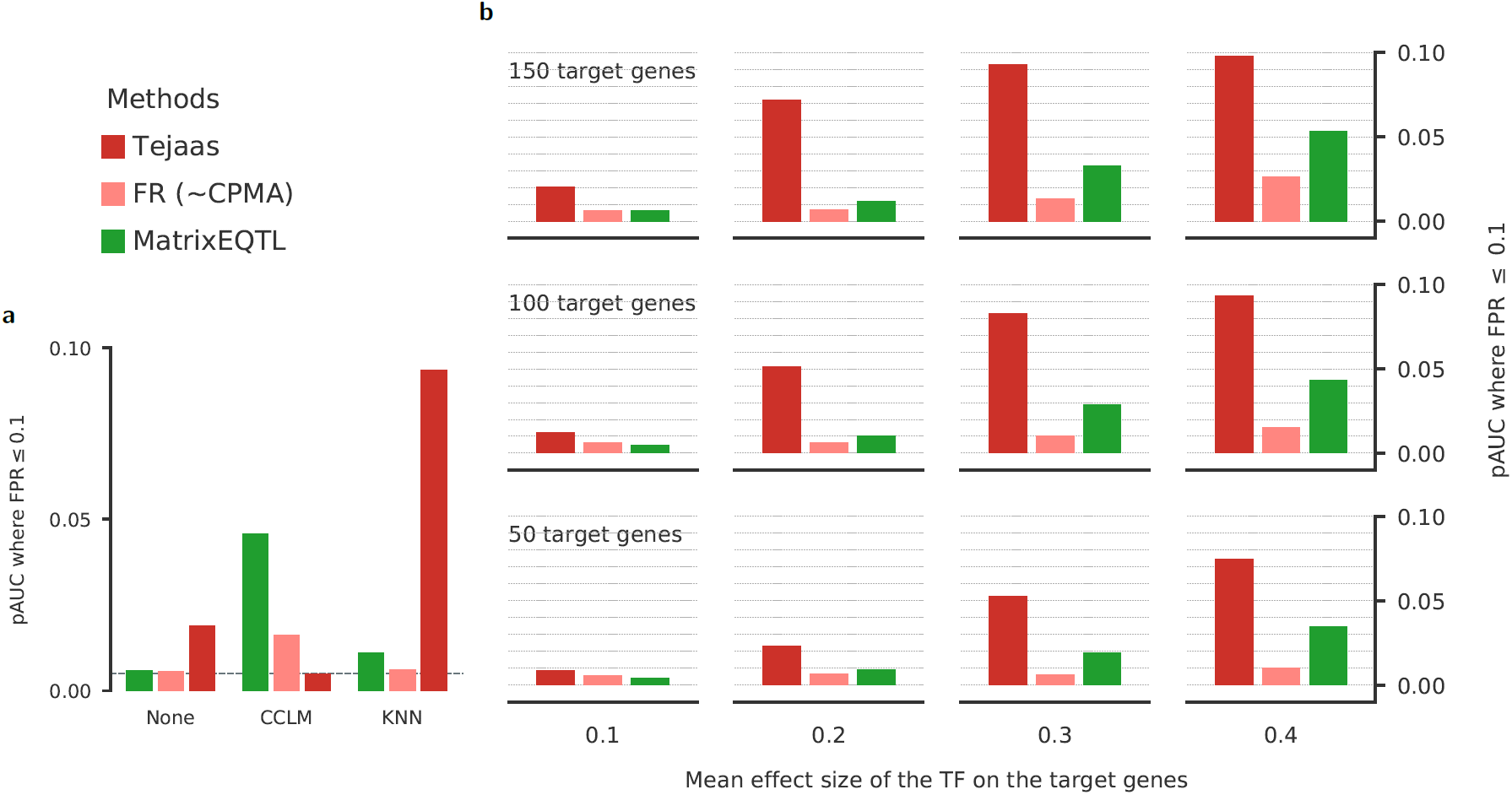
Sensitivity for trans-eQTL discovery on simulated data. We compared the performance of Tejaas reverse regression, forward regression (FR) (similar to CPMA) and MatrixEQTL, by computing the partial area under the ROC curve (pAUC) up to a false positive rate (FPR) of 0.1. A perfect method has pAUC=0.1 and a random one 0.005. pAUCs are averaged over 20 simulations. **a**, pAUC for different confounder correction methods: no correction (None), correction using linear regression of known confounders (CCLM) on inverse normal transformed gene expression, and our k-nearest neighbors correction with *K*=30 (KNN). **b**, pAUC for different numbers of target genes for the cis transcription factor (TF) mediating the trans-eQTL (from top to bottom) and different mean effect sizes of the TF on the target genes (from left to right).

The best combination of method and confounder correction is Tejaas with KNN correction (Fig. 2a). CCLM is effective for MatrixEQTL but it does not work in combination with Tejaas because it renders the null *q*_rev_ distribution non-Gaussian and thereby leads to wrong *p*-values (Supplementary Fig. S2, Sec. 2.6 and Sec. 3.1). For FR and MatrixEQTL, CCLM works much better than KNN because we provided it with the known confounders, whereas KNN did not and can not use these. Unlike in simulations, we do not have exact knowledge of most of the confounders in real data. Hence it is encouraging that the KNN correction works well even without knowledge of the confounders.

In Fig. 2b, we analyzed the methods’ performance depending on (1) the number of target genes of the TF linked to the trans-eQTL and (2) the effect size of the TF on the target genes. For MatrixEQTL and FR, we chose the CCLM correction and for Tejaas, the KNN correction. Surprisingly, FR has slightly lower pAUC than MatrixEQTL throughout. The pAUC of Tejaas is at least two-fold higher than the next best method under all conditions, although it does not use the known confounders. At mean effect size 0.2, the pAUC is up to 5 times higher than that of MatrixEQTL. The higher pAUC of Tejaas than other methods is persistent when varying the number of confounders and the effect size of confounders (Supplementary Fig. S8).

### Genotype Tissue Expression trans-eQTL analysis

We applied Tejaas to data from the Genotype Tissue Expression (GTEx) project [17–19]. The GTEx project aims to provide insights into mechanisms of gene regulation by collecting RNA-Seq gene expression measurements from 54 tissues in hundreds of human donors, of which we used 49 tissues that have ≥70 samples with both genotype and expression measurements.

We downloaded GTEx v8 data (Data Availability), converted the gene expression read counts obtained from phASER to standardized TPMs (Transcripts per Millions), and used the KNN correction with 30 nearest neighbors to remove confounders (Supplementary Sec. 5). Using a small hold-out test set for adipose subcutaneous tissue, we obtained *γ* = 0.1. We noticed that in four tissues, this choice led to non-Gaussian distribution1 s of *q*_rev_ on null SNPs. A systematic analysis of the non-Gaussianity led to a choice of *γ* = 0.006 for these remaining four tissues (Supplementary Sec. 5.5 and Fig. S10). For each candidate SNP, we removed all cis genes within ±1Mbp to avoid detecting the relatively stronger cis-eQTL signals and thereby inflating *q*_rev_ (Supplementary Fig. S12). All SNPs with a genome-wide significant *p*-value (*p* ≤ 5 × 10^−8^) were predicted as trans-eQTLs. To reduce redundancy, we pruned the list by retaining only the trans-eQTLs with lowest *p*-values in each independent LD region defined by SNPs with *r*^2^ > 0.5.

The LD-pruned lists contain 16 929 unique lead trans-eQTLs in non-brain GTEx tissues and 1 922 in brain tissues (Fig. 3a). For comparison, the latest analysis by the GTEx consortium on the the same data yielded 142 trans-eQTLs across 49 tissues analyzed at 5% false discovery rate (FDR), of which 41 were observed in testis [4]. To get a rough estimate of our FDRs at the cut-off *p*-value of 5 × 10^−8^, we note that the expectation value of the number of false positive predictions for 8 × 10^6^ tested SNPs per tissue is about 0.4, and even less after LD-pruning. Hence for a tissue with *T* predicted trans-eQTLs below the cut-off *p*-value, the FDR should be roughly ≤0.4 /*T*. It follows that 47 out of 49 tissues have FDRs at cut-off below 5% with many much below that.

**Fig. 3.**
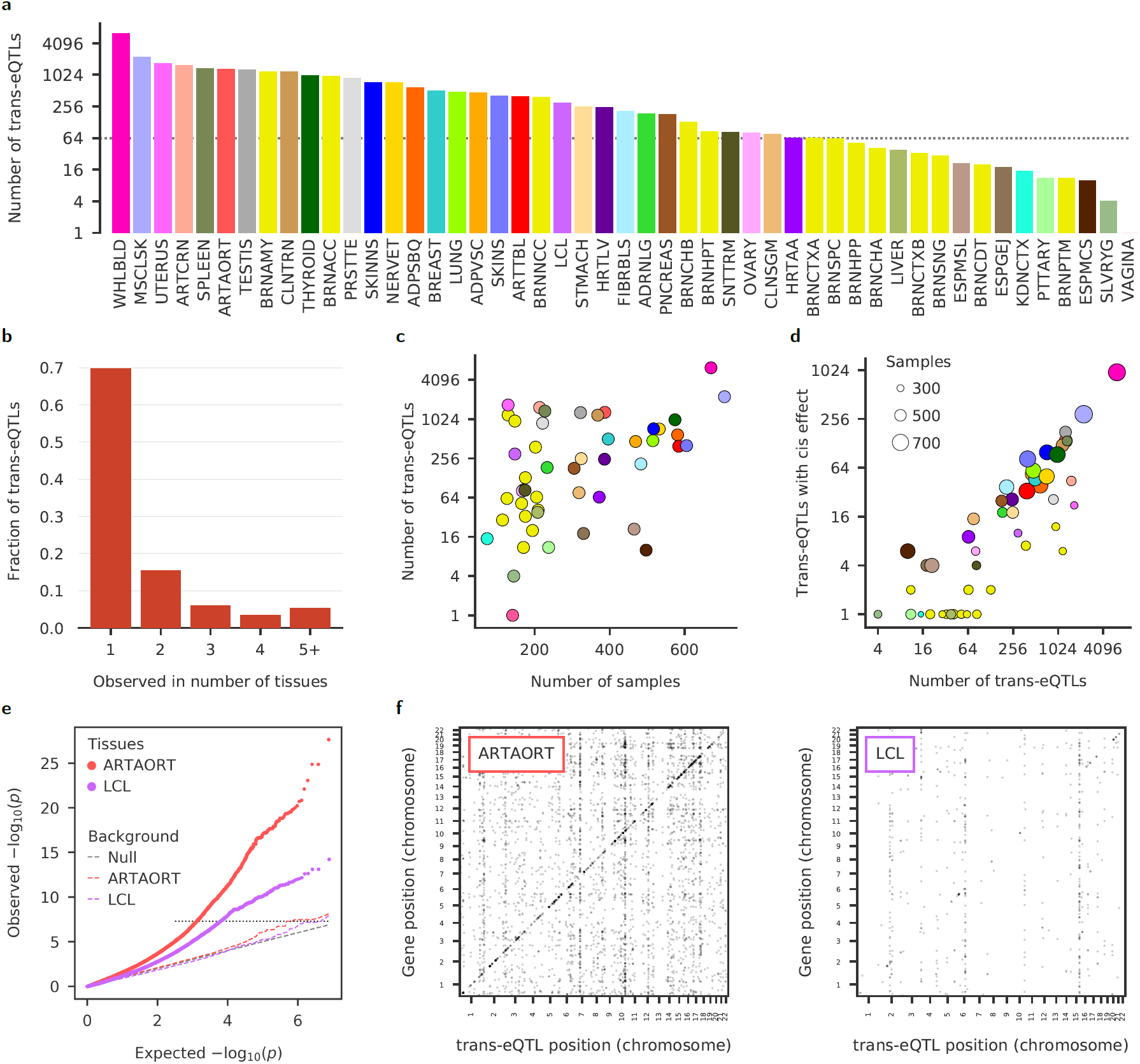
Tejaas identifies many thousands of putative trans-eQTLs in GTEx data. In each of the 49 GTEx tissues, we applied the KNN confounder correction and calculated genome-wide reverse regression *p*-values with Tejaas. Cis genes within ±1Mb of the candidate SNP were excluded from the regression. From the genome-wide significant SNPs (*p* < 5 × 10^−8^) we selected the strongest as lead trans-eQTLs, removing SNPs in strong LD (*r*^2^ ≥ 0.5) with a lead SNP. **a**, Number of lead trans-eQTLs discovered per tissue, on a logarithmic scale. For GTEx tissue abbreviations, see Supplementary Appendix 2. The dotted line indicates the cut-off used for choosing tissues for enrichment analysis. **b**, Proportion of trans-eQTLs discovered in a given number of tissues (excluding brain tissues). 70% of the lead trans-eQTLs are not in strong LD with any lead trans-eQTL from another tissue. **c**, Number of lead trans-eQTLs discovered in a tissue (log scale) versus the number of samples for that tissue. **d**, Trans-eQTLs act via cis-eGenes. Number of lead trans-eQTLs versus the number of discovered lead trans-eQTLs that also happen to be cis-eQTLs in GTEx consortium analysis [4]. **e**, Representative examples of quantile-quantile plots for artery aorta (ARTAORT) and EBV-transformed lymphocytes (LCL) with their negative controls (dashed), obtained by randomly permuting the sample IDs of genotypes. **f**, Representative examples trans-eQTL maps for ARTAORT and LCL, with genomic positions of trans-eQTLs (x-axis) against the genomic positions of their target genes (y-axis). The diagonal band corresponds to cis-eQTLs.

The predicted trans-eQTLs are tissue-specific, with 70% occurring in single tissues (Fig. 3b). The number of trans-eQTLs discovered increases roughly exponentially with the number of samples (Fig. 3c) for *N* > 250, pointing to the importance of sample size to discover more trans-eQTLs. Interestingly, about a quarter of trans-eQTLs predicted in each tissue are also significant cis-eQTLs (Fig. 3d). The effects on the target genes could plausibly be mediated by these cis-eGenes. The quantile-quantile plots for two representative tissues demonstrate the enrichment in significant Tejaas *p*-values, while the negative controls show the expected uniform distribution of *p*-values (Fig. 3e), confirming the correctness of the *p*-values reported by Tejaas. The maps of trans-eQTLs and their target genes (Fig. 3f) illustrate similar patterns as observed earlier in yeast [14].

### Functional enrichment analyses of trans-eQTLs

Given the known difficulties to replicate and validate trans-eQTLs [3, 20] and the lack of RNA-Seq datasets with coverage of tissues other than whole blood, we tested the validity of our results by analyzing the enrichment of the predicted trans-eQTLs in functionally annotated genomic regions. One would expect only true eQTLs to be enriched in these regions. The functional enrichment measurements were compared to a set of randomly chosen SNPs from the GTEx genotypes (Supplementary Sec. 5.6). The trans-eQTLs were discovered excluding all genes in the vicinity of that SNP and therefore it is unlikely to observe functional enrichments driven by falsely discovered cis-eQTLs.

In Fig. 4, we show the functional enrichment of tissues which had more than 64 trans-eQTLs, as indicated by the dotted line in Fig. 3a. This mostly includes non-brain tissues. With low number of trans-eQTLs, enrichment analyses would be statistically unreliable, as for example, observed when comparing all the brain tissues (Supplementary Fig. S16).

**Fig. 4.**
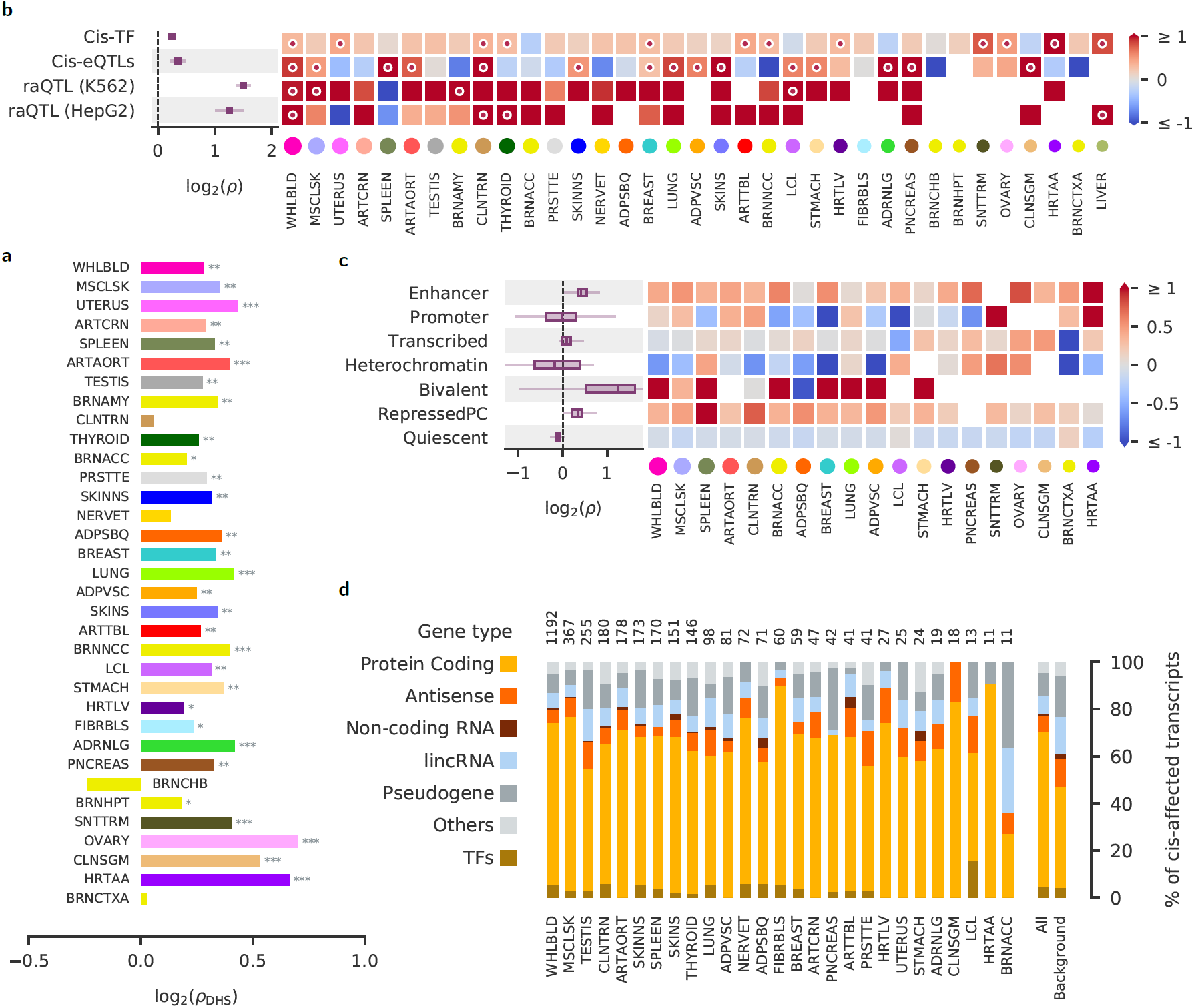
The discovered lead trans-eQTLs are enriched in open chromatin and regulatory regions. Log_2_ enrichments (x-axis) within accessible chromatin regions from [21]. The significance is denoted by * for *p* ≤ 0.05, ** for *p* ≤ 0.01, and *** for *p* ≤ 0.001. The GTEx tissues are ordered by the number of lead trans-eQTLs. For their abbreviations, see Supplementary Appendix 2. **b**, Log_2_ enrichments near known eQTLs and reporter assay QTLs (raQTLs) [22]. Cis-TF: enrichment to occur within ±100 kbp from transcription factors reported in [23]; Cis-eQTL: enrichment among cis-eQTLs SNPs reported in the GTEx v8 analysis [4]; raQTL: enrichment in raQTL regions showing enhancer-like activity in K562 or HepG2 cells [22]. Heatmap colors encode log_2_ enrichment, circular marks signify *p* < 0.01. The area of the colored circles on x-axis labels indicates the log number of discovered lead trans-eQTLs. Left plot: mean log_2_ enrichment across all tissues. **c**, Log_2_ enrichments within tissue-specific regulatory regions. Only tissues that could be matched to the corresponding tissue annotation in the Roadmap Epigenomics Project [24] and had at least 0 trans-eQTLs are shown. Enhancers and bivalently marked regions show clear enrichments for most tissues. **d**, Types of transcripts affected in cis by the lead trans-eQTLs. Only tissues with at least 0 cis-affected transcripts (numbers on top) are shown.

DNase I hypersensitive sites (DHSs) mark accessible regions of the chromatin and could indicate regulatory or biochemical activity, such as promoters, enhancers or actively transcribed regions. Predicted trans-eQTLs occur more often than expected by chance within the DHS regions measured and aggregated across 125 cell and tissue types [21], with significant positive DHS enrichment (*p* ≤ 0.05) in 30 out of 34 tissues and a *p*-value ≤ 0.01 in 26 tissues (Fig. 4a). Using data available in the GTEx Portal, we also found enrichment across a range of annotated regulatory elements such as enhancers and transcription binding sites (Supplementary Fig. S11). The enrichment in open chromatin and annotated regulatory regions suggest that the predicted trans-eQTLs possess regulatory activity more often than expected by chance.

Trans-eQTLs may also act via cis-eQTLs, where the cis-eGene (for example, some known TF) regulates other distant genes. Indeed, we observed a significant enrichment of trans-eQTLs being also cis-eQTLs [4] in the same tissue (Fig. 4b). The cis-eGenes of the novel trans-eQTLs have a higher proportion of protein-coding genes than the background distribution of all GTEx cis-eGenes (orange, Fig. 4d). Although the cis-affected genes are not enriched in TFs (gold, Fig. 4d), the trans-eQTLs are enriched proximal (≤100Kb) to TFs (first line in Fig. 4b).

In Fig. 4b, we show the enrichment of the trans-eQTLs being also reporter assay QTLs (raQTLs) for two cell types, K562 and HepG2 [22]. Reporter assay QTLs (raQTLs) are SNPs that affect the activity of promoter or enhancer elements. K562 is an erythroleukemia cell line with strong similarities to whole blood tissue and HepG2 cells are derived from hepatocellular carcinoma with similarities to liver tissue. The trans-eQTLs from whole blood and liver are strongly enriched (*p* < 0.01), suggesting that at least some trans-eQTLs act via altering the activity of putative regulatory elements in a cell-type-specific manner.

With the high sensitivity to discover trans-eQTL by Tejaas, it becomes possible to describe and disentangle tissue-specific enrichments. Using chromatin state predictions from a set of tissues from the Roadmap Epigenomics project [24], we show that the trans-eQTLs are enriched in enhancer, bivalent and repressed polycomb regions of their matched tissues (Fig. 4c). As expected, they are depleted in the inaccessible heterochromatin regions for most of the tissues.

We checked for possible confounding due to population substructure and cross-mappable genes (by ambiguously mapped reads). Some of the trans-eQTLs have quite different allele frequencies between GTEx subpopulations (Supplementary Fig. S14). After adapting our null background to match the distribution of allele frequency differences (between subpopulations) of the predicted trans-eQTLs, the enrichments in DHS and GWAS are not significantly affected (Supplementary Fig. S15). Saha *et al.* [25] had earlier raised the concern of false trans signals from ambiguously mapped reads. We found similar enrichment in DHS and cis-eQTLs even after masking all possible cross-mappable genes for each tested SNP(Supplementary Fig. S13).

### Association with complex diseases

We investigated the overlap between trans-eQTLs discovered by Tejaas and GWAS variants to search for trans-regulatory mechanisms that affect complex diseases. First, we checked for every tissue, whether more trans-eQTLs overlap with GWAS catalog SNPs [26] than expected by chance. Out of the 28 tissues that have more than 100 lead trans-eQTLs, 27 tissues showed positive enrichment in the GWAS catalog SNPs (Fig. 5a). 21 tissues had an enrichment *p*-value *p* ≤0.05, 20 had *p*≤ 0.01 and 15 had *p* ≤ 0.001. The GWAS catalog SNPs overlapping the trans-eQTLs are associated with a wide range of traits, many of which are not related to complex diseases.

**Fig. 5.**
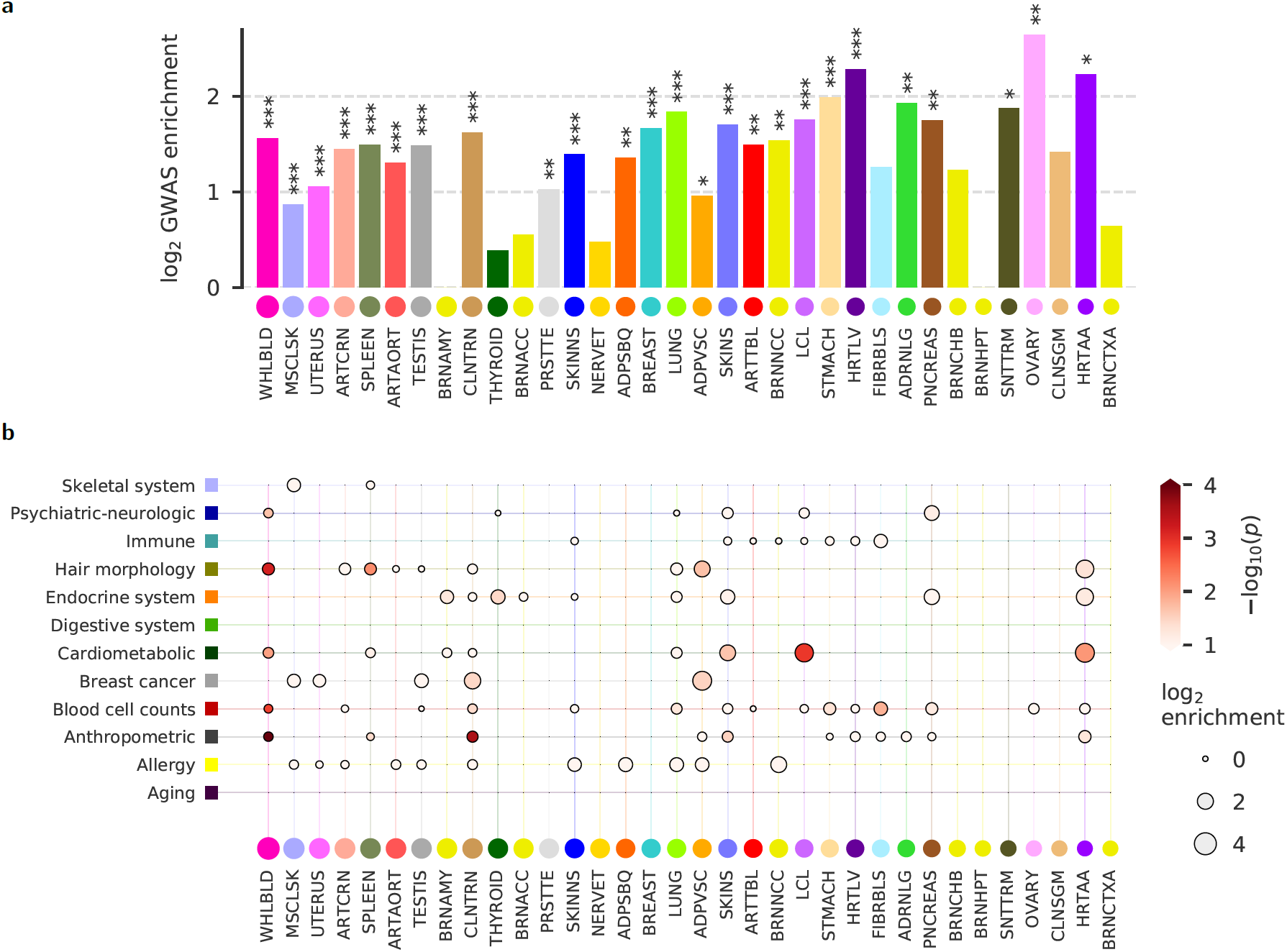
Trans-eQTLs are enriched among GWAS risk SNPs for complex diseases. **a**, Trans-eQTLs are enriched with SNPs from the GWAS Catalog. Significance is denoted by * for *p* 0.05, ** for *p*≤ 0.01, and *** for *p* ≤0.001. **b**, Enrichment of lead trans-eQTLs discovered in GTEx tissues (x-axis) among GWAS SNPs associated with specific disease categories (y-axis). Bubble size indicates log_2_ enrichment, bubble color indicates significance (−log _10_(*p*)). Bubbles are shown for positive enrichment with *p* ≤ 0.1.

To focus on associations with complex diseases, we used the imputed GWAS summary statistics from 87 complex diseases compiled by Barbeira *et al.* [27]. These 87 traits were broadly classified into 12 disease categories. Trans-eQTLs from several tissues are enriched in disease categories that suggest a physiological link (Fig. 5b). Trans-eQTLs in whole blood (WHLBLD), heart atrial appendage (HRTAA), and transformed lymphocytes (LCL) are 1.7-fold, 7-fold, and 6.42-fold enriched in cardiometabolic traits, with *p* = 0.01, *p* = 0.008 and *p* = 0.0012, respectively. Whole blood trans-eQTLs are also 1.3-fold enriched (*p* = 0.0014) in blood related traits, such as variations in different blood cell counts, *e.g.* eosinophil, granulocyte, lymphocyte, monocyte, etc. Trans-eQTLs discovered in the thyroid gland overlap (2.8-fold enriched, *p* = 0.03) with endocrine-associated SNPs. Adipose visceral (ADPVSC) trans-eQTLs are enriched among breast cancer SNPs (7.89-fold, *p* = 0.02). Some associations seem unexpected and could hint at interesting, unknown roles of certain tissues in specific diseases, for instance the overlap of the transverse colon (CLNTRN) trans-eQTLs with anthropometric and breast cancer SNPs, or the nucleus accumbens (BRNACC) with allergies. More insight can be obtained from the disease-specific enrichment for each tissue in Supplementary Fig. S18, such as stomach (STMACH) trans-eQTLs enriched in SNPs associated with Crohn’s disease (13-fold, *p* = 0.01), or heart artery aorta trans-eQTLs enriched in SNPs associated with hypothyroidism (4.94-fold, *p* = 0.06).

To investigate possible implications and mechanisms of the predicted trans-eQTLs that are also GWAS SNPs, we focused on trans-eQTLs found in tissues that are suggestive of a physiological relation to their associated GWAS traits. For each of them, we examined their top 20 target genes.

SNP rs60977503 (chr2:217006659), predicted to be a trans-eQTL in breast tissue, overlaps with a GWAS hit in estrogen receptor-negative breast cancer. Among the top 20 predicted target genes of rs60977503 we found four genes associated with breast cancer. These include FAM183A, which is upregulated in breast cancer cells in response to Notch signaling [28]; MUC4, expressed in 95% of breast carcinomas [29]; HSPB6, which is downregulated in breast cancer [30, 31] and CCL28, which promotes breast cancer proliferation, tumor growth and metastasis [32].

Similarly, SNP rs4538604, predicted as a trans-eQTL in stomach, resides in the inflammatory bowel disease (IBD) 5 locus that has also been associated with Crohn’s disease [33]. Some of its cis-genes have been linked to the disease, such as RAPGEF6, implicated in recovery after mucosal injury [34] and SLC22A5 [35]. Among the top predicted trans target genes of rs4538604 is the receptor for the chemotactic and inflammatory peptide anaphylatoxin C5a (C5AR1). It has been found to be differentially expressed in ulcerative colitis patients [36] and IBD patients that respond to Anti-TNF*α* [37]. The trans-targets RPS21 and ZNF773 are associated with colorectal cancer [38,39], and CDC42SE2 is upregulated in IBD [40]. At least seven other GWAS hits associated with Crohn’s disease overlap with predicted trans-eQTLs, four in small intestine and two more in spleen tissue [41], highlighting the potential relevance of our predictions.

As a third example, rs12040085 is a predicted trans-eQTL in adipose visceral tissue in the 1p33 locus. This region is a GWAS locus related to body mass index (BMI) and body fat percentage. Eight of the top 20 predicted trans target gene of rs12040085 are directly associated with BMI, obesity, and body height. Four of them, CDIN1 (chr15), LINGO1 (chr15), LINC01184 (chr5) and LOC105369911 (chr12), lie within reported GWAS loci related to BMI, body height and obesity and are located on different chromosomes from rs12040085 [42–45]. The target genes TRDMT1, ZNF418, NAT1 and CDC7 have been experimentally associated through their expression levels or through knockouts, or are used as biomarkers, for waist circumference, BMI, obesity or insulin resistance [46–50].

These examples point to the important role that trans-eQTLs could play in complex diseases. It will of course require larger analysis and more automated methods to integrate multiple data sources for finemapping and analyzing all predicted candidates. All our results and scripts used in this study are made public to facilitate further analyses.

## Discussion

Trans-eQTL discovery has come into focus over the past few years, since multiple studies consistently found that 60%–90% of the heritable gene expression variance is contributed by trans-eQTLs. The recently proposed omnigenic model of complex traits highlights the importance of trans-regulated networks in understanding causative disease pathways [2, 51]. According to this model, most of the genetic variance is driven by weak trans effects of peripheral genes on a set of core genes, which in turn affect the risk to develop the disease. However, trans-eQTLs are more difficult to discover than cis-eQTLs due to the extra multiple testing burden and their small effect sizes. Existing methods would require enormous sample sizes – more than one million by some estimates [52] – to reliably identify trans-eQTLs, and it will take years to develop such resources.

Here, we proposed an unconventional approach that reverses the regression direction to predict trans-eQTLs with small effects on the expression of multiple targeted genes by aggregating their explanatory signal while being unaffected by expression correlations. We created a fast, parallel open-source software and showed its power using semi-synthetic data. With its combination of reverse regression and KNN correction, Tejaas is more powerful than other existing methods to predict trans-eQTLs. We then applied Tejaas on the GTEx dataset and predicted thousands of trans-eQTLs at genome-wide significance. To our knowledge, these results represent the first systematic large-scale prediction of trans-eQTLs in the GTEx dataset. Simple regression of SNP-gene pairs could not have predicted those trans-eQTLs because of their low effect sizes. Forward regression, on the other hand, is impeded by the strong correlated noise of the gene expression levels [15].

The large number of predicted trans-eQTLs allowed us to obtain statistically significant enrichments for them in regions characterized as functional or regulatory according to various independent experimental genome-wide procedures. So far, most studies have predicted too few trans-eQTLs for such an analysis. Other studies are large-scale meta-analysis projects whose inherent selection biases did not allow for enrichment analyses. For example, the meta-analysis of 31 684 individuals on whole blood by the eQTLGen consortium [3], which predicted 3 853 trans-eQTLs, tested only GWAS-associated SNPs for trans-effects. Consequently, the discovered trans-eQTLs inherited the enrichments of the GWAS SNPs.

One major source of false trans-eQTL predictions could be population substructure. False associations between SNPs and gene expression levels can arise if both of them are influenced by subpopulation membership, for example via life style or via epistatic effects with the genetic background. We would expect such false positive trans-eQTLs to show up in several tissues. The observation that 70% of the predicted trans-eQTLs are tissue-specific and only ∼5% are found simultaneously in 5 or more tissues (Fig. 3b) indicates that false positives do not make up a large part of our predictions. Some of the trans-eQTLs have quite different allele frequencies between populations, but subsequent analyses using matched null background showed significant DHS enrichment and GWAS enrichment (Supplementary Fig. S14). This suggests weak if any confounding by population substructure in our approach.

The new KNN correction is a simple but efficient method for removing confounders. It can correct out non-linear confounding effects, therefore it should work even if those effects are not well approximated by linear, additive models. It also does not require the confounders to be known. For future eQTL pipelines, it could prove to be very useful when applied after correcting the known confounders with linear methods.

There are several limitations to our method. First, reverse regression cannot identify the target genes of a discovered trans-eQTL, because the *L*_2_ regularization does not encourage sparsity and therefore is not well suited for selecting the informative covariates. Second, the standard deviation *γ* of the normal prior is not learnt from the data, but is set empirically. As expected, a high value of *γ* (> 0.2) could lead to overfitting, whereas a low value of (e.g. *γ* < 0.001) can severely reduce the sensitivity to discover trans-eQTLs. Third, the input gene expression cannot be corrected for confounders using the standard approach of regressing the known confounders or hidden PEER factors [53] (Supplementary Sec. 3.1). Fourth, Tejaas is not designed to pick up strong, single SNP-gene associations. All trans-eQTLs identified to date, including the meta-analysis on whole blood with 31 684 individuals [3], were discovered by strong effects on a single, distant gene. Hence, by design, Tejaas might not replicate these existing trans-eQTLs with statistical significance, although we did find significant replication in whole blood (Supplementary Appendix 3). We therefore expect Tejaas and existing methods to be quite complementary.

In the future we plan to improve Tejaas by encouraging sparsity in the regression coefficients, because we expect only a small fraction of the ∼20 000 genes to be targets of a typical trans-eQTL. One widely adopted Bayesian approach is to use a sparsity-enforcing prior such as a spike-and-slab prior for the effect sizes, which has been previously used with success in other contexts such as fine-mapping in GWAS [54, 55]. Using such prior will improve trans-eQTL discovery, remove the dependency on *γ*, and enable more accurate selection of trans-eQTL target genes.

Robust identification of trans-eQTLs will help us to dissect the interplay between genetic variation, expression levels of genes and the risk for complex diseases. We will need to further increase the number of samples in eQTL datasets. In addition, we need statistical methods with high sensitivity and accuracy to discover trans-eQTLs. Tejaas represents a major step towards this goal and predicts about two orders of magnitude more trans-eQTLs on the GTEx v8 dataset than the state of the art at < 5% false discovery rate. We hope that Tejaas will help to realize the tremendous value of the RNA-seq eQTL datasets that are already available or in production.

## Methods

### Forward Regression

For each SNP, we calculated the *p*-values of association with all the *G* genes independently. Under the null hypothesis that the SNP is not a trans-eQTL, these *p*-values will be independent and identically distributed (iid) with a uniform probability density function,

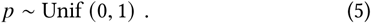

We sort the *p*-values in increasing order; the *k* ^th^ smallest value is called the *k* ^th^ order statistic and is denoted as *p* _(*k*)_ Then *p* _(*k*)_ will be a Beta-distributed random variable,

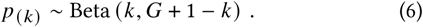

and the expectation of ln (*p*_(*k*)_) will be

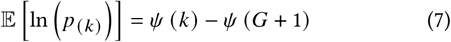

where *ψ* denotes the digamma function. If the candidate SNP is a trans-eQTL and there is an enrichment of *p*-values near zero, then the cumulative sum of (𝔼 [ln (*p* _(*k*)_)] − ln (*p* _(*k*)_)) over *k* will increase monotonically, pass through a maximum and then decrease to an asymptotic value of zero. Hence, we defined the FR-score as,

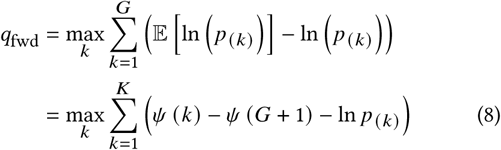

It would be sufficient to calculate the *q*_fwd_ from only the first *K* genes because the rest will not contribute to the low *p*-values. We obtained an empirical null distribution for *q*_fwd_ by permuting the columns of the real genotype matrix – thereby removing any association with the gene expression but retaining the correlation between the gene expression levels. For each SNP, we calculated the *p*-value for *q*_fwd_ from this empirical null.

### Reverse regression

Let x be the genotype vector for a candidate SNP and Y be the *G* × *N* matrix of gene expression levels for *G* genes and *N* samples. Both x and Y are centered and normalized. We model x with a univariate normal distribution whose mean depends linearly on the gene expression

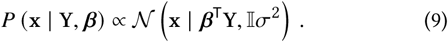

where *β* is the vector of regression coefficients. and *σ*^2^ is the variance of the candidate SNP. The number of samples *N* will usually be on the order of a hundred to a few thousand, much smaller than the number of explanatory variables *G* ≈ 20 000. Therefore, simple maximization of the likelihood would lead to overtrained *β*. Hence we define a normal prior on *β*,

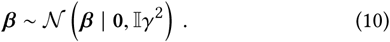

Let ℋ _1_ be the trans-eQTL model which allows *β* ≠ 0 and ℋ_0_ be the null model for which *β* = 0. According to Bayes’ theorem,

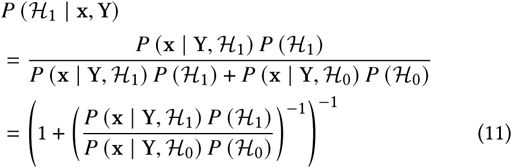

The probability for the model ℋ_1_ is a monotonically increasing function of the likelihood ratio,

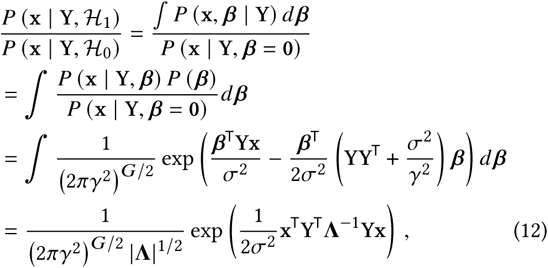

where we have defined Λ := YY^T^ + (*σ* ^2^ *γ*^2^) 𝕀_*G*_. The integration was done using the technique of quadratic complementation. Motivated by Eq. 12, we defined our test statistic RR-score, denoted *q*_rev_, as

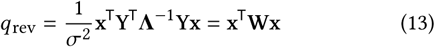

where

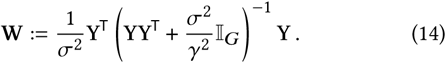

### Null model

Given *q*_rev_ for the candidate SNP, we would like to know how significant this score is. We obtain the null model 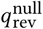 by permuting the elements of x. The distribution of 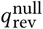 will be different for every candidate SNP depending on their minor allele frequency (MAF) and the variance of the genotype (*σ*^2^). We derived analytical expressions for the expectation value 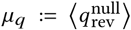 and variance 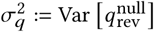 under the permutation null model for any symmetric matrix W and any centered vector x (see Supplementary Text, Appendix 1). Our analytical calculations of *µ*_*q*_ and *σ*_*q*_ match those obtained from the empirical permutation of x (Supplementary Fig. S1). We approximate 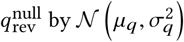.

Finally, the *p*-value of *q*_rev_ for the candidate SNP is

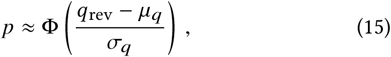

where Φ(*z*) denotes the cumulative normal distribution for a random variable *z*.

### KNN correction

Gene expression measurements are notorious for being dominated by strong confounding effects and the subtle effects of trans-eQTLs are at risk of being drowned out by these strong systematic noise. For the KNN correction, we assume that confounding effects dominate the gene expression. If the samples are close to one another in the expression space, we expect them to be affected by the same confounders. Let y_*n*_ and x_*n*_ be the vectors of expression levels and genotypes respectively for the *n* ^th^ sample. The contribution of confounding effects on y_*n*_ can be corrected by removing the average expression among the *K* nearest neighbors of that sample:

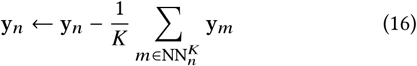

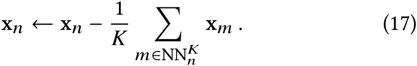

The nearest neighbors 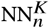 is calculated from the euclidean distances between the samples in a reduced dimension gene expression space. We also remove genotype confounders (such as population substructure) which might lead to similar gene expressions. KNN was shown to be a useful approach for many learning tasks, and since its naive form has a single parameter (*K*), overfitting does not typically occur [56, 57]. The choice of *K* should be such that it captures the locally varying effects of the confounders. A very small value of *K* would not be able to render the statistical noise, while a very large value of *K* will start removing long-range trans-effects (Supplementary Fig. S6). KNN correction does not require the knowledge of known covariates, it is unsupervised and non-linear. Since KNN does not reduce the rank of the gene expression matrix, it works well with Tejaas.

### Simulation method

Simulated data consisted of genotype and gene expression for 450 individuals. After pre-filtering of the GTEx genotype, we randomly sampled 12 639 SNPs. We randomly selected 800 SNPs to be cis-eQTLs. From these cis-eQTLs, we selected a subset 30 SNPs to be trans-eQTLs. We simulated the gene expression data for 12 639 genes, containing non-genetic signals (background noise and confounding factors) and genetic signals (*cis* and *trans* effects) following the strategy of Hore *et al.* [6]. Each gene contained only one SNP, equivalent to assuming that there is at most one cis-eQTL per gene. Hore *et al.*used heteroscedastic background noise, but we created a correlated Gaussian noise with a covariance matrix obtained from the gene expressions in the artery aorta tissue of GTEx. We used the first three principal components of the genotype along with 7 other hypothetical covariates to generate the confounding effects. Each confounding factor was assumed to be affecting a set of randomly chosen 6 320 genes with effect sizes sampled from 𝒩 (0, 1). The strength of ciseffects were sampled from Gamma (4, 0.1) and the direction was chosen randomly. For the trans-eQTLs, the strength of cis-effect was constant (0.6). Additive combination of the noise, the effect of confounding factors and the effect of cis-eQTLs gives a temporary gene expression matrix, on top of which the effects of trans-eQTLs were added. The cis target gene of the trans-eQTLs is considered a transcription factor (TF), which regulated multiple target genes downstream. This ensured that the trans-eQTLs were indirectly associated with the target genes with practically low effect sizes. The effect sizes of the TF on the target genes were sampled from Gamma (*ψ*^trans^, 0.02). We performed simulations with 50, 100 and 150 target genes and sampled the effect sizes of the TFs on the target genes according to a Gamma distribution with mean effect size between 0.1 and 0.4. See Supplementary Sec. 3 for further details.

### GTEx data and quality control

We analyzed 49 tissues with ≥70 samples with available genotype and expression measurements from the GTEx v8 project. We downloaded the genotype files and phased RNA-seq read count expression matrix. The obtained genotype was quality filtered by the GTEx consortium [4]. Genotype was split in chromosomes, variants with missing values were filtered out and sex chromosomes were removed. 8 048 655 variants with minor allele frequency (MAF) ≥ 0.01 were retained for further analysis. We calculated TPMs (Transcripts Per Million) from the phASER expression matrix. We retained genes with expression values > 0.1 and more than 6 mapped reads in at least 20% of the samples.

For finding target genes of the trans-eQTLs, we needed the explicit covariate-corrected gene expression. We downloaded the covariate files from the GTEx portal [58] and used the first 5 principal components of the genotype, donor sex, WGS sequencing platform (HiSeq 2000 or HiSeq X) and WGS library construction protocol (PCR-based or PCR-free). Additionally, from phenotype files available in dbGaP, we included donor age and post mortem interval in minutes (‘TRISCHD’) as covariates. We inverse normal transformed the TPMs and used CCLM to remove the effect of covariates.

### LD pruning

We calculated LD between variants with PLINK using an *r*^2^ > 0.5 within an 200kbp sliding window. We pruned the list of trans-eQTLs by retaining only those lowest *p*-values in each independent LD regions.

### Functional enrichment

For every functional annotation, we sampled 5000 random SNPs from the GTEx genotype. The fraction of random annotated SNPs averaged over 50 replicates gives the background frequency. The fraction of annotated trans-eQTLs divided by the background frequency gives the annotation enrichment. We used a binomial test to calculate the *p*-values for the enrichment *ρ*. If *T* is the number of trans-eQTLs in the tissue, then the probability of finding *k* annotated trans-eQTLs is,

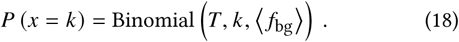

where ⟨*f*_bg_ ⟩ is the background frequency and *P* (*x* > *k*) gives us the *p*-value for the tissue-GWAS pair. See also Supplementary Sec. 5.6.

### GWAS data

We obtained GWAS summary statistics for 87 complex traits compiled by Barbeira *et al.* [27]. These studies were imputed and harmonized to GTEx v8 variants with MAF ≥ 0.01 in European samples.

### GWAS enrichment

For every GWAS, we sampled 5000 random SNPs from the GTEx genotype. The fraction of random SNPs that overlap with the GWAS averaged over 300 replicates gives the background frequency. The fraction of trans-eQTLs that overlap with the GWAS divided by the background frequency gives the GWAS enrichment. We calculated the *p*-values for enrichment in the same way as functional enrichment. For category-wise enrichment, we checked the overlap of trans-eQTLs with all disease-associated SNPs in that category. For global enrichment, we checked the overlap of trans-eQTLs with all disease-associated SNPs in the dataset. For the 87 GWAS traits, all SNPs with *p* < 10^−7^ were considered to be a significant GWAS hit. See also Supplementary Sec. 6.2.

## Data availability

This study analyzed data from the GTEx project, which are publicly available by application from dbGap (Study Accession phs000424.v8.p2). The results for the GTEx Analysis v8 were downloaded from the GTEx portal (https://gtexportal.org). The GWAS catalog was downloaded from https://www.ebi.ac.uk/gwas/home, and the GWAS summary statistics from 87 traits harmonized and imputed to GTEx v8 variants are available at https://doi.org/10.5281/zenodo.3657902. We have publicly released the trans-eQTLs discovered by applying our Tejaas method to GTEx data; the summary association statistics for 49 tissues are available at http://www.user.gwdg.de/~compbiol/tejaas/2020_03. Reporter Assay QTLs were obtained from https://sure.nki.nl/. DHS annotations were obtained from [21] https://resources.altius.org/publications/Nature_Thurman_et_al/. Tissue-matched regulatory elements were downloaded from the Roadmap Epigenomics Project https://egg2.wustl.edu/roadmap/web_portal/chr_state_learning.html. GENCODE annotations v26 downloaded from https://www.gencodegenes.org/human/release_26.html Transcription Factors dataset was obtained from [23] http://humantfs.ccbr.utoronto.ca/download.php

## Code Availability

Tejaas is open-source code released under the GNU General Public License version 3. It is available at https://github.com/soedinglab/tejaas.

The code used for simulations is available at https://github.com/banskt/trans-eqtl-simulation. The code used for GTEx analyses is available at https://github.com/banskt/trans-eqtl-pipeline. Other software used: MatrixEQTL [16], downloaded from http://www.bios.unc.edu/research/genomic_software/Matrix_eQTL; PLINK [59], downloaded from https://www.cog-genomics.org/plink/2.0/; LDstore [60], downloaded from http://www.christianbenner.com/; VCFTools [61], downloaded from http://vcftools.sourceforge.net/.

## Acknowledgments

This work was supported by the German Federal Ministry of Education and Research (BMBF) within the framework of the e:Med research and funding concept (grants e:AtheroSysMed 01ZX1313A-2014) and by the German Research Foundation (DFG) under Germany’s Excellence Strategy - EXC 2067/1-390729940. RM and RN were supported by a DAAD fellowship. We thank Markus Scholz for helpful communications and alerting us to the problem of strong cis-eQTLs, which led us to use cis-masking. We are grateful to Hae Kyung Im and Alvaro Barbeira for email communications and kindly providing us the GWAS summary statistics from 87 traits harmonized and imputed to GTEx v8 variants. We thank our colleagues, especially Eli Levy Karin, Christian Roth, Wanwan Ge, Salma Sohrabi-Jahromi, Milot Mirdita and Ruoshi Zhang for discussions and feedback. We used the data generated by the GTEx Consortium for the trans-eQTL analysis and thank the participants of the GTEx Consortium and the research staff who worked on the data collection.

## Author Contributions

S.B. and F.L.S. wrote the software with assistance from A.K. and R.M. S.B. designed and performed the simulations. F.L.S. performed the GTEx preprocessing. F.L.S. and S.B. analyzed the GTEx data F.L.S. checked the contribution of known covariates in KNN and the effect of cross-mappable genes. K.E.D. analyzed the GWAS data. R.N. contributed to the initial phase of the project.S.B. drafted the manuscript and supplementary with assistance from F.L.S.; J.S., S.B. and F.L.S. reviewed and edited the draft. J.S. designed the reverse regression with input from S.B., F.L.S., A.K. and R.M., acquired funding, and supervised the project.

## Additional Information

### Supplementary Text and Figures

Supporting information available for download.

