## Supplemental Text, Figures and Tables for "Reverse regression increases power for detecting trans-eQTLs"

### Supplementary Text and Figures

**Appendix 2. Abbreviation of GTEx tissues** 37

**Appendix 3. eQTLGen Replication** 38

**References** 39

### 1 Forward regression

A trans-eQTL is generally expected to influence the expression levels of tens to hundreds of genes and we take advantage of this signature to increase the sensitivity to detect them. Brynedal *et al.* proposed a method called cross-phenotype meta-analysis (CPMA) to analyze the $p$-value distribution of pairwise associations of a candidate SNP with all measured genes [1]. We follow the same idea but use a different statistic for forward regression (FR) to find the trans-eQTLs.

#### 1.1 Notation

We use bold capital cases for matrices. $\mathbf{X}$ is the $I \times N$ matrix of genotypes for $I$ SNPs and $N$ samples. $\mathbf{Y}$ is the $G \times N$ matrix of gene expression levels for $G$ genes and $N$ samples. For the FR analysis, both $\mathbf{X}$ and $\mathbf{Y}$ are centered and normalized. We use bold small cases for vectors. The rows of $\mathbf{X}$ and $\mathbf{Y}$ are denoted as $\mathbf{x}_i$ and $\mathbf{y}_g$ respectively for every SNP $i \in \{1, \dots, I\}$ and for every gene $g \in \{1, \dots, G\}$. The columns of $\mathbf{X}$ and $\mathbf{Y}$ are denoted as $\mathbf{x}_n$ and $\mathbf{y}_n$ respectively for every sample $n \in \{1, \dots, N\}$. Both $\mathbf{x}_i$ and $\mathbf{y}_g$ are vectors of size $N$, while $\mathbf{x}_n$ and $\mathbf{y}_n$ are of sizes $I$ and $G$ respectively.

#### 1.2 FR-score

For each SNP $\mathbf{x}_i$, we calculated the $p$-values of association with $\mathbf{y}_g$ for all the $g \in \{1, \dots, G\}$ genes independently. Under the null hypothesis that the SNP is not a trans-eQTL, these $p$-values will be independent and identically distributed (iid) with a uniform probability density function,

$$p \sim \text{Unif}(0, 1) \; . \tag{1}$$

If SNP $i$ is a trans-eQTL, then more genes than expected by chance will have low $p$-values for association with the SNP, leading to a higher density of $p$-values near zero. We defined the FR-score ($q_{\text{fwd}}$) as a statistic that estimates the difference between the observed $p$-value distribution from the data and the uniform distribution expected by chance.

We sort the $p$-values in increasing order and the $k^{\text{th}}$ smallest value is called the $k^{\text{th}}$ order statistic, and is denoted as $p_{(k)}$. If $p$ is uniformly distributed ($p \in (0, 1)$), then $p_{(k)}$ will be a Beta-distributed random variable,

$$p_{(k)} \sim \text{Beta}(k, G + 1 - k) \; . \tag{2}$$

For any random variable $z \sim \text{Beta}(\alpha, \beta)$,

$$\begin{aligned}\mathbb{E}[z] &= \frac{\alpha}{\alpha+\beta} \\ \mathbb{E}[\ln z] &= \psi(\alpha) - \psi(\alpha+\beta) \ ,\end{aligned} \tag{3}$$

where $\psi$ denotes the digamma function. Using the identities in Eq. 3 and noting that $\alpha = k$ and $\beta = G + 1 - k$, we obtained the expectation of $\ln p_{(k)}$ as,

$$\mathbb{E}\left[\ln p_{(k)}\right] = \psi(k) - \psi(G+1) \tag{4}$$

If the candidate SNP is a trans-eQTL and there is an enrichment of $p$-values near zero, then the observed values of $\ln p_{(k)}$ will be lower than that expected by chance when $k$ is small. Therefore, the cumulative sum of $\left(\mathbb{E}\left[\ln p_{(k)}\right] - \ln p_{(k)}\right)$ over $k$ will increase monotonically, pass through a maximum and then decrease to an asymptotic value of zero. Hence, we defined the FR-score as,

$$q_{\text{fwd}} = \max_{k} \sum_{k=1}^{G} \left(\mathbb{E}\left[\ln p_{(k)}\right] - \ln p_{(k)}\right) \tag{5}$$

Intuitively, if the candidate SNP is not a trans-eQTL, $\sum_{k=1}^{G} \left(\mathbb{E}\left[\ln p_{(k)}\right] - \ln p_{(k)}\right)$ will fluctuate around zero. Hence, $q_{\text{fwd}}$ will remain close to 0 for these SNPs. However, a trans-eQTL will be associated with many genes and the FR-score will be high ($q_{\text{fwd}} \gg 0$) because there will be many genes with a lower $p$-value than expected by chance. Of course, there will be other genes which are not associated with the trans-eQTL and the $p$-values for association with these genes will not contribute to the $q_{\text{fwd}}$. Therefore, it would be sufficient to calculate the $q_{\text{fwd}}$ from the first $K$ genes with lowest $p$-values, instead of all $G$ genes:

$$\begin{aligned}q_{\text{fwd}} &= \max_{k} \sum_{k=1}^{K} \left(\mathbb{E}\left[\ln p_{(k)}\right] - \ln p_{(k)}\right) \\ &= \max_{k} \sum_{k=1}^{K} \left(\psi(k) - \psi(G+1) - \ln p_{(k)}\right)\end{aligned} \tag{6}$$

#### 1.3 Null model and *p*-values for FR-score

We need to define a null distribution to evaluate the significance of any observed $q_{\text{fwd}}$. Let $q_{\text{fwd}}^{\text{null}}$ be an iid sample from the null distribution. Since it is analytically intractable to define a probability density of $q_{\text{fwd}}^{\text{null}}$, we define an empirical null distribution. The empirical cumulative distribution function (ECDF) of $q_{\text{fwd}}^{\text{null}}$ can be used to ascertain significance of any observed $q_{\text{fwd}}$ to obtain a $p$-value, denoted as $p_{q_{\text{fwd}}}$,

$$p_{q_{\text{fwd}}} = 1 - \text{ECDF}\left(q_{\text{fwd}}\right) \tag{7}$$

One possible way to obtain a null model is by calculating $q_{\text{fwd}}^{\text{null}}$ for a large number of *null* SNPs assuming they are not trans-eQTLs. Following our hypothesis in Eq. (1), the $p$-values for association of each of these *null* SNPs with the genes can be sampled each time from $\text{Unif}(0,1)$ distribution. However, the iid assumption of $p$-values breaks down in real data because the gene expressions are strongly correlated. Therefore, the $p$-values for association between the SNP and the genes are strongly correlated and cannot be sampled from the $\text{Unif}(0,1)$ distribution.

To account for the correlation, we randomized the sample labels of $\mathbf{X}$ by permuting the columns of the real genotype matrix – thereby removing any association with the gene expression $\mathbf{Y}$ but retaining the correlation between the gene expression levels. We then calculated $q_{\text{fwd}}^{\text{null}}$ for all SNPs using the ‘permuted’ genotype and the gene expression. We used this empirical null distribution to calculate $p_{q_{\text{fwd}}}$ from Eq. 7 .

The estimate of $p_{q_{\text{fwd}}}$ gets increasingly noisy with higher values of $q_{\text{fwd}}$ (low $p$-values) because of lack of sampling points in that region. In that regime it is better to rely on a parametric fit of the exponentially falling part of the cumulative distribution. We sorted the $q_{\text{fwd}}^{\text{null}}$ in an increasing order. We set the limit beyond which we will use the exponential extrapolation at $n^{\text{top}} = \min\{100, 0.1I\}$, where $I$ is the total number of SNPs being used for the calibration of the null model. This ensures that the extrapolation is applied at most to the top 10% of points, and, if more data are available, only to the top 100. Let us denote the $q_{\text{fwd}}^{\text{null}}$ at this cut-off point as $q'$. For any observed $q_{\text{fwd}} \geq q'$, we use a maximum-likelihood estimate based on the $n^{\text{top}}$ data points and is given by,

$$p_{q_{\text{fwd}}} = \frac{n_{\text{top}}}{I} \exp\left(-\frac{q_{\text{fwd}} - q'}{\lambda}\right) , \tag{8}$$

with

$$\lambda := \frac{1}{n_{\text{top}}} \sum_{j=1}^{n_{\text{top}}} \left(q_{\text{jpa},j}^{\text{null}} - q'\right) . \tag{9}$$

where $q_{\text{jpa},j}^{\text{null}}$ is the $j^{\text{th}}$ value in the ordered sequence of $q_{\text{fwd}}^{\text{null}}$. Note that for $q_{\text{fwd}} = q'$ this equation yields $p_{q'} = n_{\text{top}}/I$, the same as the empirical value.

##### 1.4 Comparison with CPMA

Brynedal *et al.*used a similar idea for finding trans-eQTLs by testing each SNP for enrichment of association $-\ln(p)$ values across all genes with a null hypothesis that $-\ln(p)$ should be exponentially distributed, with a decay parameter $\lambda = 1$. Under the joint alternative hypothesis, a subset of association statistics are non null and $\lambda \neq 1$. They compared the evidence for these hypotheses as a likelihood ratio test for CPMA, with the statistic $S_{\text{CPMA}}$ defined as,

$$S_{\text{CPMA}} = -2 \ln\left(\frac{P\,(\text{data} \mid \lambda = 1)}{P\left(\text{data} \mid \lambda = \hat{\lambda}\right)}\right) \tag{10}$$

where $\hat{\lambda}$ is the observed exponential parameter from the data. They accounted for the extensive correlation among the gene expression levels by testing the significance empirically using a simulated gene expression matrix with the same covariance as the real gene expression. To define high-confidence trans-eQTLs, they combined empirical CPMA statistic from multiple data sets using sample size weighted meta-analysis.

FR measures the significance of the enrichment in the $p$-value distribution near zero, and CPMA measures the significance of the enrichment in the $-\ln(p)$ distribution. Theoretically, both should give similar ranking of SNPs for being trans-eQTLs. Since there are no currently available software for CPMA, we implemented FR-score in Tejaas and compared the performance with RR-score.

#### 2 Reverse regression

In this section, we discuss the reverse regression (RR) model and introduce the RR-score, denoted as $q_{\text{rev}}$ for finding the trans-eQTLs. We also describe a null model and explain how to obtain significance $p$-values for the $q_{\text{rev}}$ of a candidate SNP, denoted as $p_{q_{\text{rev}}}$. We also discuss the numerical implementation of RR-score in Tejaas and several other options used in Tejaas.

#### 2.1 Notation

We use the same notations from Sec. 1.1. The genotype vector $\mathbf{x}_i$ for every $i^{\text{th}}$ SNP is centered but not normalized to allow for null model calculation. $\mathbf{Y}$ is centered and normalized.

#### 2.2 Model description

We model $\mathbf{x}_i$ with a univariate normal distribution whose mean depends linearly on the gene expression through a column vector of regression coefficients $\boldsymbol{\beta}_i$ ($\in \mathbb{R}^G$),

$$P(\mathbf{x}_i \mid \mathrm{Y}, \boldsymbol{\beta}_i) \propto \mathcal{N}\left(\mathbf{x}_i \mid \boldsymbol{\beta}_i^{\mathsf{T}} \mathrm{Y}, \sigma_i^2 \mathbb{I}_N\right) . \tag{11}$$

where the variance of the $i^{\text{th}}$ SNP is given as $\sigma_i^2$. For the rest of the manuscript, we drop the subscript $i$ for ease of reading although we note that all calculations are done for the $i^{\text{th}}$ SNP. The log likelihood of this regression task is

$$\begin{aligned} \ln \mathcal{L}(\boldsymbol{\beta}) &= \ln P(\mathbf{x} \mid \mathrm{Y}, \boldsymbol{\beta}) \\ &= -\frac{1}{2\sigma^2} \sum_{n=1}^{N} \left(x_n - \boldsymbol{\beta}^{\mathsf{T}} \mathbf{y}_n\right)^2 + \text{const}, \end{aligned} \tag{12}$$

where $x_n$ is the minor allele count of the $i^{\text{th}}$ SNP in $n^{\text{th}}$ sample. The number of samples $N$ will usually be on the order of a hundred to a few thousand, much smaller than the number of explanatory variables $G \approx 20\,000$. Therefore, simple maximization of the likelihood would lead to a dramatically overtrained $\boldsymbol{\beta}$ which would perfectly predict $\mathbf{x}$ on the training data but which would achieve very bad performance on unseen data. The solution is to define a prior on $\boldsymbol{\beta}$. We assume that every $g^{\text{th}}$ element ($g \in \{1, \ldots, G\}$) of $\boldsymbol{\beta}$ is sampled from a normal distribution with variance $\gamma^2$,

$$\beta_g \sim \mathcal{N}\left(\beta_g \mid 0, \gamma^2\right) \tag{13}$$

and maximize the log posterior probability,

$$\begin{aligned} \ln P(\boldsymbol{\beta} \mid \mathbf{x}, \mathrm{Y}) &= \ln \frac{P(\mathbf{x} \mid \mathrm{Y}, \boldsymbol{\beta})\, P(\boldsymbol{\beta})}{P(\mathbf{x} \mid \mathrm{Y})} \\ &= -\frac{1}{2\sigma^2} \sum_{n=1}^{N} \left(x_n - \boldsymbol{\beta}^{\mathsf{T}} \mathbf{y}_n\right)^2 - \frac{1}{2\gamma^2} \sum_{g=1}^{G} \beta_g^2 + \text{const}. \end{aligned} \tag{14}$$

#### 2.3 Bayesian probability

Let $\mathcal{H}_1$ be the trans-eQTL model which allows $\boldsymbol{\beta} \neq \mathbf{0}$ and $\mathcal{H}_0$ be the null model for which $\boldsymbol{\beta} = \mathbf{0}$. According to Bayes' theorem,

$$\begin{aligned} P(\mathcal{H}_1 \mid \mathbf{x}, \mathrm{Y}) &= \frac{P(\mathbf{x} \mid \mathrm{Y}, \mathcal{H}_1)\, P(\mathcal{H}_1)}{P(\mathbf{x} \mid \mathrm{Y}, \mathcal{H}_1)\, P(\mathcal{H}_1) + P(\mathbf{x} \mid \mathrm{Y}, \mathcal{H}_0)\, P(\mathcal{H}_0)} \\ &= \left(1 + \left(\frac{P(\mathbf{x} \mid \mathrm{Y}, \mathcal{H}_1)\, P(\mathcal{H}_1)}{P(\mathbf{x} \mid \mathrm{Y}, \mathcal{H}_0)\, P(\mathcal{H}_0)}\right)^{-1}\right)^{-1} \end{aligned} \tag{15}$$

The probability for the model $\mathcal{H}_1$ is a monotonically increasing function of the likelihood ratio,

$$\frac{P(\mathbf{x} \mid \mathrm{Y}, \mathcal{H}_1)}{P(\mathbf{x} \mid \mathrm{Y}, \mathcal{H}_0)} = \frac{\int P(\mathbf{x}, \boldsymbol{\beta} \mid \mathrm{Y})\, d\boldsymbol{\beta}}{P(\mathbf{x} \mid \mathrm{Y}, \boldsymbol{\beta} = \mathbf{0})}$$

$$
\begin{aligned}
&= \int \frac{P(\mathbf{x} \mid \mathrm{Y}, \boldsymbol{\beta})\, P(\boldsymbol{\beta})}{P(\mathbf{x} \mid \mathrm{Y}, \boldsymbol{\beta} = \mathbf{0})} d\boldsymbol{\beta} \\
&= \int \frac{1}{(2\pi\gamma^2)^{G/2}} \exp\left(\frac{1}{\sigma^2}\boldsymbol{\beta}^{\mathsf{T}}\mathrm{Y}\mathbf{x} - \frac{1}{2\sigma^2}\boldsymbol{\beta}^{\mathsf{T}}\left(\mathrm{Y}\mathrm{Y}^{\mathsf{T}} + \frac{\sigma^2}{\gamma^2}\right)\boldsymbol{\beta}\right) d\boldsymbol{\beta}\,, \qquad (16)
\end{aligned}
$$

where the second line is obtained by using the model defined in Eq. 11 and the prior for $\boldsymbol{\beta}$ defined in Eq. 13. We then defined $\boldsymbol{\Lambda} := \mathrm{Y}\mathrm{Y}^{\mathsf{T}} + (\sigma^2/\gamma^2)\, \mathbb{I}_G$ and used the technique of quadratic complementation to obtain,

$$
\begin{aligned}
&\frac{P(\mathbf{x} \mid \mathrm{Y}, \mathcal{H}_1)}{P(\mathbf{x} \mid \mathrm{Y}, \mathcal{H}_0)} \\
&\quad = \int \frac{1}{(2\pi\gamma^2)^{G/2}} \exp\left(-\frac{1}{2\sigma^2}\left(\boldsymbol{\beta} - \boldsymbol{\Lambda}^{-1}\mathrm{Y}\mathbf{x}\right)^{\mathsf{T}} \boldsymbol{\Lambda} \left(\boldsymbol{\beta} - \boldsymbol{\Lambda}^{-1}\mathrm{Y}\mathbf{x}\right) + \frac{1}{2\sigma^2}\mathbf{x}^{\mathsf{T}}\mathrm{Y}^{\mathsf{T}}\boldsymbol{\Lambda}^{-1}\mathrm{Y}\mathbf{x}\right) d\boldsymbol{\beta} \\
&\quad = \frac{1}{(2\pi\gamma^2)^{G/2}\, |\boldsymbol{\Lambda}|^{1/2}} \exp\left(\frac{1}{2\sigma^2}\mathbf{x}^{\mathsf{T}}\mathrm{Y}^{\mathsf{T}}\boldsymbol{\Lambda}^{-1}\mathrm{Y}\mathbf{x}\right). \qquad (17)
\end{aligned}
$$

Using Eqs. 15 and 17, we obtained the probability of the trans-eQTL model,

$$
P(\mathcal{H}_1 \mid \mathbf{x}, \mathrm{Y}) = \left(1 + \left(2\pi\gamma^2\right)^{G/2} |\boldsymbol{\Lambda}|^{1/2} \exp\left(-\frac{1}{2\sigma^2}\mathbf{x}^{\mathsf{T}}\mathrm{Y}^{\mathsf{T}}\boldsymbol{\Lambda}^{-1}\mathrm{Y}\mathbf{x}\right) \frac{P(\mathcal{H}_0)}{P(\mathcal{H}_1)}\right)^{-1} \qquad (18)
$$

Thus, it is possible to obtain the probability of each SNP being a trans-eQTL but the calculation is computationally expensive and requires the prior probabilities $P(\mathcal{H}_0)$ and $P(\mathcal{H}_1)$. These prior probabilities can be set to a constant.

#### 2.4 RR-score: Definition

Motivated by the probability obtained in Eq. 18, we defined our test statistic RR-score, denoted $q_{\mathrm{rev}}$, as follows:

$$
\begin{aligned}
q_{\mathrm{rev}} &= \frac{1}{\sigma^2}\mathbf{x}^{\mathsf{T}}\mathrm{Y}^{\mathsf{T}}\boldsymbol{\Lambda}^{-1}\mathrm{Y}\mathbf{x} \\
&= \mathbf{x}^{\mathsf{T}}\mathbf{W}\mathbf{x} \qquad (19) \\
\text{where, } \mathbf{W} &:= \frac{1}{\sigma^2}\mathrm{Y}^{\mathsf{T}}\left(\mathrm{Y}\mathrm{Y}^{\mathsf{T}} + \frac{\sigma^2}{\gamma^2}\mathbb{I}_G\right)^{-1} \mathrm{Y}\,. \qquad (20)
\end{aligned}
$$

Suppose we have obtained a score $q_{\mathrm{rev}}$ for the $i^{\mathrm{th}}$ SNP and we would like to know how significant this score is. In order to rank it with respect to all the other SNPs in the genome, we need to calculate a null distribution of $q_{\mathrm{rev}}^{\mathrm{null}}$ and from it the $p$-value of our observed score $q_{\mathrm{rev}}$.

Note that the statistic $q_{\mathrm{rev}}$ is not proportional to the probability given by Eq. 18 and hence cannot be directly used for ranking the SNPs. However, in Sec. 2.6, we derive a $p$-value to ascertain the significance of the $q_{\mathrm{rev}}$ of each SNP.

#### 2.5 Numerical calculation

The RR-score defined in Eq. (19) involves computing the inverse of a $G \times G$ matrix $\mathbf{W}$. To reduce the complexity, we first perform a singular value decomposition of $\mathrm{Y}^{\mathsf{T}}$,

$$
\mathrm{Y}^{\mathsf{T}} = \mathrm{U}^{\mathsf{T}}\mathrm{S}\mathrm{V} \qquad (21)
$$

with two orthogonal matrices, $\mathbf{U} \in \mathbb{R}^{N\times N}$ and $\mathbf{V} \in \mathbb{R}^{G\times G}$, and a diagonal matrix $\mathbf{S} \in \mathbb{R}^{N\times G}$ that whose all elements are zero except for the $K = \min\{G, N\}$ singular values $s_k$ on its diagonal. Substituting the SVD into $\mathbf{W}$ gives

$$\mathbf{W} = \frac{1}{\sigma^2}\mathbf{U}^\mathsf{T}\mathbf{S}\left(\mathbf{S}^\mathsf{T}\mathbf{S} + \frac{\sigma^2}{\gamma^2}\mathbb{I}_G\right)^{-1}\mathbf{S}^\mathsf{T}\mathbf{U} \tag{22}$$

This allows us to expand the matrix $\mathbf{W}$ using the singular values $s_k$ and the eigenvectors $\mathbf{u}_k$,

$$\mathbf{W} = \frac{1}{\sigma^2}\sum_{k=1}^{K}\frac{s_k^2}{s_k^2 + \sigma^2/\gamma^2}\mathbf{u}_k\mathbf{u}_k^\mathsf{T} \tag{23}$$

Therefore, we can write the RR-score as

$$q_{\text{rev}} = \frac{1}{\sigma^2}\sum_{k=1}^{K}\frac{s_k^2}{s_k^2 + \sigma^2/\gamma^2}\left(\mathbf{u}_k^\mathsf{T}\mathbf{x}\right)^2 . \tag{24}$$

#### 2.6 Null model

To obtain a $p$-value for the $q_{\text{rev}}$ score of SNP $i$ with column $\mathbf{x}_i$ in the genotype matrix $\mathbf{X}$, we need a null distribution of $q_{\text{rev}}$ scores. The $p$-value is then the probability mass of the null distribution with values higher than the actually obtained $q_{\text{rev}}$ score. We can obtain such null distribution by permuting the genotype entries in $\mathbf{x}_i$ while keeping the rows of the gene expression matrix $\mathbf{Y}$ unpermuted. The distribution of the resulting $q_{\text{rev}}^{\text{null}}$ scores can differ between SNPs depending on their minor allele frequency (MAF) and the variance of the genotype ($\sigma^2$).

In principle we could derive the distribution of $q_{\text{rev}}^{\text{null}}$ empirically by permuting the elements of $\mathbf{x}_i$ a large number of times. However, to obtain $p$-values of $5 \times 10^{-8}$ or below, corresponding to genome-wide significance, we would need to draw at least $2 \times 10^7$ permuted samples and compute their $q_{\text{rev}}$ scores. This would be too time consuming. Instead, we derive in Appendix 1 analytical expressions for the expectation value $\mu_q := \left\langle q_{\text{rev}}^{\text{null}}\right\rangle$ and variance $\sigma_q^2 := \text{Var}\left[q_{\text{rev}}^{\text{null}}\right]$ of $q_{\text{rev}}^{\text{null}} = \mathbf{x}^\mathsf{T}\mathbf{W}\mathbf{x}$ under the permutation null model for any symmetric matrix $\mathbf{W}$ and any centered vector $\mathbf{x}$. In Fig. S1, we show that our analytical calculation of $\mu_q$ and $\sigma_q$ match those obtained from the empirical permutation of $\mathbf{x}$.

**Normal approximation.** Under the null model assumption that the SNP is not a trans-eQTL, the $\mathbf{u}_k^\mathsf{T}\mathbf{x}$ are distributed as the projections of a unit vector with random direction onto $K$ Cartesian coordinates given by the eigenvectors $\mathbf{u}_k$. The $\mathbf{u}_k^\mathsf{T}\mathbf{x}$ will be normally distributed and $\left(\mathbf{u}_k^\mathsf{T}\mathbf{x}\right)^2$ will have a chi-square distribution. In the limit of $K \gg 1$, the sum over $\left(s_k^2/(s_k^2 + \sigma^2/\gamma^2)\right)\left(\mathbf{u}_k^\mathsf{T}\mathbf{x}\right)^2$ will have a normal distribution according to the central limit theorem. Therefore, we approximate $q_{\text{rev}}^{\text{null}}$ by $\mathcal{N}\left(\mu_q, \sigma_q^2\right)$. Finally, the $p$-value of $q_{\text{rev}}$ for a candidate SNP is

$$p \approx \Phi\left(\frac{q_{\text{rev}} - \mu_q}{\sigma_q}\right) , \tag{25}$$

where $\Phi(z)$ denotes the cumulative normal distribution for a random variable $z$.

**Constraint on covariate correction.** Generally, the gene expression matrix $\mathbf{Y}$ should have rank $(\mathbf{Y}) = K = \min\{G, N\}$. This means that the SVD in Eq. (21) should have $K$ singular values $s_k$. To obtain $q_{\text{rev}}$, the sum in Eq. (24) runs over $K$ components of $\left(\mathbf{u}_k^\mathsf{T}\mathbf{x}\right)^2$. If $C$ known covariates are now corrected from $\mathbf{Y}$ using linear regression (see CCLM below, Sec. 3.1), then $C$ of these singular

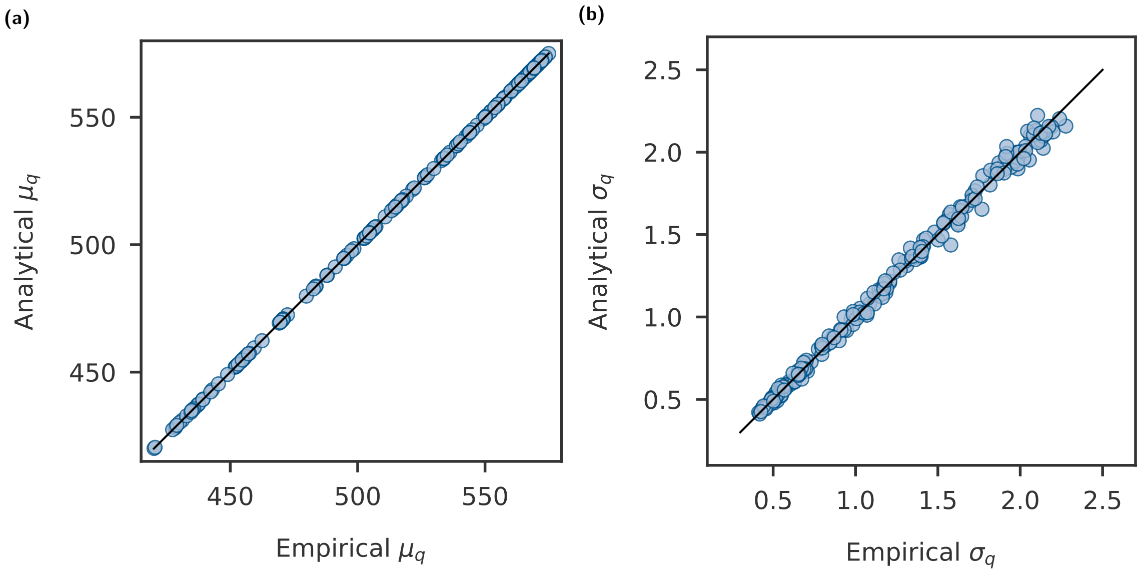

**Fig. S1 | Validation of analytical calculation of mean and variance of RR-score null distribution.** We compare the analytical calculation of **(a)** mean $\mu_q$ and **(b)** standard deviation $\sigma_q$ of $q_{\text{rev}}^{\text{null}}$ with those obtained empirically for 200 randomly selected SNPs (GTEx data) with different minor allele frequencies. Each point represents a SNP. The empirical null distribution of $q_{\text{rev}}$ for each SNP was created by permuting the genotype 500 times. Here, we used the gene expression of adipose subcutaneous tissue from GTEx.

values become zero for the residual expression $\mathbf{Y}'$. This is equivalent to subtracting $C$ components from the sum in Eq. (24). When the subtracted part is not normally distributed, as would happen if $C$ is too small for the central limit theorem to be valid, then $q_{\text{rev}}^{\text{null}}$ is not normally distributed and the $p$-values are not accurate. If $C$ is large, then the subtracted part and consequently, $q_{\text{rev}}^{\text{null}}$, remains normally distributed. In practical situations, only a few known covariates are used to correct the gene expression and in this limit of small $C$, the null model is not Gaussian.

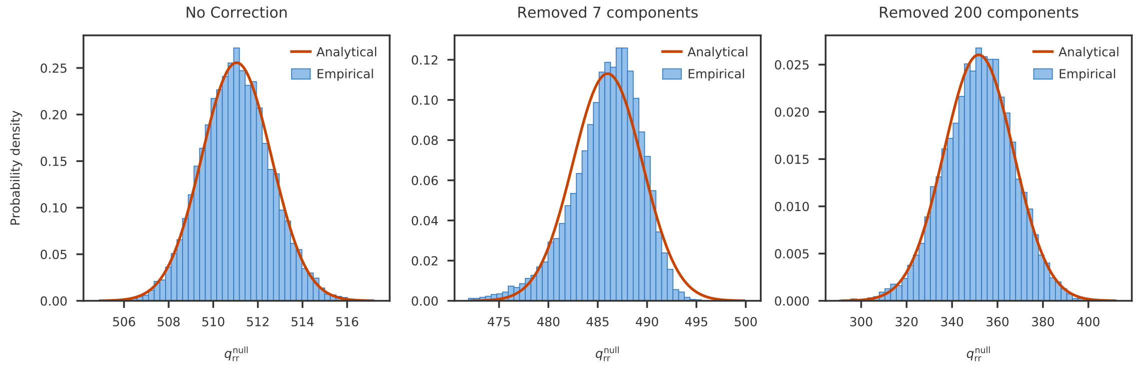

**Fig. S2 | Effect of covariate correction on null model.** We calculate the null model empirically for a SNP with minor allele frequency (MAF) 0.1 using the gene expression from adipose subcutaneous tissue (581 samples and 15763 genes) of GTEx. The blue histogram is the null distribution calculated empirically and the solid red line shows the analytically approximated normal distribution $\mathcal{N}(\mu_q, \sigma_q^2)$. Our analytical model is indeed valid without any covariate correction (left panel). When the gene expression is corrected for a few covariates, then the normal approximation breaks down as explained in the text (center panel). However, when a large number of singular values are removed, then the central limit theorem becomes valid again (right panel).

In Fig. S2, we show that $q_{\text{rev}}^{\text{null}}$ is normally distributed for a given SNP and a gene expression matrix with $N = 581$ (from adipose subcutaneous tissue of GTEx). Setting the first seven singular values to zero (equivalent to correcting seven covariates) breaks the normal distribution of $q_{\text{rev}}^{\text{null}}$. However, if 200 singular values are set to zero, then the assumption of normal distribution of $q_{\text{rev}}^{\text{null}}$ becomes valid again. This illustrates the problem of using CCLM with few confounders in combination with Tejaas.

#### 2.7 Choice of the regularizer

The standard deviation $\gamma$ of the normal prior is not learnt from the data, but is chosen empirically. It is important to choose $\gamma$ such that it avoids overfitting and also restricting the regression coefficients too much..

In the limit of large $\gamma$, the normal approximation to $q_{\text{rev}}$ will break down. To see this, we first observe that from Eq. (24)

$$q_{\text{rev}} < \frac{1}{\sigma^2} \sum_{k=1}^{K} (\mathbf{u}_k^{\mathsf{T}} \mathbf{x})^2 = \frac{\|\mathbf{x}\|^2}{\sigma^2} = 1. \tag{26}$$

Therefore, as $\sigma^2/\gamma^2$ becomes smaller than most squared singular values $\{s_k^2 : 1 \le k \le K\}$, $q_{\text{rev}}$ will approach 1. The distribution of $q_{\text{rev}}$ under the permutation null model will therefore become extremely skewed with an abrupt drop towards 1. Such a distribution will not be approximated well by a Gaussian, but luckily this situation corresponds to the irrelevant regime of extreme overfitting. This can be seen by noting that the prediction of $\mathbf{x}$ at the maximum likelihood solution $\hat{\boldsymbol{\beta}} = \boldsymbol{\Lambda}^{-1}\mathbf{Y}\mathbf{x} = (\mathbf{Y}\mathbf{Y}^{\mathsf{T}} + (\sigma^2/\gamma^2)\,\mathbb{I}_G)^{-1}\mathbf{Y}\mathbf{x}$ is $\mathbf{Y}^{\mathsf{T}}\hat{\boldsymbol{\beta}} \approx \mathbf{x}$, which means that the regression results in a perfect prediction on the training samples, in other words, it results in complete overfitting.

At the other extreme, when $\gamma$ gets so *small* that $\sigma^2/\gamma^2 \gg \max\{s_k^2 : 1 \le k \le K\}$, we can approximate

$$\begin{aligned} q_{\text{rev}} &\approx \frac{\gamma^2}{\sigma^4} \sum_{k=1}^{K} s_k^2 \left(\mathbf{u}_k^{\mathsf{T}} \mathbf{x}\right)^2 \\ &= \frac{\gamma^2}{\sigma^4} \mathbf{x}^{\mathsf{T}} \left(\mathbf{Y}^{\mathsf{T}}\mathbf{Y}\right) \mathbf{x} \\ &= N \frac{\gamma^2}{\sigma^4} \mathbf{x}^{\mathsf{T}} \boldsymbol{\Sigma} \mathbf{x} \end{aligned} \tag{27}$$

where $\boldsymbol{\Sigma}$ is the $N \times N$ sample covariance matrix. Since our gene expression matrix $\mathbf{Y}$ is normalized, the diagonal elements of $\boldsymbol{\Sigma}$ should be equal to 1, whereas the off-diagonal elements will be more or less randomly distributed around 0. Intuitively, we expect that in this regime $q_{\text{rev}}$ will be will be too restrictive for the model and lead to false signals even with chance correlations of a single gene with a genotype.

**Non-Gaussian parameter.** From Eq. (25), the $q_{\text{rev}}^{\text{null}}$ should have a normal distribution and the standardized $q_{\text{rev}}^{\text{null}}$ should have a standard normal distribution,

$$P\left(\frac{q_{\text{rev}}^{\text{null}} - \mu_q}{\sigma_q}\right) = \mathcal{N}(0, 1)\ . \tag{28}$$

The kurtosis of a standard normal distribution is 3. We use this property to define a non-Gaussian parameter ($\alpha$) to measure the deviation of the distribution from a standard normal distribution,

$$\alpha(\gamma) = \left| \frac{\langle f(q;\gamma)^4 \rangle}{3} - 1 \right| , \tag{29}$$

where $f(q;\gamma) = \left(q_{\text{rev}}^{\text{null}} - \mu_q\right)/\sigma_q$. Given a gene expression matrix, we simulated genotypes for 5000 SNPs and calculated $\alpha(\gamma)$ for different values of $\gamma$. We recommend choosing $\gamma_{\text{opt}}$ such that $\gamma_{\text{opt}} \geq \arg\min_{\gamma} \alpha(\gamma)$. Ideally, we also want a high value for the standard deviation of $\sigma_q$ to ensure a broad distribution of $q_{\text{rev}}^{\text{null}}$.

#### 2.8 Target genes

Reverse regression offers evidence for the presence of a trans-eQTL without identifying *which* genes are targeted. Hence, we do not get the set of target genes from reverse regression. In fact, multiple regression with $L_2$ penalty (ridge regression) is not optimal for variable selection. Therefore, we have included a secondary step in Tejaas to perform single SNP-gene pairwise regression to find a set of candidate genes as targets for each trans-eQTL discovered.

As discussed below (Sec. 3), reverse regression works better with KNN correction while pairwise regression (for target gene discovery) works better with conventional covariate-corrected gene expression. To get the best of both worlds, our software accepts two separate input options for gene expression files: (1) `--gx` to provide the raw gene expression file for KNN correction and trans-eQTL discovery, and (2) `--gxcorr` to provide the covariate-corrected gene expression file for single SNP-gene pair regression to discover target genes.

#### 2.9 Cis-masking

In Tejaas, we are interested to find long-range SNP-gene interactions. However, the strong effect size of the cis-eQTLs can sometimes lead to high $q_{\text{rev}}$. To avoid wrongly identifying cis-eQTLs as trans-eQTLs, we introduced an option in our software to exclude all genes in the vicinity of each SNP. We call this procedure 'cis-masking'. It can be invoked with the flag `--cismask`. The width of the vicinity can be specified using the option `--window`.

To implement cis-masking, we calculated the SVD of the expression matrix (see Eq. (21)) after removing (masking out) the genes that occur within ±1Mb (or any distance specified by `--window`) from the SNP position. Doing this for every SNP would be computationally expensive as it would involve $\sim 10^7$ matrix inversions (once for each SVD). But since many SNPs generally have identical genes in the vicinity, we group them and calculate the SVD once for each group, which brings down the number of SVD calculations to $\sim 10^4$.

#### 2.10 Computational requirements

To determine run times and memory requirements of Tejaas, we used the adipose subcutaneous tissue expression from GTEx, which contains a total of 15673 genes for 581 samples. We used the GTEx genotype for chromosome 1. All tests were run on a node with 2× Intel Xeon CPU E5-2640 v3 @ 2.60GHz with 8 physical cores each and 128Gb of RAM. We subsampled the expression matrix as well as the number of SNPs used to run Tejaas to evaluate different possible scenarios.

The most expensive step for Tejaas is the SVD decomposition of the gene expression matrix. The number of times the SVD decomposition is performed depends directly on the number of SNPs selected. While using `--cis-masking` (Sec. 2.9), different SNPs will have different cis-genes that are required to be masked out and hence, a new SVD decomposition is required. Run times increase with the number of SNPs (Fig. S3a, left panel), since the number of cismasks needed (top x-axis) is proportional to the number of SNPs (bottom x-axis). Memory usage increases with the number of SNPs and samples, since the dosage matrix is being held in memory.

We implemented Tejaas with an option to run Tejaas on a multicore server using MPI parallelization. Fig. S3b shows how run times decrease with the number of cores used. Total memory usage, as expected, increases with the number of cores. In the designed parallelization scheme, each core will compute the SVD decomposition for a given cismask. With an increase in the number of cores, the expression matrix needs to be copied to each of them for processing and hence the memory usage scales proportionally.

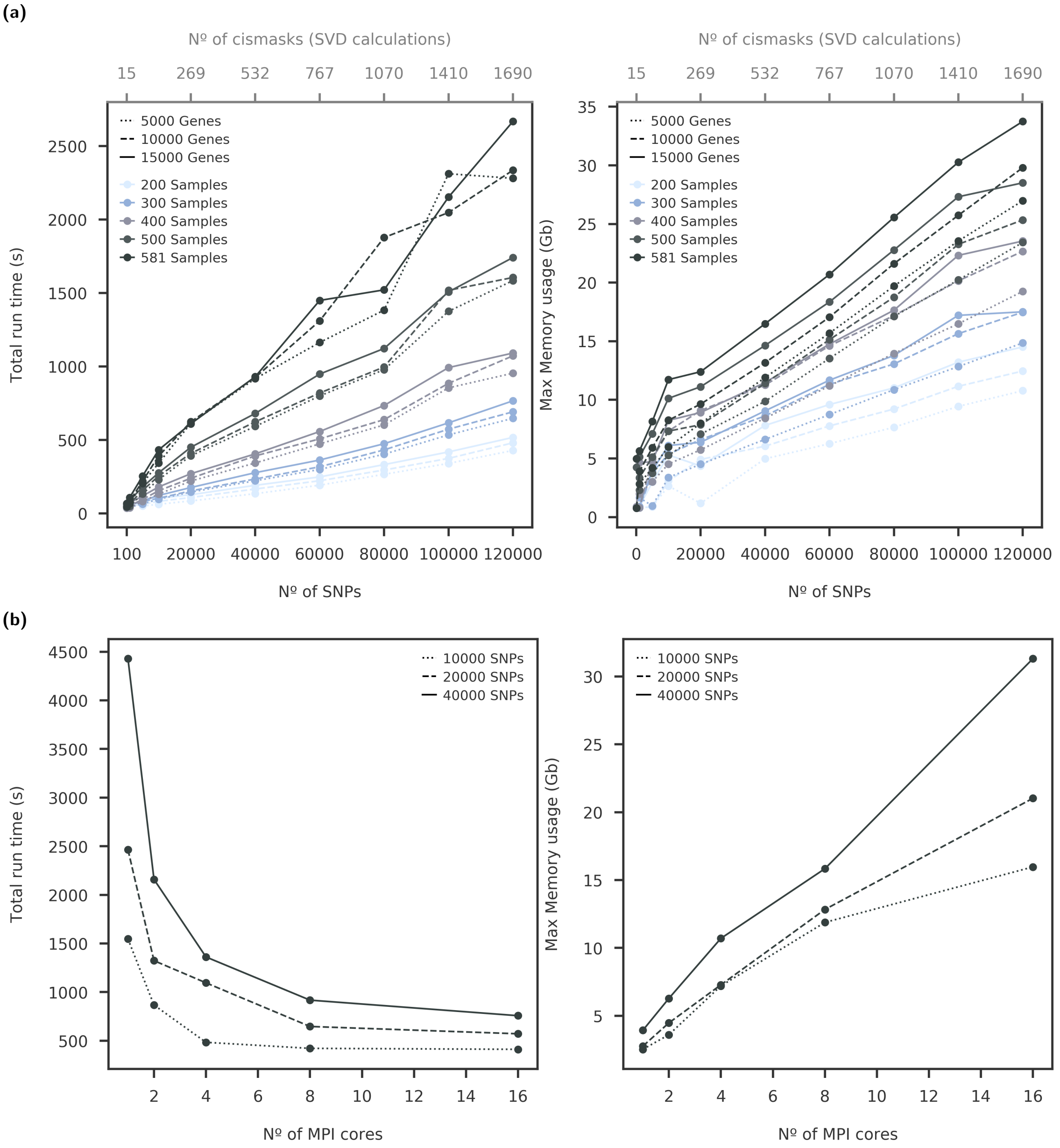

**Fig. S3 | Computational time and memory requirements of Tejaas. (a)** Total run time and maximum memory usage as a function of SNPs are shown on the left and right panels, respectively. All runs of Tejaas used 8 CPU cores for parallelization. The top x-axis reflects how many cismasks are required in chromosome 1 for a given number of SNPs. **(b)** Total run time and maximum memory usage as a function of the number of CPU cores. All runs of Tejaas used 15673 genes for 581 samples.

### 3 Confounder correction

Gene expression measurements are notorious for being dominated by strong confounding effects that can arise from technical details of RNA recovery, conservation, and sequencing or from environmental and biological factors such as age, gender, diseases, nutrition and lifestyle, drug regimes etc. The subtle effects of trans eQTLs are at risk of being drowned out by the strong systematic noise from confounding effects.

In this section, we first discuss the standard confounder correction method for eQTL analyses. Then, we introduce our K-nearest neighbor correction that is specially suitable for use with Tejaas.

#### 3.1 Residuals from multiple linear regression

Given a matrix $\mathbf{C} \in \mathbb{R}^{C \times N}$ of $C$ covariates and $N$ samples, a multiple linear regression is performed for the $g^{\text{th}}$ gene expression $\mathbf{y}_g$,

$$\mathbf{y}_g = \boldsymbol{\alpha}_g \mathbf{C} + \boldsymbol{\xi}_g , \tag{30}$$

where $\boldsymbol{\alpha}_g \in \mathbb{R}^C$ is a vector of effect sizes of the covariates for gene $g$. The estimate $\widehat{\boldsymbol{\alpha}_g}$ of the effect sizes from the multiple linear regression is then used to calculate the expression residuals $\mathbf{y}'_g = \mathbf{y}_g - \widehat{\boldsymbol{\alpha}_g}\mathbf{C}$ which are linearly uncorrelated with the covariates. The residual $\mathbf{y}'_g$ is used as the corrected gene expression for downstream analyses. In this manuscript, we denote this correction as 'CCLM'.

This is a simple and effective method to correct gene expression confounders and is routinely used to discover biologically significant eQTLs. CCLM can be used for known covariates such as age, gender, etc. as well as other hidden covariates inferred using PEER [2] or other similar methods. However, if $\operatorname{rank}(\mathbf{Y}) = N$, then it can be shown that for the residual matrix, $\operatorname{rank}(\mathbf{Y}') \le N - C$. Therefore, as noted in Sec. 2.6, this may compromise the normality of the distribution of $q_{\text{rev}}^{\text{null}}$. Hence, the CCLM correction is problematic to use with Tejaas.

#### 3.2 K-nearest neighbor (KNN) correction

For the KNN correction, we assume that confounding effects dominate the gene expression. We expect that the expression levels of the genes of $K$ nearest neighbors ($\text{NN}_n^K$) of each sample $n$ will be affected by the same dominant confounder variables. In other words, if the samples are close to one another in the expression space, we expect them to be close to one another in the confounder space. Hence, we can correct at least a good part of the confounding effects by centering the expression $\mathbf{y}_n$ and genotype $\mathbf{x}_n$ of sample $n$ using the $K$ samples whose gene expressions are the most similar to that of sample $n$,

$$\mathbf{y}_n \leftarrow \mathbf{y}_n - \frac{1}{K} \sum_{m \in \text{NN}_n^K} \mathbf{y}_m \tag{31}$$

$$\mathbf{x}_n \leftarrow \mathbf{x}_n - \frac{1}{K} \sum_{m \in \text{NN}_n^K} \mathbf{x}_m . \tag{32}$$

The nearest neighbors $\text{NN}_n^K$ are calculated using the denoised Euclidean distances between gene expression vectors. To reduce noise in the distance calculation, we remove all but the leading $M$ principle components of the singular value decomposition of the expression matrix $\mathbf{Y}$. After subtracting the mean gene expressions and genotype vectors of the $K$ nearest neighbors, we center the resulting corrected gene expression and genotype vectors again.

The choice of $K$ should be such that it captures the locally varying effects of the confounders. As shown later in Fig. S7, too small a value of $K$ would lead to excessive statistical noise, while increasing $K$ too much will remove too little of the confounding effects as these start to average out within the $K$ neighbors of each data point.

In Fig. S4, we show how KNN correction removes the strong dissimilarity between samples in the data. Before KNN correction (Fig. S4a), the samples are strongly dissimilar. We could observe

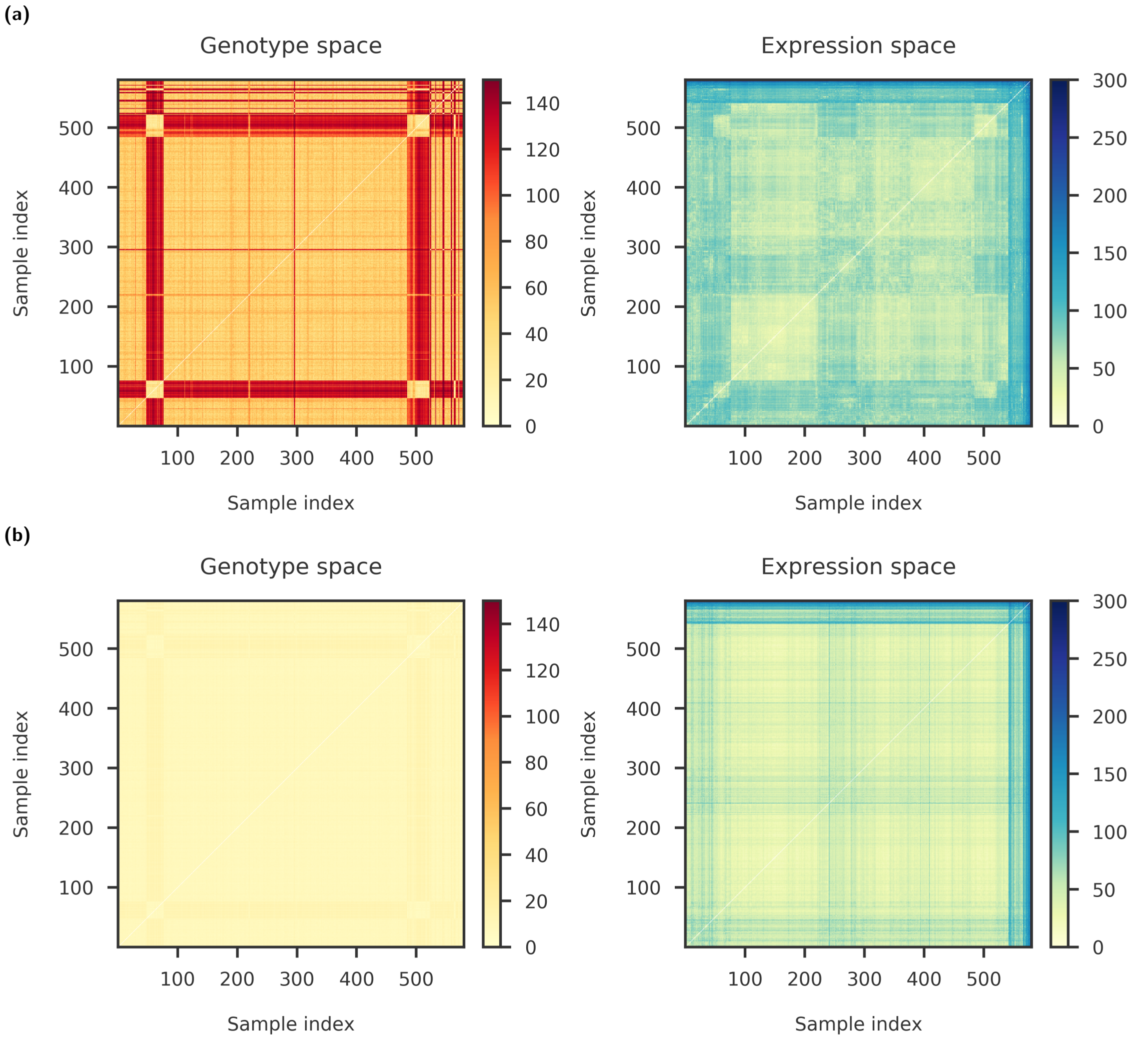

**Fig. S4 | Distance between samples before and after KNN correction.** Heatmaps with color indicating the distance between samples ($N = 581$) in both the genotype space (using LD pruned genotype from GTEx) and the expression space (using adipose subcutaneous tissue from GTEx): **(a)** Before KNN correction, and **(b)** after KNN correction. The samples are clustered using their distance in the expression space before KNN correction. For the example shown here, we reduced the expression matrix to 30 dimensions ($M = 30$) and considered 30 nearest neighbors ($K = 30$) for centering the genotype and expression.

several sample groups which are neighbors in the expression space, probably due to external confounders. Although the samples were clustered in the expression space, we observed distinctive patches in the genotype space. This happens because several confounders (such as population substructure) affect both genotype and gene expression, leading to unwanted correlation between the first principal component of the genotype and the expression levels. We could correct for these confounders by centering both $\mathbf{x}_n$ and $\mathbf{y}_n$ in the KNN method (Fig. S4b).

The KNN correction has several benefits in comparison to CCLM and other linear corrections. First, it can correct out non-linear confounding effects, so, in contrast to the CCLM correction, it should also work when the confounding effects are not well approximated by linear, additive effects. Second, since it is non-parametric (except for $K$), overfitting does not typically occur [3, 4]. Third, it does not require the confounders to be known. Fourth, since KNN does not reduce the rank of the gene expression matrix, it works well with Tejaas.

### 4 Trans-eQTL simulation

This section describes details of a simulation study that evaluates the power of our method to find *trans* effects in gene expression data. We followed the strategy of Hore *et al.* for the simulation, as described in the Supplementary section of their article [5]. We discuss their simulation details below, partly verbatim. Any difference with their strategy is explicitly noted.

#### 4.1 Data simulation

Simulated data consisted of genotype and gene expression for $N = 450$ individuals to closely resemble the sample size of the Genotype Tissue Expression (GTEx) project [6–8]. We simulated the gene expression data for $G = 12\,639$ genes, containing non-genetic signals (background correlation estimated from the GTEx tissues and confounding factors) and genetic signals (*cis* and *trans* effects). Following the strategy of Hore *et al.* [5], we assumed that each gene contained only one SNP; this simplified case is equivalent to assuming that there is at most one cis-eQTL for each gene.

##### 4.1.1 Genotypes

We used the real genotype data from the GTEx individuals, while Hore *et al.*used simulated genotypes. After pre-filtering of the GTEx genotype, we randomly sampled $I = 12\,639$ SNPs (corresponding to $G = 12\,639$ genes) from the genotype data of 450 individuals. From the $I$ SNPs sampled, we randomly selected $I_{\text{cis}} = 800$ SNPs to be cis-eQTLs. From these cis-eQTLs, we selected a subset $I_{\text{trans}} = 30$ SNPs to be trans-eQTLs. These trans-eQTLs were originally cis-eQTLs associated with a nearby gene, which in turn regulated the expression of other distant genes. All SNPs were sampled independently from the genotype. Let $\mathbf{X} \in \mathbb{R}^{I \times N}$ be the sampled matrix of genotypes.

##### 4.1.2 Gene expression

Let $\mathrm{Y} \in \mathbb{R}^{G \times N}$ be the matrix of simulated gene expression data. The data was simulated in stages. First, we generated the non-genetic component of all genes using background correlation (noise) and confounding factors. Second, we added the cis effects and finally, we added the *trans* component on the target genes of the trans-eQTLs.

**Noise.** Hore *et al.*used heteroscedastic background noise. However, to replicate the underlying correlation of the gene expressions in the GTEx samples, we created a Gaussian noise with a covariance matrix obtained from the gene expressions in the artery aorta tissue of GTEx. Let $\mathrm{Y}_{\text{as}}$ be the observed gene expression in GTEx samples. We decomposed the covariance matrix of $\mathrm{Y}_{\text{as}}$ such that

$$\mathrm{Cov}\,(\mathrm{Y}_{\text{as}}) = \mathbf{Q}\boldsymbol{\Lambda}\mathbf{Q}^{\mathsf{T}} \tag{33}$$

We defined the noise as

$$\mathbf{Y}^{\text{noise}} = \mathbf{Q}\sqrt{\boldsymbol{\Lambda}}\mathbf{Z} \tag{34}$$

where $\mathbf{Z}$ is a $G \times N$ matrix, with every row sampled from a normal distribution with zero mean and unit variance. Note that $\mathrm{Cov}\,(\mathrm{Y}_{\text{noise}}) = \mathrm{Cov}\,(\mathrm{Y}_{\text{as}})$ and therefore $\mathbf{Y}^{\text{noise}}$ retains the background correlation structure of the adipose subcutaneous tissue.

**Confounding factors.** We generated $C = 10$ confounding factors, of which $C_{\text{pop}} = 3$ were considered to be due to population substructure. Let $\mathbf{W}$ be the $C \times N$ matrix of confounding factors. The three population substructure confounders ($\mathbf{W}^{\text{pop}}$) were set to be the first, second and third principal components of the corresponding genotype matrix. The remaining $C_{\text{ind}} = C - C_{\text{pop}}$ independent confounders ($\mathbf{W}^{\text{ind}}$) were sampled from a standard normal distribution,

$$\mathbf{W}_i^{\text{ind}} \sim \mathcal{N}(0, 1) \tag{35}$$

Each confounding factor was assumed to be affecting only a few target genes. Hence, the effect size $\beta_{cg}$ for the $c^{\text{th}}$ confounding factor on the $g^{\text{th}}$ gene was sampled with a sparsity $\alpha_c$,

$$\beta_{cg} \sim \alpha_c \mathcal{N}\left(0, \sigma_c^2\right) + (1 - \alpha_c)\,\delta_0, \;\; \alpha_c \sim \text{Beta}(2, 5) \tag{36}$$

to give us the coefficient matrix $\boldsymbol{\beta}$ of size $C \times G$. We used $\sigma_c = 1.0$ unless otherwise mentioned. Finally, we obtained the effect of confounding factors on the gene expression using an additive model,

$$\mathrm{Y}^{\text{cf}} = \boldsymbol{\beta}^{\mathrm{T}}\mathbf{W} \tag{37}$$

**Cis effects.** In this simulation framework, every gene spatially overlaps with a corresponding SNP in the genotype, *i.e.*, we have $I$ (= $G$) SNP-gene pairs. If the SNP in the $i^{\text{th}}$ position has a cis effect ($i \in I_{\text{cis}}$), then the expression of the gene $i$ is modified with a strength $\alpha_i$,

$$\mathrm{Y}_i^{\text{cis}} = \alpha_i \mathrm{X}_i \tag{38}$$

where the direction of the effect size is random, *i.e.*, $\text{sgn}(\alpha_i)$ is assigned a value of $-1$ or $+1$ randomly. The strength of the cis effect, *i.e.*, $|\alpha_i|$ depends on whether the SNP also has a trans effect or not and is given by

$$|\alpha_i| = \begin{cases} 0.6, & \text{if SNP } i \text{ also acts as a trans-eQTL } (\in I_{\text{trans}}) \\ \text{Gamma}(4, 0.1), & \text{otherwise.} \end{cases} \tag{39}$$

For all the remaining SNP-gene pairs ($i \notin I_{\text{cis}}$) with no cis effects, $\mathrm{Y}_i^{\text{cis}} = \mathbf{0}$.

**Combining noise, confounding factors and cis effects.** We obtained a temporary set of expression levels for all genes by additively combining the noise, the effect of confounding factors and the effect of cis-eQTLs,

$$\mathrm{Y}^{\text{tmp}} = \mathrm{Y}^{\text{noise}} + \mathrm{Y}^{\text{cf}} + \mathrm{Y}^{\text{cis}} \tag{40}$$

Note that the $\mathrm{Y}^{\text{cis}}$ has non-zero values only for the targets of $I_{\text{cis}} = 800$ cis-eQTLs, and has zero values for all remaining genes.

**Trans effects.** For simulating the trans effects, Hore *et al.*assumed that the trans-eQTLs regulated a nearby gene (via cis effect) and this cis target gene was a transcription factor (TF) and regulated multiple target genes downstream (excluding other TF genes; could be any other gene including other cis target genes). This ensured that the trans-eQTLs were indirectly associated with the target genes with practically low effect sizes.

Let $M_{\text{trans}}$ be the number of target genes regulated by the TF and the target genes of every TF were selected at random. Let $S_g$ be the set of TFs which regulate the expression of the $g^{\text{th}}$ gene and the trans effect on this gene was simulated as,

$$\mathrm{Y}_g^{\text{trans}} = \sum_{j \in S_g} \beta_{gj} \mathrm{Y}_j^{\text{tmp}}, \quad \text{where} \tag{41}$$

$$\left|\beta_{gj}\right| \sim \text{Gamma}\left(\psi^{\text{trans}}, 0.02\right). \tag{42}$$

$\beta_{gj}$ was the relative effect of TF $j$ on the gene $g$. If a gene was not a target for any of the $I_{\text{trans}} = 30$ TFs, then $\mathbf{Y}_g^{\text{trans}} = \mathbf{0}$.

**Combining all of the data.** Finally, the contributions from the trans-eQTLs were incorporated to create a final set of simulated expression levels,

$$\mathbf{Y} = \mathbf{Y}^{\text{tmp}} + \mathbf{Y}^{\text{trans}} \tag{43}$$

and was used for subsequent analyses.

#### 4.2 ROC pAUC analysis

We sorted the trans eQTL predictions from each tool by descending score and computed the cumulative number of true and false positives (TP, FP) up to each score. We obtained the true positive rate (TPR) as TP divided by maximum number of TPs and the false positive rate (FPR) as FP divided by the maximum number of FPs. The Receiver Operating Characteristic (ROC) curve is the curve of TPR versus FPR. To measure predictive performance of the methods we used the partial area under the ROC curve up to an FPR of 0.1, since larger FPRs are irrelevant in practice.

#### 4.3 Methods compared

We compared Tejaas RR-score with two methods: (1) MatrixEQTL [9] as a representative for current standard eQTL pipelines, and (2) Tejaas FR-score, as an alternative for CPMA [1] which uses the same underlying property of trans-eQTLs that they target multiple genes simultaneously.

**MatrixEQTL.** We used the R package for MatrixEQTL with default options. The covariate correction was done separately but used the same linear model correction implemented in MatrixEQTL. Since we were not interested in the cis-eQTLs, we used a cis-window of 0Kb.

**FR (~CPMA).** Since CPMA is not implemented as a software, we implemented the FR-score ($q_{\text{fwd}}$) within Tejaas as an alternative. Internally, the method runs over the data twice: (1) First, create an empirical null model from the data and (2) then calculate $q_{\text{rev}}$ as well as an empirical $p$-value using the previously generated null model. This is the same procedure as CPMA, as described in their manuscript [1].

#### 4.4 Supplementary simulation results

**Selecting the regularizer.** We plotted the non-Gaussian parameter $\alpha(\gamma)$ and the standard deviation of $\sigma_q$ at different values of $\gamma$ in Fig. S5a. As noted above in Sec. 2.7, the best choice of $\gamma$ should be greater than $\arg\min_\gamma \alpha(\gamma)$. Ideally, we also want a high value for the standard deviation of $\sigma_q$ to ensure a broad distribution of $q_{\text{rev}}^{\text{null}}$. Therefore, we chose $\gamma = 0.2$.

To validate our choice, we looked at the ranking of the trans-eQTLs at different values of $\gamma$ in Fig. S5b. Each ROC curve was obtained by averaging over 20 simulations using default simulation parameters (see above) and KNN correction with 30 neighbors. In the inset, we show the partial areas under the ROC curves (pAUC) where the false positive rate (FPR) $\leq 0.1$. We found that Tejaas works best at this choice of $\gamma$. The rest of the simulations were performed using $\gamma = 0.2$.

**Note on inverse normal transform.** Current GTEx pipeline uses a two-step approach for normalizing the read counts of the genes: (1) read counts are normalized using TMM (Trimmed Mean of M-values [10]) and filtered for thresholds, (2) expression values for each gene are then

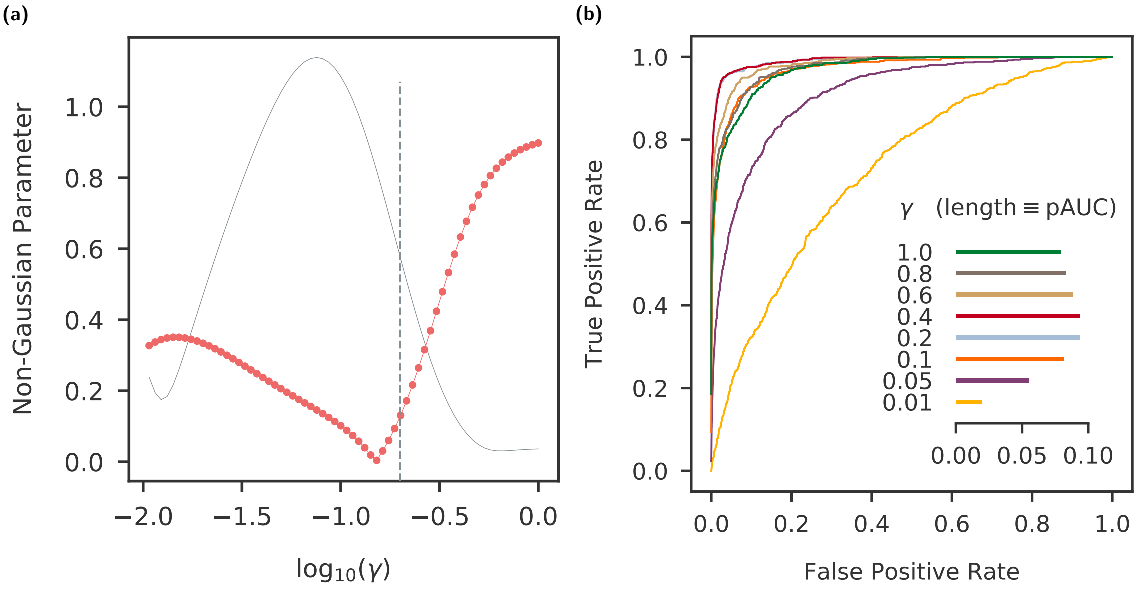

**Fig. S5 | Selection of standard deviation of the regularizer. (a)** To choose $\gamma$, we plotted the non-Gaussian parameter (red) at different values of $\gamma$. The solid gray line shows the standard deviation of $\sigma_q$. We chose $\gamma$ = 0.2 shown by the dotted line. **(b)** We validated our choice by checking the ROC curve averaged over 20 simulations at different values of $\gamma$. In the inset we show the partial area under the ROC curve (pAUC) up to a false positive rate (FPR) of 0.1.

inverse normal transformed across samples. The gene expression generated in our simulations are not read counts, but equivalent to the TMM values. Hence we skipped the first step. We did perform the second step of inverse normal transformation before the confounder corrections.

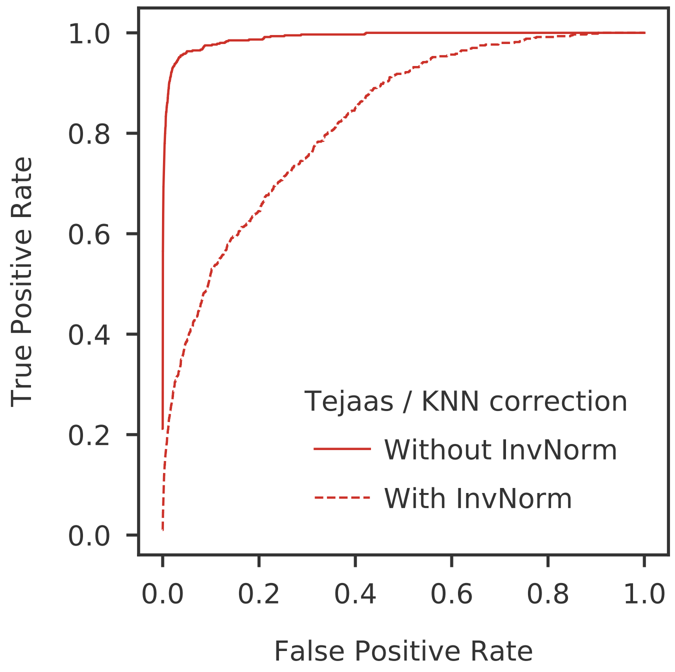

**Fig. S6 | KNN correction after inverse normal transform reduces power of Tejaas.** We plot the ROC curves averaged over 20 simulations by applying KNN with 30 nearest neighbors with and without inverse normal transformation (InvNorm) of the gene expression data.

As expected, we found that CCLM correction (Sec. 3.1) benefits significantly by using the inverse normal transformation (not shown). For the KNN correction (Sec. 3.2), however, we found that the inverse normal transformation reduces the accuracy of Tejaas (Fig. S6). It might be because the neighbor information gets skewed with this transformation. Hence, we performed KNN correction without the inverse normal transformation both for the simulations and the GTEx application.

**Number of neighbors for KNN correction.** Another important aspect of Tejaas is to choose the number of neighbors for the unsupervised KNN correction. To ascertain the robustness of the KNN correction, we performed simulations with different number of confounding factors, $C = 10, 20$ and 30. We then compared the ranking of Tejaas using different number of nearest neighbors, as shown in Fig. S7. The rankings were compared using the pAUC where the false positive rate (FPR) $\leq 0.1$. We found that the choice of KNN neighbors does not depend on the number of confounding factors, and we obtained the best accuracy with 30 nearest neighbors.

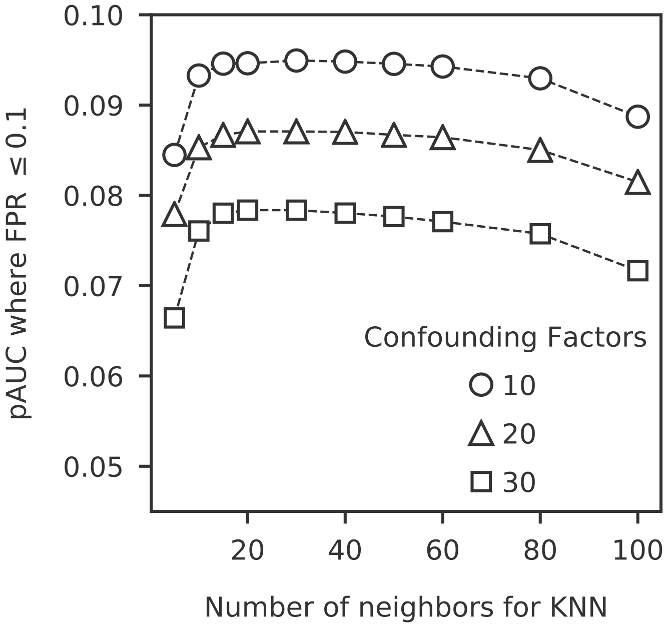

**Fig. S7 | Number of neighbors used for KNN correction.** We performed simulations with different number of confounding factors, $C = 10, 20$ and 30. The ranking of Tejaas using different number of nearest neighbors for KNN correction was measured using the pAUC where the false positive rate (FPR) $\leq 0.1$, averaged over 20 simulation replicates.

**Dependence on confounding factors.** In Fig. S8, we compared different methods for discovering trans-eQTLs at different levels of confounding. The varying confounding effects were simulated by tuning: (1) the number of covariates, $C = 10, 20$ and 30, and (2) the effect size of the covariates on the gene expression obtained by varying the standard deviation ($\sigma_c = 0.4, 0.6, 0.8$ and 1.0) of the normal distribution (see Eq. (36)) from which the effect size is sampled. For discovering trans-eQTLs, Tejaas used KNN correction directly on the gene expression and $q_{\mathrm{rev}}$ with $\gamma = 0.2$. For MatrixEQTL and FR (~CPMA), all the known covariates were corrected using CCLM on the inverse normal transformed gene expression. We compared the accuracy of the methods using the partial area under the ROC curve (pAUC) where the false positive rate (FPR) $\leq 0.1$.

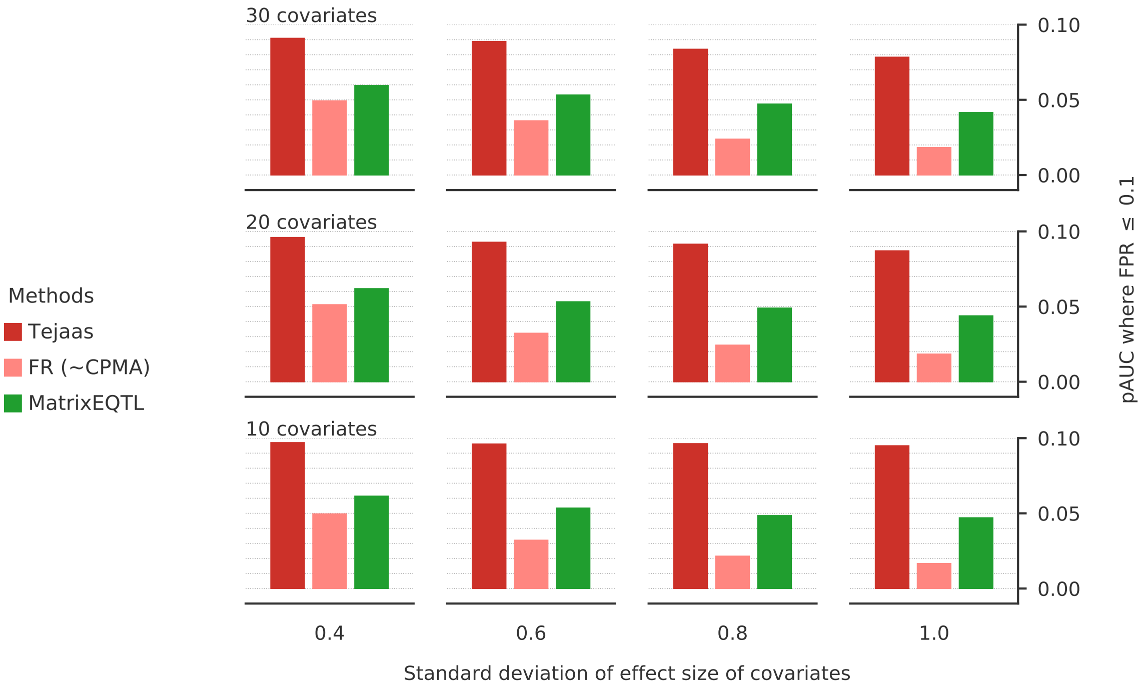

**Fig. S8 | Effect of covariates in finding trans-eQTLs.** The varying confounding effects were simulated by tuning: (1) the number of covariates, $C$ = 10, 20 and 30, and (2) the effect size of the covariates on the gene expression obtained by varying the standard deviation ($\sigma_c$ = 0.4, 0.6, 0.8 and 1.0) of the normal distribution from which the effect size is sampled. For discovering trans-eQTLs, Tejaas used KNN correction directly on the gene expression and $q_{\text{rev}}$ with $\gamma$ = 0.2. For MatrixEQTL and FR (∼CPMA), all the known covariates were corrected using CCLM on the inverse normal transformed gene expression. We compared the accuracy of the methods using the partial area under the ROC curve (pAUC) where the false positive rate (FPR) $\leq$ 0.1.

### 5 GTEx analysis

To illustrate Tejaas in a real data set, we analyzed trans-eQTLs across 49 human tissues using data from the Genotype Tissue Expression version 8 (GTEx v8) project [6–8]. In this section, we explain the preprocessing steps used for the GTEx analysis and report supporting results related to our analysis.

#### 5.1 Genotype data

We downloaded the genotype files from the dbGaP portal (accession phs000424.v8.p2). The obtained genotype was quality filtered by the GTEx consortium [11]. The file downloaded from dbGaP (filename: `GTEx_Analysis_2017-06-05_v8_WholeGenomeSeq_838Indiv_Analysis_Freeze.SHAPEIT2_phased.vcf.gz`) contained 46 562 292 variants for 838 individuals. Genotype was split in chromosomes, variants with missing values were filtered out and sex chromosomes were removed. Finally, we retained only variants with minor allele frequency (MAF) ≥ 0.01 for further analysis (8 048 655 variants).

#### 5.2 RNA-Seq expression data

We downloaded the phased RNA-seq read count expression matrix from the dbGaP portal (filename: `phASER_GTEx_v8_matrix.txt.gz`). The phASER processed expression uses read-backed mapping for each RNA-seq sample, using the genotype from the same individual as reference to map and phase RNA-seq reads. For each gene, two read count values are reported, one for each expressed allele. We added up both count values and calculated TPMs (Transcripts Per Million). For quality control, we retained genes with expression values $> 0.1$ and more than 6 mapped reads in at least 20% of the samples.

#### 5.3 Covariate correction

Tejaas uses the TPMs (or TMMs) for KNN correction to remove confounders and subsequent trans-eQTL discovery. However, as discussed in Sec. 2.8, the explicit covariate-corrected gene expression is required for finding target genes of the trans-eQTLs. We downloaded the covariate files from the GTEx portal [12] (filename: `GTEx_Analysis_v8_eQTL_covariates.tar.gz`). The covariates include the first 5 principal components of the genotype (see Supplementary Material of [11] for details), donor sex, WGS sequencing platform (HiSeq 2000 or HiSeq X) and WGS library construction protocol (PCR-based or PCR-free). Additionally, from phenotype files available in dbGaP, we included donor age and post mortem interval in minutes ('TRISCHD') as covariates. We inverse normal transformed the TPMs (or TMMs) and used CCLM (Sec. 3.1) to remove the contribution of all the covariates and used the residuals for target gene discovery.

#### 5.4 Efficacy of KNN correction

One possible way to check the efficacy of the unsupervised KNN correction is to check whether it can be explained by the confounding effects of known covariates and if so, to what extent. To test this, we checked the correlation of the 'KNN confounder' with known technical and biological covariates of GTEx samples and subjects. For each tissue $t$, the KNN confounder ($\mathbf{C}_t^{\mathrm{knn}} \in \mathbb{R}^{G \times N}$) is simply the term we subtracted from the expression values of all the $G$ genes in Eq. (31),

$$\mathbf{C}_t^{\mathrm{knn}} = \frac{1}{K} \sum_{m \in \mathrm{NN}_n^K} \mathbf{Y}_{mt} \,. \tag{44}$$

For every tissue $t$, we also obtained a list of $C$ known covariates ($\mathbf{C}_{jt}^{\mathrm{gtex}} \in \mathbb{R}^{G \times N}$ where $j \in \{1, \ldots, C\}$) by combining those obtained from the GTEx portal with a subset of sample and subject covariates from dbGaP (from filenames `phs000424.v8.pht002742.v8.GTEx_Subject_Phenotypes.data_dict.xml` and `phs000424.v8.pht002743.v8.GTEx_Sample_Attributes.data_dict.xml`). Categorical covariates with $\leq 4$ categories were binarized or converted to integers otherwise. Values with 'Not Reported' or 'Unknown' status were considered as missing values and were not considered in the analysis. For each covariate, we selected only those with at least 50 observations in any given tissue.

We then performed a simple linear regression (SLR) to explain $\mathbf{C}_t^{\mathrm{knn}}$ with $\mathbf{C}_{jt}^{\mathrm{gtex}}$ as predictors using Python's `sklearn.linear_model.LinearRegression`,

$$\mathbf{C}_t^{\mathrm{knn}} = \alpha_{jt} \mathbf{C}_{jt}^{\mathrm{gtex}} + \boldsymbol{\epsilon}_{jt} \,. \tag{45}$$

From the fitted models, we obtained the coefficient of determination $r_{jt}^2$ for every covariate $j$ in tissue $t$. In other words, the $r_{jt}^2$ corresponds to the proportion of the variance of $\mathbf{C}_t^{\mathrm{knn}}$ that can be explained by the $j^{\mathrm{th}}$ known covariate in tissue $t$. The resulting $r_{jt}^2$ values are reported in Fig. S9 (bottom panel). Indeed, the variance of KNN correction are explained to different extents by several known covariates. For example, the first principal component of the genotype (PC1) and the sample's reported race both explain a significant proportion of the variance of $\mathbf{C}^{\mathrm{knn}}$. The next four principal components, PC2 to PC5 do not contribute to explaining the variance of $\mathbf{C}^{\mathrm{knn}}$. Interestingly, sample covariates related to library size and quality as well as various rates for

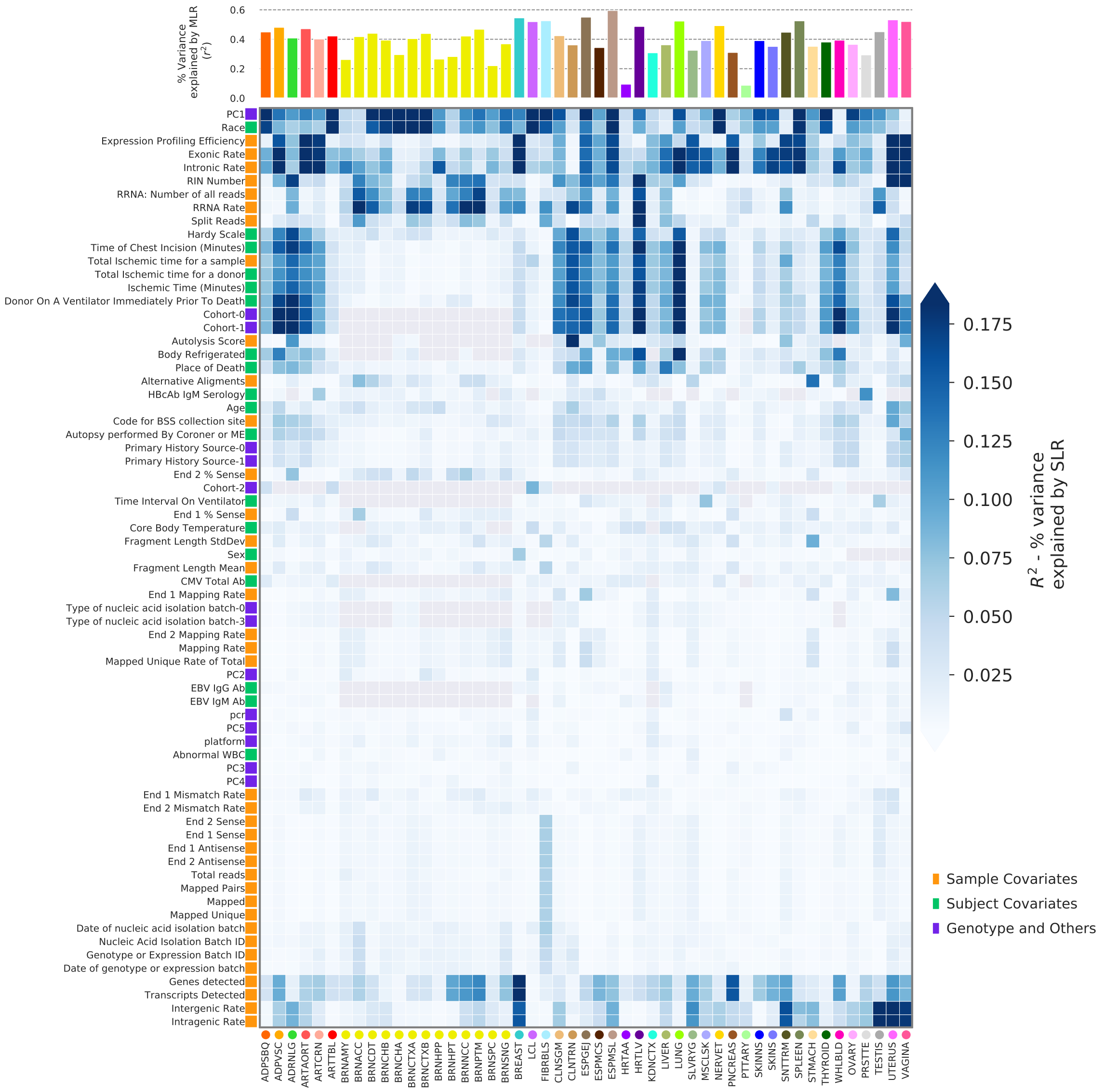

**Fig. S9 | Known covariates of GTEx can explain a significant proportion of KNN correction.** The proportion of variance of the KNN correction that can be explained by the known technical and biological covariates from GTEx can help us to understand the efficacy of KNN correction. In the bottom panel, each cell shows the coefficient of determination $r^2$ for a linear regression of the KNN confounder with the corresponding known covariate (y-axis) for each tissue (x-axis). On the top bar plot, we show the proportion of variance of the KNN confounder explained jointly by all known covariates with $r^2 > 0.05$.

mapped reads explain different proportions of the variance of $\mathbf{C}_t^{\text{knn}}$ in different tissues, indicating technical confounders in many GTEx tissues.

Next, we wanted to check the total proportion of the variance of $\mathbf{C}_t^{\text{knn}}$ explained by known covariates, which cannot be done using the SLR set up described above. For each tissue $t$, we selected those covariates with $r_{jt}^2 \geq 0.05$ in the SLR set up. We then performed a multiple linear regression (MLR),

$$\mathbf{C}_t^{\text{knn}} = \sum_{j=1}^{C'_t} \boldsymbol{\alpha}_{jt} \mathbf{C}_{jt}^{\text{gtex}} + \boldsymbol{\epsilon}\,, \tag{46}$$

where $C'_t$ is the number of covariates retained in tissue $t$. The percentage of total explained variance of $\mathbf{C}_t^{\text{knn}}$ for each tissue is shown in the top panel of Fig. S9. On average across all tissues, the known covariates that we used explained 39.9% of the variance of KNN correction.

#### 5.5 Tejaas regularizer selection for GTEx

As in simulations (Sec. 4.4), we selected the regularizer for the GTEx tissues using the non-Gaussian parameter (Sec. 2.7). From Fig. S10, we found that we could use $\gamma = 0.1$ for most of the tissues in GTEx v8. However, there were four tissues for which $\alpha(\gamma)$ at $\gamma = 0.1$ deviated strongly from the minimum. Hence for these four tissues, namely heart atrial appendage, pancreas, spleen and whole blood, we chose $\gamma = 0.006$.

#### 5.6 Method for calculating feature enrichments

We calculated whether the trans-eQTLs are enriched in several functional genomic features. For every such feature, we picked 5000 random SNPs from the GTEx genotype. and calculated the background fraction of SNPs expected by chance to have that feature,

$$f_{\text{bg}} = \frac{\text{number of SNPs with feature}}{\text{number of SNPs chosen randomly}} \tag{47}$$

This is repeated 50 times and the mean $\langle f_{\text{bg}} \rangle$ is considered as the background for that feature. Next, we calculated the fraction of trans-eQTLs with that feature,

$$f_{\text{trans-eqtl}} = \frac{\text{number of trans-eQTLs with feature}}{\text{number of trans-eQTLs in the tissue}} \tag{48}$$

Finally, we calculated the feature enrichment of trans-eQTLs in that tissue,

$$\rho_{ft} = \frac{f_{\text{trans-eqtl}}}{\langle f_{\text{bg}} \rangle} \tag{49}$$

We used a binomial test to calculate the $p$-values for the enrichment $\rho_{ft}$. If $T$ is the number of trans-eQTLs in the tissue, then the probability of finding $k$ of them with a feature is,

$$P(x = k) = \text{Binomial}\left(T, k, \langle f_{\text{bg}} \rangle\right) \, . \tag{50}$$

and $P(x > k)$ gives us the $p$-value for the tissue-GWAS pair.

#### 5.7 Regulatory element enrichment

To better understand the possible molecular mechanisms that drive trans-eQTL effects, we calculated enrichments of the trans-eQTLs at functional annotations in the genome. Annotations were obtained from GTEx Portal (`WGS_Feature_overlap_collapsed_VEP_short_4torus.MAF01.txt.gz`). For our analysis, we considered three sets of trans-eQTLs: (i) the set of all trans-eQTLs, (ii) trans-eQTLs that are only found in one tissue and (iii) trans-eQTLs that are found in two or more tissues (Fig. S11 'All', 'N1' and 'N2+', respectively). We found that trans-eQTLs are significantly enriched in enhancers, open chromatin regions, promoter flanking regions and transcription factor binding sites ($p < 2.42 \times 10^{-05}$, $p < 4.4 \times 10^{-05}$, $p < 2.42 \times 10^{-16}$ and $p < 9.9 \times 10^{-04}$ respectively). On the other hand, trans-eQTLs are only slightly enriched in promoter regions ($\rho = 1.10$, $p = 0.08$) and depleted on CTCF binding sites ($\rho = 0.86$).

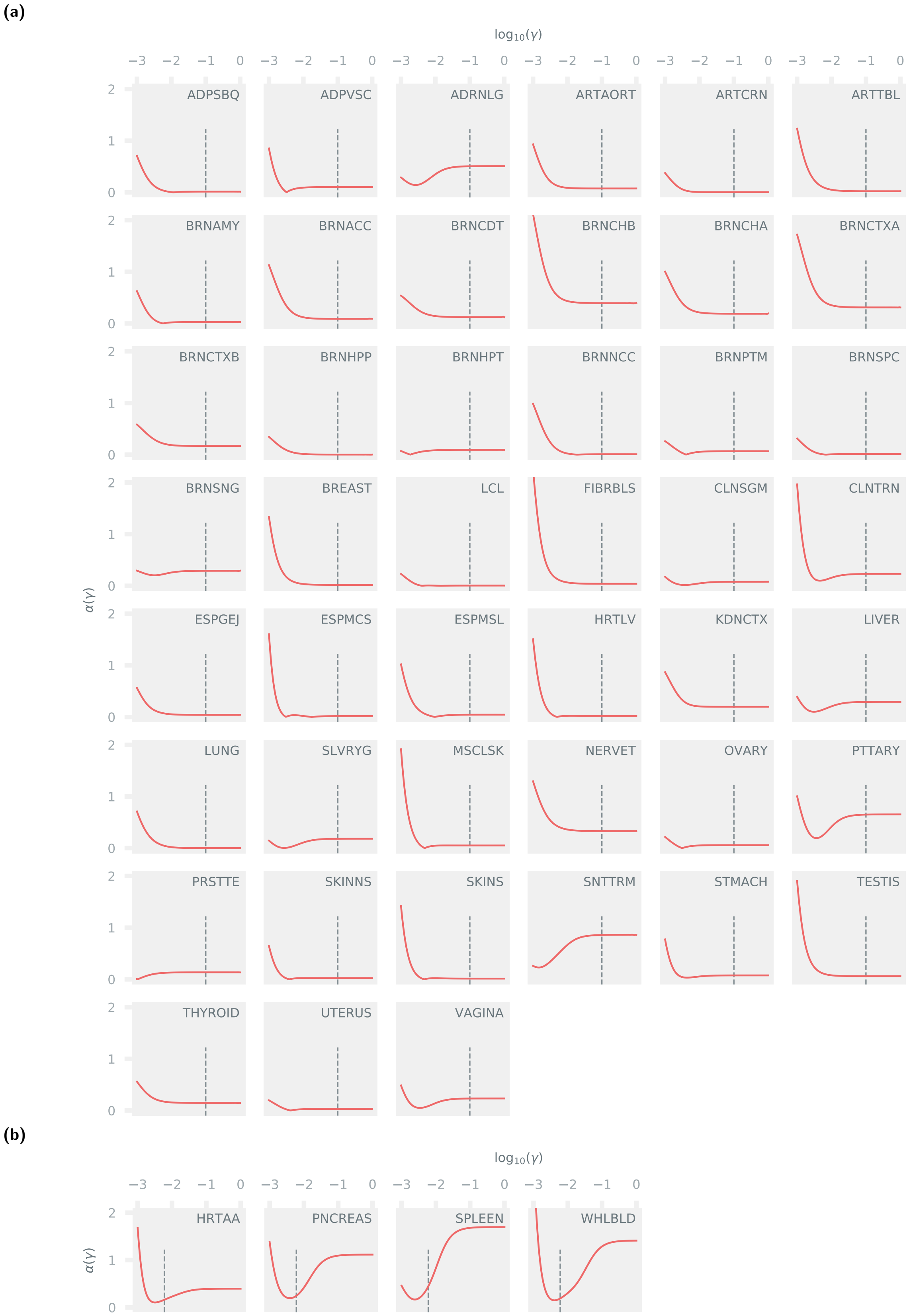

**Fig. S10 | Non-Gaussian parameters for all GTEx tissues.** We calculated the non-Gaussian parameters at different values of $\gamma$ for all tissues in GTEx. We used the preprocessed gene expression from the GTEx v8 data and used simulated genotype to calculate the $\alpha(\gamma)$ for each tissue. The solid red line in each panel shows the variation of $\alpha(\gamma)$ for a given tissue with $-\log_{10}(\gamma)$. We divided the tissues into two groups: **(a)** for these tissues, we chose $\gamma$ = 0.1, and **(b)** for these tissues, we chose $\gamma$ = 0.006.

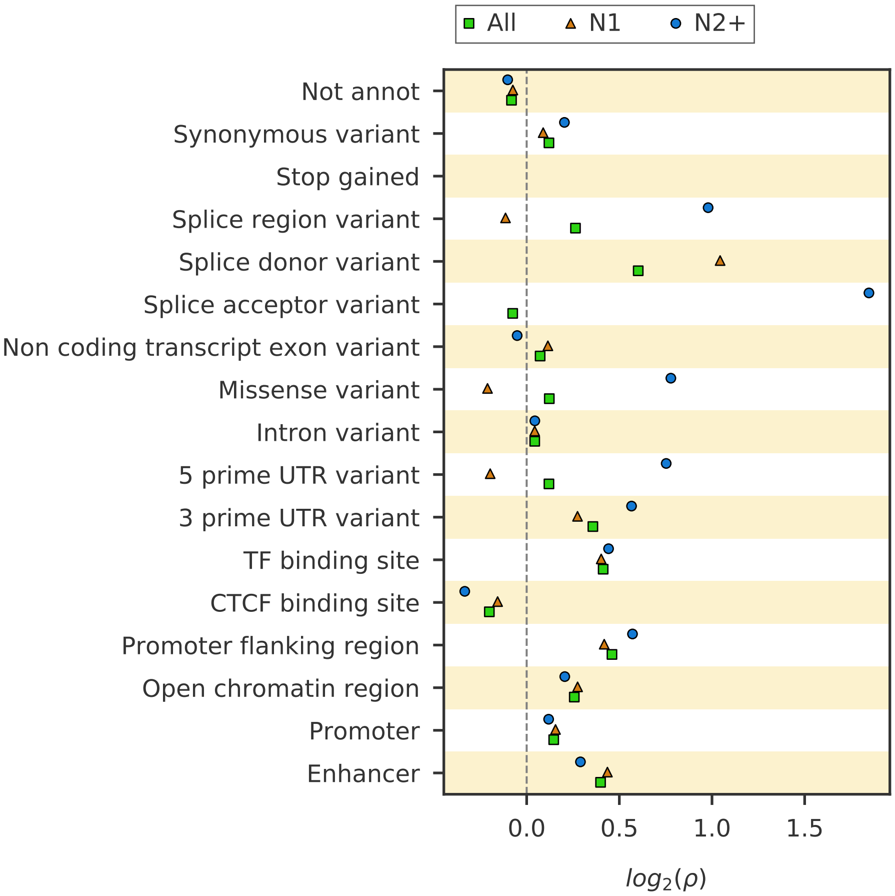

**Fig. S11 | Trans-eQTLs are enriched in known regulatory elements.** Enrichment in genomic functional annotations can give us insights into how trans-eQTLs affect gene expression. We considered three sets of trans-eQTLs: (i) the set of all trans-eQTLs ('All'), (ii) trans-eQTLs that are found in only one tissue ('N1') and (iii) trans-eQTLs that are found in two or more tissues ('N2+'). Genomic functional annotations were obtained from the GTEx portal [12].

#### 5.8 Effect of cis-masking

We applied Tejaas on 38 GTEx tissues (except brain tissues and bladder) with and without cis-masking (Sec. 2.9) to check the effect of cis-masking option of Tejaas for discovering trans-eQTLs in the GTEx data. We found that $\sim$ 97% of the trans-eQTLs discovered with cis-masking (shown with green in Fig. S12a) are also discovered without cis-masking (shown with red in Fig. S12a). However, if no cis-masking is used, Tejaas discovers $\sim$ 11% more trans-eQTLs compared to that with cis-masking. This increase in significant trans-eQTLs might be due to strong cis-eQTLs, as envisioned in Sec. 2.9.

We can understand the effect of cis-masking by looking at how many of our trans-eQTLs are also reported as cis-eQTLs by the GTEx Consortium [11]. We defined the proportion of cis-eQTLs among the trans-eQTLs compared to the proportion expected by chance as cis-eQTL enrichment ($\rho_{\text{cis-eQTL}}$). In Fig. S12b, we compare the $\log_2(\rho_{\text{cis-eQTL}})$ for trans-eQTLs discovered with and without cis-masking in all the 38 tissues. The mean $\log_2$ enrichment of cis-eQTLs across all tissues (shown in inset of Fig. S12b) is 0.10 with cis-masking and increases to 0.77 without cis-masking. This increase comes from the extra $\sim$ 11% SNPs discovered without cis-masking, indicating that cis-masking is crucial to avoid strong cis-eQTLs being falsely reported as trans-eQTLs by Tejaas.

**(a)**

**(b)**

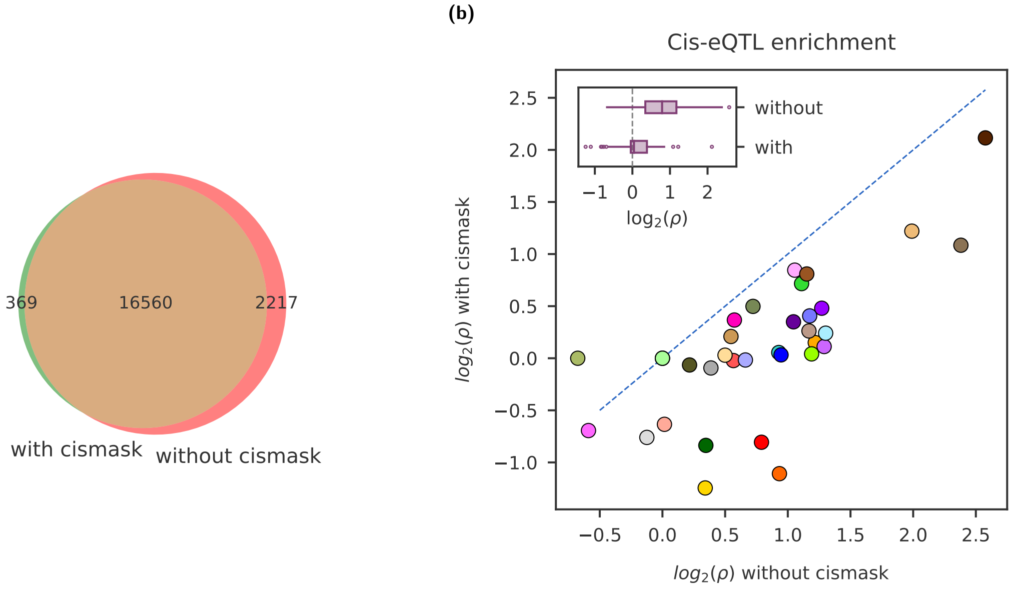

**Fig. S12 | Cis-masking controls false discovery of cis-eQTLs.** Cis-eQTLs with strong association with cis-eGenes may lead them to be discovered as trans-eQTLs by Tejaas. Exclusion of cis-genes greatly reduces the discovered cis-eQTLs while having minimal impact on the discovery of trans-eQTLs. **(a)** Overlap of trans-eQTLs with and without cis-masking. 97% of trans-eQTLs, which are found with cis-masking are also discovered without cismasking. **(b)** Comparison of cis-eQTL enrichment with and without cis-masking. Cis-eQTL enrichment reduces significantly in all tissues when cis-masking is enabled, indicating that the extra trans-eQTLs discovered without cis-masking are mostly cis-eQTLs.

#### 5.9 Effect of cross-mappable genes

False trans-eQTL signals could arise from multi-mapped reads within genes with high sequence similarity, where reads from a given gene are mapped to one or more other genes elsewhere in the genome and showing an association resembling a long range regulatory effect. This problem was explored by Saha and Battle [13] and can be mitigated using cross-mappability scores. We obtained pre-computed cross-mappability scores from [13], with settings of k-mer length 74 for exons and 36 for UTRs. In Tejaas we extended the cis-masking approach to exclude all genes that cross-map with any of the cis-genes listed in a given cis-masking group and we call this the crossmap filter. In the GTEx non-brain tissues, we discovered a total of 19556 significant trans-eQTLs using the crossmap filter, compared to 16929 trans-eQTLs without the cross-mappability filter. 13438 trans-eQTLs were found in common in both sets. Enrichment of trans-eQTLs in DHS regions and cis-eQTLs does not change significantly ($p > 0.1$) with and without the cross-mappability filter. (Fig. S13).

#### 5.10 Effect of GTEx populations in predicted trans-eQTLs

To assess the effect of population differentiation in GTEx, we calculated fixation index values ($F_{ST}$ values) as in [14] on GTEx genotype data. We compared the distribution of $F_{ST}$ values for all GTEx SNPs with the set of trans-eQTLs predicted with Tejaas (Fig. S14, left panel). The average $F_{ST}$ value for the predicted trans-eQTLs in each tissue is shown on the right panel of Fig. S14. High $F_{ST}$ values are indicative of allele frequency differences between subpopulations present in the data. Population substructure could give spurious indications of eQTL associations [15]. To rule out confounding effects of population differentiation from our enrichment analysis, we sampled null sets with the same $F_{ST}$ distribution as that of the predicted trans-eQTLs. We re-calculated

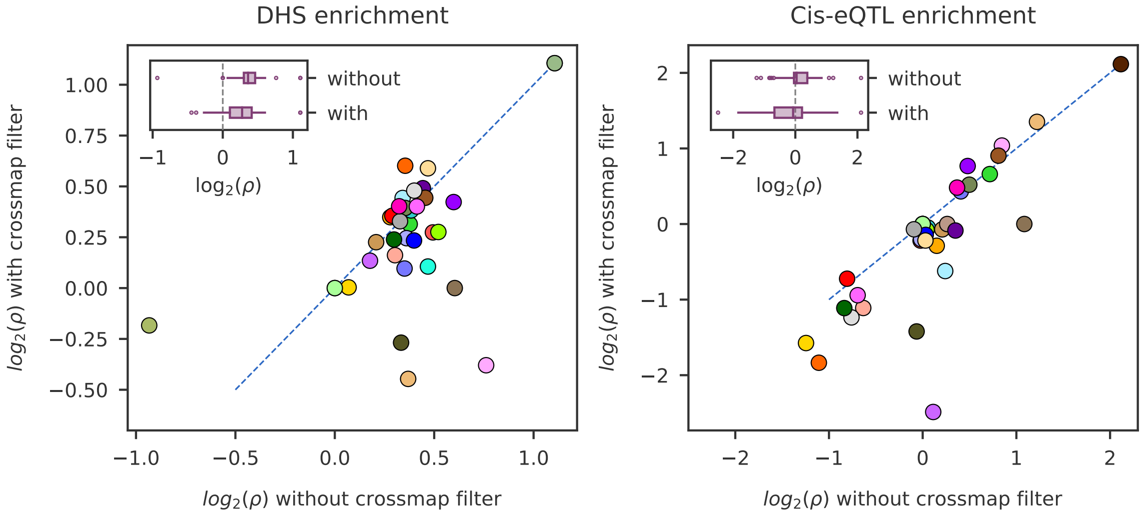

**Fig. S13 | Effect of cross-mappable genes.** Comparison of DHS enrichment and cis-eQTL enrichment with and without cross-mappable genes show that there is no significant difference. A number of GTEx tissues show lower enrichments when the cross-mappable filter is applied.

DHS functional enrichments (Fig. S15) and GWAS enrichemnts (data not shown). However, we did not observe any significant deviation from the previously calculated enrichments.

#### 5.11 Regulatory element enrichment in brain tissues

Brain tissues in GTEx have generally lower number of RNA-seq samples compared to other tissues. This is one of the reasons why they display lower number of predicted trans-eQTLs with Tejaas. Other unknown confounders in the expression measurements could also affect Tejaas predictions, as it has been reported that brain tissues have markedly distinct expression signatures in GTEx that separates them from the rest of the tissues. This could be due to the post mortem nature of the samples. We report DHS, raQTL and functional region enrichments for trans-eQTLs predicted in brain tissues in Fig. S16. Due to the low number of predictions, the reported enrichments are not reliable.

#### 5.12 Performance of Tejaas on null data

On null data, Tejaas does not show any spurious association (Fig. S17). We used the same gene expression as used for trans-eQTL discovery to check if the correlation of the gene expression can lead to false discovery. We removed any possible trans-eQTL signal by permuting the sample labels of the genotype. We found no significant association in any GTEx tissue.

#### 5.13 Replication in eQTLGen

We downloaded the list of significant trans-eQTLs discovered in the eQTLGen study from `https://www.eqtlgen.org/trans-eqtls.html` (version from 2018-09-04). It contains 59 786 SNP-gene pairs and a total of 3 853 unique trans-eQTLs in whole blood. In Appendix 3, we report the overlap between trans-eQTL from eQTLGen with the lead trans-eQTLs discovered with Tejaas in all GTEx tissues. Enrichments were calculated as in Sec. 5.6.

Replication across all GTEx tissues is low, with whole blood showing the highest number of replicated variants. Although we don't expect to replicate other studies that used single SNP-gene pairs association methods, it is reassuring to see that Tejaas can replicate trans-eQTLs discovered

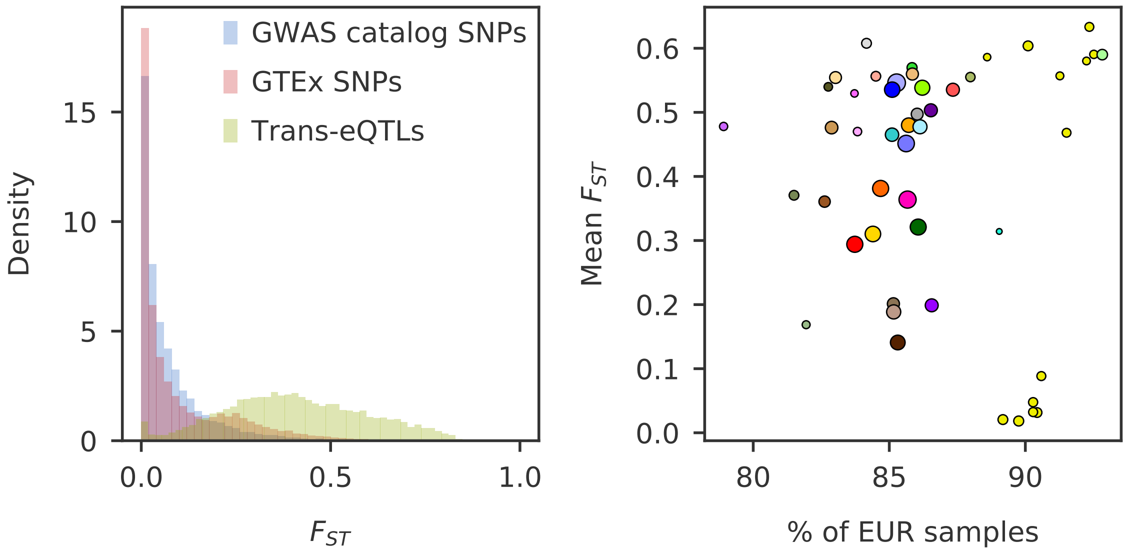

**Fig. S14 | Distribution of $F_{ST}$ values in GTEx and predicted trans-eQTLs.** Left panel: Distribution of $F_{ST}$ values for all the GTEx SNPs analyzed (red), all SNPs present in GTEx and in the GWAS catalog (blue) and the trans-eQTLs predicted with Tejaas (green). Area under the curve is normalized to 1. Right panel: Mean $F_{ST}$ values of predicted trans-eQTLs in each tissue, colored in GTEx colors. Size of the dot indicates number of samples.

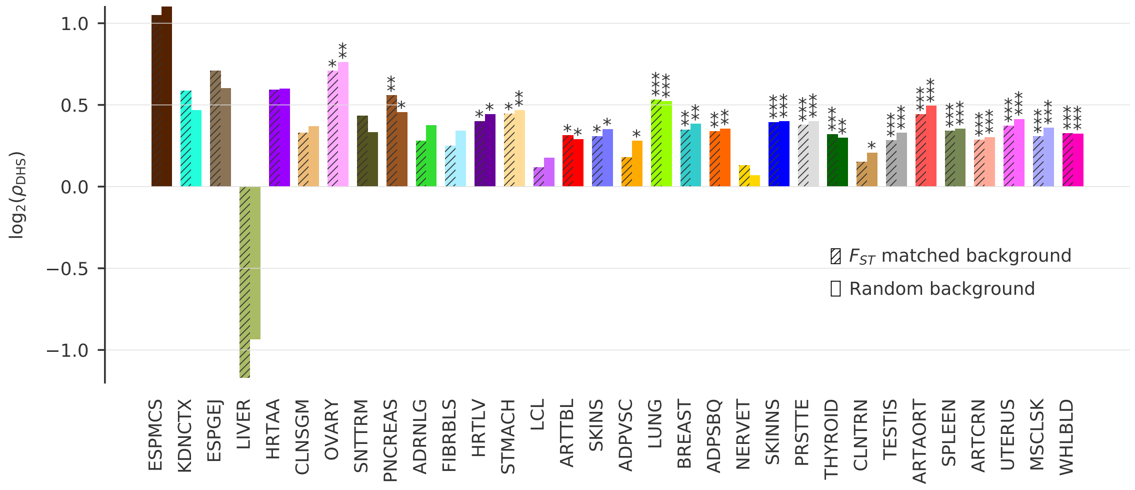

**Fig. S15 | Trans-eQTL DHS enrichment after accounting for $F_{ST}$.** We account for the enrichment of predicted trans-eQTLs in $F_{ST}$ values by adjusting the background enrichment to have the same $F_{ST}$ distribution as the one for predicted trans-eQTLs in each tissue.

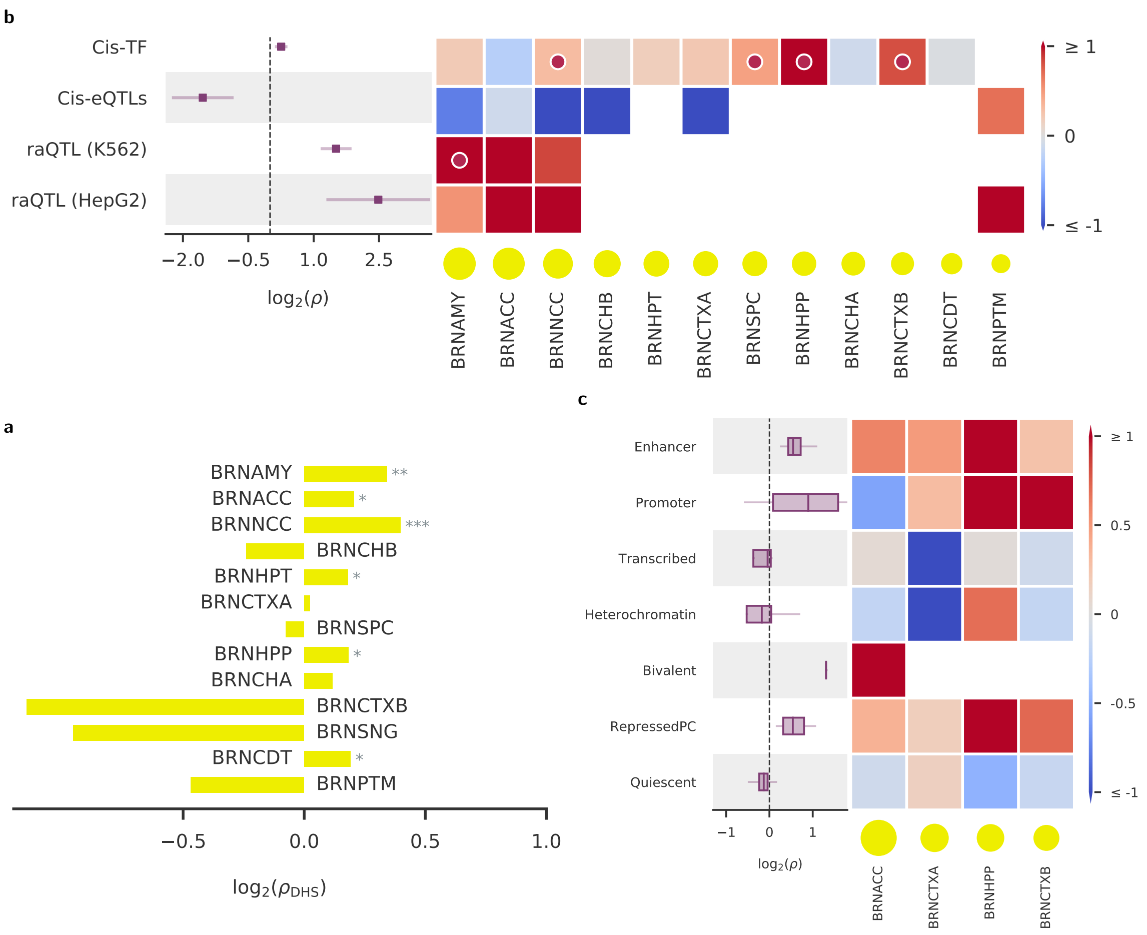

**Fig. S16 | Functional enrichments in brain tissues. a**, $Log_2$ enrichments (x-axis) within accessible chromatin regions from [16]. **b**, $Log_2$ enrichments near known eQTLs, Transcription Factors and reporter assay QTLs (raQTLs). **c**, $Log_2$ enrichments within tissue-specific regulatory regions. For more details, see caption of Fig. 04 in main text.

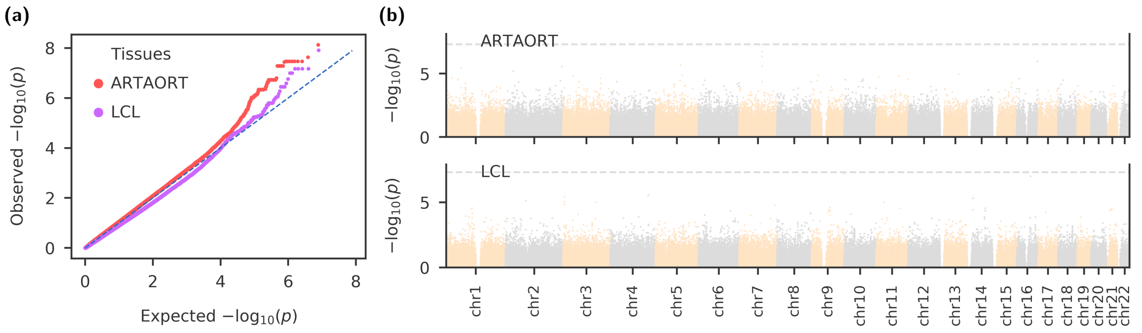

**Fig. S17 | Performance of Tejaas on null data.** We removed possible trans-eQTL signal from the GTEx data by permuting the sample labels of the genotype. We did not permute the gene expression of the tissues to retain the inherent correlation among the genes. **(a)** Representative example of quantile-quantile plot from null data in two tissues: artery aorta (ARTAORT) and EBV-transformed lymphocytes (LCL). **(b)** Corresponding Manhattan plot for the above two tissues, showing the $-\log_{10}(p)$ values for genome-wide variants.

in the same tissue type. Nonetheless, we must highlight that the eQTLGen meta-analysis study includes GTEx whole blood expression.

### 6 GWAS analysis

We investigated the overlap of the novel trans-eQTLs discovered by Tejaas with GWAS variants of complex traits to find transcriptional regulatory mechanisms through which SNPs affect complex diseases. In this section, we discuss the details of our analyses and report supplementary results.

#### 6.1 Data source for GWAS summary statistics

The GWAS Catalog was obtained from the EBI website [17]. We used version 'e98_r2020-03-08' which contains 4493 studies with a total of 179 364 associations [18, 19]. The set of 87 GWAS studies was harmonized and imputed to GTEx v8 variants with MAF $> 0.01$ using only European samples by Barbeira *et al.* [20]. These 87 traits were broadly classified into 11 disease categories.

#### 6.2 Calculation of GWAS enrichment

For every GWAS, we used 5000 randomly chosen SNPs from the GTEx genotype to calculate the fraction of SNPs that overlap with GWAS SNPs expected by chance,

$$f_{\text{bg}} = \frac{\text{number of random SNPs overlapping with GWAS SNPs}}{5000} \tag{51}$$

This is repeated 300 times and the mean $\langle f_{\text{bg}} \rangle$ is considered as the background for that GWAS. We then calculated the fraction of trans-eQTLs observed in the GWAS,

$$f_{\text{trans-eqtl}} = \frac{\text{number of trans-eQTLs overlapping with GWAS SNPs}}{\text{number of trans-eQTLs in the tissue}} \tag{52}$$

and the GWAS enrichment of trans-eQTLs in that tissue,

$$\rho_{\text{GWAS}} = \frac{f_{\text{trans-eqtl}}}{\langle f_{\text{bg}} \rangle} \tag{53}$$

We used a binomial test to calculate the $p$-values for the enrichment of each tissue-GWAS pair. If $T$ is the number of trans-eQTLs in the tissue, then the probability of finding $k$ of them in a GWAS is given by,

$$P\left(x = k\right) = \text{Binomial}\left(T, k, \langle f_{\text{bg}} \rangle\right) \; . \tag{54}$$

and $P\left(x > k\right)$ gives us the $p$-value for the tissue-GWAS pair.

For category-wise enrichment, we checked the overlap of trans-eQTLs with all disease-associated SNPs in that category. For global enrichment, we checked the overlap of trans-eQTLs with all disease-associated SNPs in the dataset.

#### 6.3 Result

For the 87 GWAS studies, we show the GWAS enrichment for every tissue-study pair in Fig. S18. The majority of the trans-eQTLs are tissue-specific. We find them to be enriched only in a group of traits related to the same tissue. Six tissues that affect a range of traits across different disease categories show global enrichment. Two good examples are thyroid and whole-blood. The trans-eQTLs in thyroid are significantly enriched in hypothyroidism with $\rho = 3.58$ and $p = 0.014$.

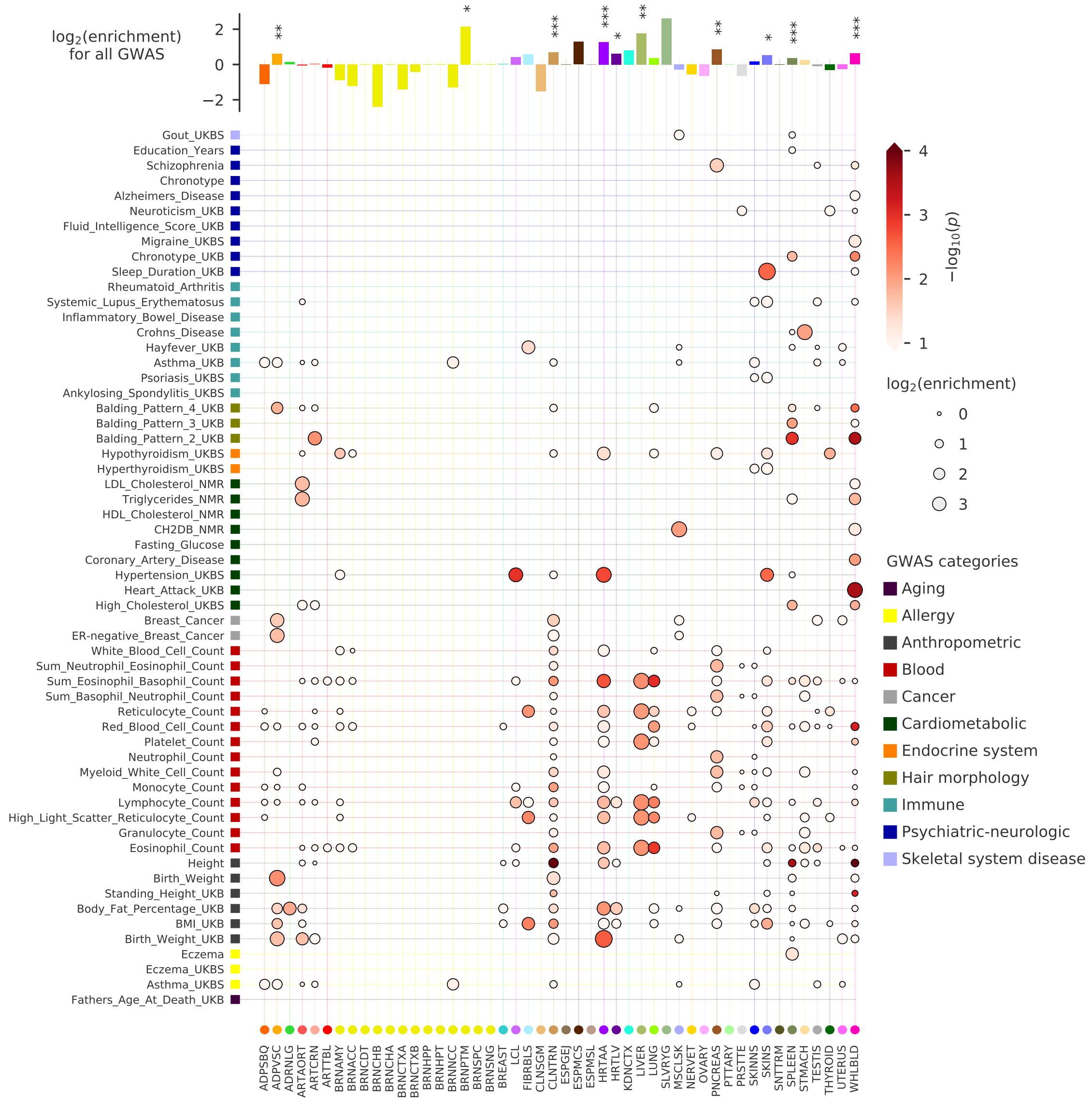

**Fig. S18 | Trans-eQTLs overlap with GWAS SNPs for 87 complex diseases.** We calculated the enrichment of lead trans-eQTLs predicted by Tejaas across all tissues (x-axis) in GWAS hits of 87 GWAS (y-axis). Each point in the plot represents the enrichment of trans-eQTLs of a tissue in a GWAS, the size of the point scales with the $\log_2$ enrichment, and the color scales with the significance ($-\log_{10}(p)$) of the enrichment. In the top panel, we show the GWAS enrichment for all studies combined.

However, they do not show global enrichment (top panel Fig. S18). Whole blood is enriched in a wide range of studies across multiple study categories, such as 'Blood' ($\rho = 1.3$, $p = 0.0014$), 'Cardiometabolic' ($\rho = 1.66$, $p = 0.01$) and 'Psychiatric-neurologic' ($\rho = 1.4$, $p = 0.02$) among others. Hence, they also show a global enrichment.

Some of the results have a clear biological relationship. As mentioned in the manuscript, the enrichment of thyroid trans-eQTLs in hypothyroidism has a direct connection. Another interesting example is adipose visceral omentum which is the body's primary reserve for lipids, where trans-eQTLs show 3-fold enrichment in both body fat percentage ($\rho = 3.36$, $p = 0.033$) and BMI ($\rho = 3.17$, $p = 0.022$) studies. Other examples are those found in blood, with $\rho = 13.4$, $p = 0.003$ for heart attack and $\rho = 4$, $p = 0.009$ for coronary artery disease (CAD), while Heart Atrial Appendage tissue contains trans-eQTLs enriched in hypertension GWAS ($\rho = 13.1$, $p = 0.002$).

### Appendix 1. Expectation and variance of RR-score under the null hypothesis

Here we show the derivation of the expectation value and the variance of $q_{\text{rev}}$ (Eq. 19) under the permutation null model for any symmetric matrix $\mathbf{W}$ (Eq. 20) and any centered vector $\mathbf{x}$. We rewrite $q_{\text{rev}}$ as,

$$q_{\text{rev}} = \sum_{n,m=1}^{N} W_{nm} x_n x_m \,. \tag{55}$$

#### Expectation value of RR-score

We denote the permutation function of the indices between 1 and $N$ as $\pi(\cdot)$ and we write $\mathbb{E}_\pi[\ldots]$ for the expectation value over multiple independent permutations. For example $\mathbb{E}_\pi\left[x_{\pi(n)}\right] = \sum_m x_m/N = 0$ where $x_{\pi(n)}$ is the $n^{\text{th}}$ entry of the vector $\mathbf{x}$ under permutation $\pi(\mathbf{x})$. For the expectation value of $q_{\text{rev}}$ under random permutations we get

$$\mathbb{E}_\pi\left[\sum_{n,m=1}^{N} W_{nm} x_{\pi(n)} x_{\pi(m)}\right] = \sum_{n,m=1}^{N} W_{nm}\,\mathbb{E}_\pi\left[x_{\pi(n)} x_{\pi(m)}\right] \tag{56}$$

Their are only two values adopted by the expectation value $\mathbb{E}_\pi\left[x_{\pi(n)} x_{\pi(m)}\right]$, one in the case $n = m$ and the other in the case $n \neq m$. The first value is just $\mathbb{E}_\pi\left[x^2_{\pi(n)}\right] = (1/N)\sum_{n=1}^{N} x_n^2$, which is easily computed from $\mathbf{x}$. The value for the case $n \neq m$ is more tricky:

$$\begin{aligned}
\mathbb{E}_\pi\left[x_{\pi(n)} x_{\pi(m)}\right] &= \mathbb{E}_{n\neq m}\left[x_n x_m\right] \\
&= \frac{1}{N(N-1)} \sum_{n,m:n\neq m}^{N} x_n x_m \\
&= \frac{1}{N(N-1)} \left(\sum_{n,m}^{N} x_n x_m - \sum_{n=1}^{N} x_n^2\right) \\
&= \frac{1}{N(N-1)} \left(\left(\cancel{\sum_{n=1}^{N} x_n}\right)^2 - \sum_{n=1}^{N} x_n^2\right),
\end{aligned} \tag{57}$$

where the sum over $x_n$ cancels out due to the centering of $\mathbf{x}$. We define the following moments of $\mathbf{x}$:

$$\begin{aligned}
\mu_2 &:= \frac{1}{N} \sum_{n=1}^{N} x_n^2 \\
\mu_4 &:= \frac{1}{N} \sum_{n=1}^{N} x_n^4
\end{aligned} \tag{58}$$

with which the expectation value for $n \neq m$ becomes

$$\mathbb{E}_\pi\left[x_{\pi(n)} x_{\pi(m)}\right] = -\frac{\mu_2}{N-1} \tag{59}$$

Inserting this into equation Eq. (56) yields

$$\mathbb{E}_\pi\left[\sum_{n,m=1}^{N} W_{nm}\, x_{\pi(n)}x_{\pi(m)}\right] = -\frac{\mu_2}{N-1}\sum_{n,m:n\neq m}^{N} W_{nm} + \mu_2\sum_{n=1}^{N} W_{nn} \tag{60}$$

The first sum over $W_{nm}$ can be expressed in terms of simpler sums,

$$\sum_{n,m:n\neq m}^{N} W_{nm} = \sum_{n,m=1}^{N} W_{nm} - \sum_{n=1}^{N} W_{nn} = w_{11} - w_2, \tag{61}$$

where we have defined the following moments of $\mathbf{W}$,

$$w_{11} := \sum_{n,m=1}^{N} W_{nm} \tag{62}$$

$$w_2 := \sum_{n=1}^{N} W_{nn}. \tag{63}$$

Inserting this into Eq. (60) yields

$$\langle q_{\text{rev}}\rangle = \mathbb{E}_\pi\left[\sum_{n,m=1}^{N} W_{nm}\, x_{\pi(n)}x_{\pi(m)}\right] = \frac{\mu_2}{N-1}\left(Nw_2 - w_{11}\right)\ . \tag{64}$$

#### Variance of RR-score

We will determine the variance with the equation

$$\text{Var}\left[q_{\text{rev}}\right] = \mathbb{E}_\pi\left[\left(\sum_{n,m=1}^{N} W_{nm}\, x_{\pi(n)}x_{\pi(m)}\right)^2\right] - \mathbb{E}_\pi\left[\sum_{n,m=1}^{N} W_{nm}\, x_{\pi(n)}x_{\pi(m)}\right]^2, \tag{65}$$

and we therefore need to calculate the first term on the right hand side:

$$\mathbb{E}_\pi\left[\left(\sum_{n,m=1}^{N} W_{nm}\, x_{\pi(n)}x_{\pi(m)}\right)^2\right] = \sum_{n,m,n',m'=1}^{N} W_{nm}W_{n'm'}\,\mathbb{E}_\pi\left[x_{\pi(n)}x_{\pi(m)}x_{\pi(n')}x_{\pi(m')}\right]. \tag{66}$$

The term $\mathbb{E}_\pi\left[x_{\pi(n)}x_{\pi(m)}x_{\pi(n')}x_{\pi(m')}\right]$ can only take five different values, depending on how many of the four indices are identical to each other. To simplify the following, quite long derivation of the variance of $q_{\text{rev}}$, we will introduce some notation. First, to simplify writing down sums over two, three or four mutually different index variables, we define sets

$$\begin{aligned}
\mathcal{N}_2 &:= \{(n,m) : 1 \le n,m \le N, n\neq m\}\\
\mathcal{N}_3 &:= \{(n,m,n') : 1 \le n,m,n' \le N, n\neq m, n\neq n', m\neq m'\}\\
\mathcal{N}_4 &:= \{(n,m,n',m') : 1 \le n,m,n',m' \le N, n\neq m, n\neq n',\dots,n'\neq m'\}\,.
\end{aligned} \tag{67}$$

We denote by $\mathbb{E}_{(n.m)\in\mathcal{N}_2}\left[\cdot\right]$ or simply $\mathbb{E}_{\mathcal{N}_2}\left[\cdot\right]$ the expectation value over a random variable $(n,m)\in\mathcal{N}_2$, and analogously for $\mathbb{E}_{\mathcal{N}_3}\left[\cdot\right]$ and $\mathbb{E}_{\mathcal{N}_4}\left[\cdot\right]$. We further abbreviate

$$\begin{aligned}
v_{1111} &= \mathbb{E}_{\mathcal{N}_4}\left[x_n x_m x_{n'} x_{m'}\right]\\
v_{211} &= \mathbb{E}_{\mathcal{N}_3}\left[x_n^2 x_m x_{n'}\right]
\end{aligned}$$

$$
\begin{aligned}
\nu_{22} &= \mathbb{E}_{\mathcal{N}_2}\left[x_n^2 x_m^2\right] \\
\nu_{31} &= \mathbb{E}_{\mathcal{N}_2}\left[x_n^3 x_m\right].
\end{aligned} \tag{68}
$$

With these definitions, Eq. (66) becomes

$$
\begin{aligned}
&\mathbb{E}_\pi\left[\left(\sum_{n,m=1}^{N} W_{nm}\, x_{\pi(n)} x_{\pi(m)}\right)^2\right] \\
&= \nu_{1111} \sum_{(n,m,n',m')\in\mathcal{N}_4} W_{nm} W_{n'm'} \\
&+ \nu_{211} \sum_{(n,m,n')\in\mathcal{N}_3} \left(W_{nn}W_{mn'} + W_{nm}W_{n'n'} + W_{nm}W_{nn'} + W_{nm}W_{n'n} + W_{nm}W_{mn'} + W_{nm}W_{n'm}\right) \\
&+ \nu_{22} \sum_{(n,m)\in\mathcal{N}_2} \left(W_{nn}W_{mm} + W_{nm}W_{nm} + W_{nm}W_{mn}\right) \\
&+ \nu_{31} \sum_{(n,m)\in\mathcal{N}_2} \left(W_{nn}W_{nm} + W_{nn}W_{mn} + W_{nm}W_{nn} + W_{nm}W_{mm}\right) \\
&+ \mu_4 \sum_{n=1}^{N} W_{nn}^2
\end{aligned} \tag{69}
$$

and, using the symmetry of $\mathbf{W}$,

$$
\begin{aligned}
\mathbb{E}_\pi\left[\left(\sum_{n,m=1}^{N} W_{nm}\, x_{\pi(n)} x_{\pi(m)}\right)^2\right] &= \nu_{1111} \sum_{(n,m,n',m')\in\mathcal{N}_4} W_{nm} W_{n'm'} \\
&+ \nu_{211} \sum_{(n,m,n')\in\mathcal{N}_3} \left(2W_{nn}W_{mn'} + 4W_{nm}W_{nn'}\right) \\
&+ \nu_{22} \sum_{(n,m)\in\mathcal{N}_2} \left(W_{nn}W_{mm} + 2W_{nm}W_{nm}\right) \\
&+ \nu_{31} \sum_{(n,m)\in\mathcal{N}_2} 4W_{nn}W_{nm} + \mu_4 \sum_{n=1}^{N} W_{nn}^2
\end{aligned} \tag{70}
$$

We will first derive expressions for the $\nu$ terms and then for the sums of elements of $\mathbf{W}$.

$$
\begin{aligned}
\nu_{31} &= \mathbb{E}_{\mathcal{N}_2}\left[x_n^3 x_m\right] \\
&= \frac{1}{N(N-1)} \sum_{(n,m)\in\mathcal{N}_2} x_n^3 x_m \\
&= \frac{1}{N(N-1)} \left(\cancel{\sum_{n,m=1}^{N} x_n^3 x_m} - \sum_{n=1}^{N} x_n^4\right) \\
&= -\frac{1}{N-1}\mu_4
\end{aligned} \tag{71}
$$

$$
\begin{aligned}
\nu_{22} &= \mathbb{E}_{\mathcal{N}_2}\left[x_n^2 x_m^2\right] \\
&= \frac{1}{N(N-1)} \sum_{(n,m)\in\mathcal{N}_2} x_n^2 x_m^2
\end{aligned}
$$

$$
\begin{aligned}
&= \frac{1}{N(N-1)} \left( \sum_{n,m=1}^{N} x_n^2 x_m^2 - \sum_{n=1}^{N} x_n^4 \right) \\
&= \frac{N}{N-1}\mu_2^2 - \frac{1}{N-1}\mu_4
\end{aligned} \tag{72}
$$

$$
\begin{aligned}
v_{211} &= \mathbb{E}_{\mathcal{N}_3}\left[x_n^2 x_m x_{n'}\right] \\
&= \frac{1}{N(N-1)(N-2)} \sum_{(n,m,n')\in\mathcal{N}_3} x_n^2 x_m x_{n'} \\
&= \frac{1}{N(N-1)(N-2)} \sum_{(n,m)\in\mathcal{N}_2} \left( \cancel{\sum_{n'=1}^{N} x_n^2 x_m x_{n'}} - x_n^3 x_m - x_n^2 x_m^2 \right) \\
&= -\frac{1}{N-2}\left(v_{31} + v_{22}\right) \\
&= -\frac{1}{(N-1)(N-2)}\left(N\mu_2^2 - 2\mu_4\right)
\end{aligned} \tag{73}
$$

$$
\begin{aligned}
v_{1111} &= \mathbb{E}_{\mathcal{N}_3}\left[x_n x_m x_{n'} x_{m'}\right] \\
&= \frac{1}{N(N-1)(N-2)(N-3)} \sum_{(n,m,n',m')\in\mathcal{N}_4} x_n x_m x_{n'} x_{m'} \\
&= \frac{1}{N(N-1)(N-2)(N-3)} \sum_{(n,m,n')\in\mathcal{N}_3} \left( \cancel{\sum_{m'=1}^{N} x_n x_m x_{n'} x_{m'}} - x_n^2 x_m x_{n'} - x_n x_m^2 x_{n'} - x_n x_m x_{n'}^2 \right) \\
&= -\frac{3}{N-3}\, v_{211} \\
&= \frac{3}{(N-1)(N-2)(N-3)}\left(N\mu_2^2 - 2\mu_4\right)
\end{aligned} \tag{74}
$$

and, in summary,

$$
\begin{aligned}
v_{31} &= -\frac{1}{N-1}\mu_4 \\
v_{22} &= \frac{N}{N-1}\mu_2^2 + v_{31} \\
v_{211} &= -\frac{1}{N-2}\left(v_{31} + v_{22}\right) \\
v_{1111} &= -\frac{3}{N-3}\, v_{211}\,.
\end{aligned} \tag{75}
$$

We will now derive expressions for the sums of elements of $\mathbf{W}$ in equation (70):

$$
\sum_{(n,m)\in\mathcal{N}_2} W_{nn} W_{nm} = \sum_{n,m=1}^{N} W_{nn} W_{nm} - \sum_{n=1}^{N} W_{nn}^2 = w_{31} - w_4 \tag{76}
$$

where

$$
\begin{aligned}
w_{31} &:= \sum_{n,m=1}^{N} W_{nn} W_{nm} = \sum_{n=1}^{N} W_{nn} W_{n}. \\
w_4 &:= \sum_{n=1}^{N} W_{nn}^2
\end{aligned}
$$

$$w_{22} := \sum_{n,m=1}^{N} W_{nm}^2$$

$$w_{211} := \sum_{n,m,n'=1}^{N} W_{nm} W_{nn'} = \sum_{n=1}^{N} \left( \sum_{m=1}^{N} W_{nm} \right)^2 = \sum_{n=1}^{N} W_n^2. \tag{77}$$

Also

$$\sum_{(n,m)\in\mathcal{N}_2} W_{nn} W_{mm} = \sum_{n,m=1}^{N} W_{nn} W_{mm} - \sum_{n=1}^{N} W_{nn}^2 = w_2^2 - w_4 \tag{78}$$

$$\sum_{(n,m)\in\mathcal{N}_2} W_{nm} W_{nm} = \sum_{n,m=1}^{N} W_{nm} W_{nm} - \sum_{n=1}^{N} W_{nn}^2 = w_{22} - w_4 \tag{79}$$

$$\begin{aligned}
\sum_{(n,m,n')\in\mathcal{N}_3} W_{nn} W_{mn'} &= \sum_{(n,m)\in\mathcal{N}_2} \sum_{n'=1}^{N} W_{nn} W_{mn'} - \sum_{(n,m)\in\mathcal{N}_2} (W_{nn} W_{mn} + W_{nn} W_{mm}) \\
&= \sum_{(n,m)\in\mathcal{N}_2} \sum_{n'=1}^{N} W_{nn} W_{mn'} - \sum_{n,m=1}^{N} (W_{nn} W_{mn} + W_{nn} W_{mm}) + 2w_4 \\
&= \sum_{n,m,n'=1}^{N} W_{nn} W_{mn'} - \sum_{n,n'=1}^{N} W_{nn} W_{nn'} - w_{31} - \sum_{n=1}^{N} W_{nn} \sum_{m=1}^{N} W_{mm} + 2w_4 \\
&= \sum_{n=1}^{N} W_{nn} \sum_{m,n'=1}^{N} W_{mn'} - w_{31} - w_{31} - w_2^2 + 2w_4 \\
&= w_2\, w_{11} - 2w_{31} - w_2^2 + 2w_4
\end{aligned} \tag{80}$$

$$\begin{aligned}
\sum_{(n,m,n')\in\mathcal{N}_3} W_{nm} W_{nn'} &= \sum_{(n,m)\in\mathcal{N}_2} \sum_{n'=1}^{N} W_{nm} W_{nn'} - \sum_{n,m=1}^{N} (W_{nm} W_{nm} + W_{nm} W_{nn}) + 2w_4 \\
&= \sum_{n,m=1}^{N} \sum_{n'=1}^{N} W_{nm} W_{nn'} - \sum_{n=1}^{N} \sum_{n'=1}^{N} W_{nn} W_{nn'} - w_{22} - w_{31} + 2w_4 \\
&= (w_{211} - w_{31}) - w_{22} - w_{31} + 2w_4 \\
&= w_{211} - 2w_{31} - w_{22} + 2w_4
\end{aligned} \tag{81}$$

$$\begin{aligned}
\sum_{(n,m,n',m')\in\mathcal{N}_4} W_{nm} W_{n'm'} &= \sum_{(n,m,n')\in\mathcal{N}_3} \sum_{m'=1}^{N} W_{nm} W_{n'm'} - \sum_{(n,m,n')\in\mathcal{N}_3} (W_{nm} W_{n'n} + W_{nm} W_{n'm} + W_{nm} W_{n'n'}) \\
&= \sum_{(n,m,n')\in\mathcal{N}_3} \sum_{m'=1}^{N} W_{nm} W_{n'm'} - 2 \sum_{(n,m,n')\in\mathcal{N}_3} W_{nm} W_{nn'} - \sum_{(n,m,n')\in\mathcal{N}_3} W_{nn} W_{mn'} \\
&\overset{(81)(80)}{=} \sum_{(n,m)\in\mathcal{N}_2} \left( \sum_{n',m'=1}^{N} W_{nm} W_{n'm'} - \sum_{m'=1}^{N} (W_{nm} W_{nm'} + W_{nm} W_{mm'}) \right) \\
&\quad - 2(w_{211} - 2w_{31} - w_{22} + 2w_4) - (w_2\, w_{11} - 2w_{31} - w_2^2 + 2w_4) \\
&\overset{(61)}{=} (w_{11} - w_2)\, w_{11} - 2 \sum_{(n,m)\in\mathcal{N}_2} \sum_{m'=1}^{N} W_{nm} W_{nm'} \\
&\quad - 2(w_{211} - 2w_{31} - w_{22} + 2w_4) - (w_2\, w_{11} - 2w_{31} - w_2^2 + 2w_4) \\
&\overset{(81)}{=} (w_{11} - w_2)\, w_{11} - 2(w_{211} - w_{31})
\end{aligned}$$

$$
\begin{aligned}
&- 2(w_{211} - 2w_{31} - w_{22} + 2w_4) - (w_2\, w_{11} - 2w_{31} - w_2^2 + 2w_4) \\
&= w_{11}^2 - 2w_2\, w_{11} - 4w_{211} + 8w_{31} + 2w_{22} + w_2^2 - 6w_4
\end{aligned} \tag{82}
$$

We can check that the last term is correct by checking if the number of $W_{nm}W_{n'm'}$ terms in the last line is equal to $N(N-1)(N-2)(N-3) = N^4 - 6N^3 + 11N^2 - 6N$. To do that, we check that each of the $w$ terms sums up $N^k$ terms, where $k$ is the number of digits in its subscript. For example $w_4$ and $w_2$ sum up $N$ terms, $w_{11}$ and $w_{22}$ and $w_{31}$ sum up $N^2$, and $w_{211}$ sums up $N^3$. The last line of the previous equation therefore sums up $N^4 - 2NN^2 - 4N^3 + 8N^2 + 2N^2 + N^2 - 6N = N^4 - 6N^3 + 11N^2 - 6N$ terms, which gives exactly what it should. Doing that with the other terms also checks. We insert the expressions for the sums over $\mathbf{W}$ terms into Eq. (70) and obtain

$$
\begin{aligned}
\mathbb{E}_\pi \left[ \left( \sum_{n,m=1}^{N} W_{nm}\, x_{\pi(n)} x_{\pi(m)} \right)^2 \right] &= \nu_{1111} \left( w_{11}^2 - 2w_2\, w_{11} - 4w_{211} + 8w_{31} + 2w_{22} + w_2^2 - 6w_4 \right) \\
&+ \nu_{211} \left( 2(w_2\, w_{11} - 2w_{31} - w_2^2 + 2w_4) + 4(w_{211} - 2w_{31} - w_{22} + 2w_4) \right) \\
&+ \nu_{22} \left( (w_2^2 - w_4) + 2(w_{22} - w_4) \right) \\
&+ 4\, \nu_{31}\, (w_{31} - w_4) + \mu_4 w_2
\end{aligned} \tag{83}
$$

and, by adding up terms with the same $w$ moments, finally

$$
\begin{aligned}
\mathbb{E}_\pi \left[ \left( \sum_{n,m=1}^{N} W_{nm}\, x_{\pi(n)} x_{\pi(m)} \right)^2 \right] &= \nu_{1111} \left( w_{11}^2 - 2w_2\, w_{11} - 4w_{211} + 8w_{31} + 2w_{22} + w_2^2 - 6w_4 \right) \\
&+ 2\, \nu_{211} \left( w_2\, w_{11} + 2w_{211} - 6w_{31} - 2w_{22} - w_2^2 + 6w_4 \right) \\
&+ \nu_{22} \left( w_2^2 + 2w_{22} - 3w_4 \right) + 4\, \nu_{31}\, (w_{31} - w_4) + \mu_4 w_4
\end{aligned} \tag{84}
$$

### Appendix 2. Abbreviation of GTEx tissues

| Tissue | Abbreviation |
|---|---|
| Adipose - Subcutaneous | ADPSBQ |
| Adipose - Visceral (Omentum) | ADPVSC |
| Adrenal Gland | ADRNLG |
| Artery - Aorta | ARTAORT |
| Artery - Coronary | ARTCRN |
| Artery - Tibial | ARTTBL |
| Brain - Amygdala | BRNAMY |
| Brain - Anterior cingulate cortex (BA24) | BRNACC |
| Brain - Caudate (basal ganglia) | BRNCDT |
| Brain - Cerebellar Hemisphere | BRNCHB |
| Brain - Cerebellum | BRNCHA |
| Brain - Cortex | BRNCTXA |
| Brain - Frontal Cortex (BA9) | BRNCTXB |
| Brain - Hippocampus | BRNHPP |
| Brain - Hypothalamus | BRNHPT |
| Brain - Nucleus accumbens (basal ganglia) | BRNNCC |
| Brain - Putamen (basal ganglia) | BRNPTM |
| Brain - Spinal cord (cervical c-1) | BRNSPC |
| Brain - Substantia nigra | BRNSNG |
| Breast - Mammary Tissue | BREAST |
| Cells - EBV-transformed lymphocytes | LCL |
| Colon - Sigmoid | CLNSGM |
| Colon - Transverse | CLNTRN |
| Esophagus - Gastroesophageal Junction | ESPGEJ |
| Esophagus - Mucosa | ESPMCS |
| Esophagus - Muscularis | ESPMSL |
| Heart - Atrial Appendage | HRTAA |
| Heart - Left Ventricle | HRTLV |
| Kidney - Cortex | KDNCTX |
| Liver | LIVER |
| Lung | LUNG |
| Minor Salivary Gland | SLVRYG |
| Muscle - Skeletal | MSCLSK |
| Nerve - Tibial | NERVET |
| Ovary | OVARY |
| Pancreas | PNCREAS |
| Pituitary | PTTARY |
| Prostate | PRSTTE |
| Skin - Not Sun Exposed (Suprapubic) | SKINNS |
| Skin - Sun Exposed (Lower leg) | SKINS |
| Small Intestine - Terminal Ileum | SNTTRM |
| Spleen | SPLEEN |
| Stomach | STMACH |
| Testis | TESTIS |
| Thyroid | THYROID |
| Uterus | UTERUS |
| Vagina | VAGINA |
| Whole Blood | WHLBLD |

### Appendix 3. eQTLGen Replication

| Tissue | Replicated trans-eQTLs | Enrichment |
|---|---|---|
| **Whole Blood** | **37 (0.96%)** | **3.16 ($p = 2.99012e^{-09}$)** |
| Adipose Subcutaneous | 0 (0.00%) | - |
| Adipose Visceral Omentum | 1 (0.03%) | 1.45 ($p$ = 0.498496) |
| Adrenal Gland | 3 (0.08%) | 8.33 ($p$ = 0.00593454) |
| Artery Aorta | 2 (0.05%) | 1.03 ($p$ = 0.577555) |
| Artery Coronary | 3 (0.08%) | 1.08 ($p$ = 0.5258) |
| Artery Tibial | 0 (0.00%) | - |
| Brain Amygdala | 0 (0.00%) | - |
| Brain Anterior cingulate cortex BA24 | 2 (0.05%) | 1.41 ($p$ = 0.415078) |
| Brain Caudate basal ganglia | 0 (0.00%) | - |
| Brain Cerebellar Hemisphere | 0 (0.00%) | - |
| Brain Cerebellum | 0 (0.00%) | - |
| Brain Cortex | 0 (0.00%) | - |
| Brain Frontal Cortex BA9 | 0 (0.00%) | - |
| Brain Hippocampus | 0 (0.00%) | - |
| Brain Hypothalamus | 0 (0.00%) | - |
| Brain Nucleus accumbens basal ganglia | 0 (0.00%) | - |
| Brain Putamen basal ganglia | 0 (0.00%) | - |
| Brain Spinal cord cervical c-1 | 0 (0.00%) | - |
| Brain Substantia nigra | 0 (0.00%) | - |
| Breast Mammary Tissue | 3 (0.08%) | 4.55 ($p$ = 0.0294283) |
| Cells EBV-transformed lymphocytes | 2 (0.05%) | 5.00 ($p$ = 0.061526) |
| Cells Cultured fibroblasts | 1 (0.03%) | 5.00 ($p$ = 0.181291) |
| Colon Sigmoid | 1 (0.03%) | 7.14 ($p$ = 0.130668) |
| Colon Transverse | 9 (0.23%) | 4.29 ($p$ = 0.00033596) |
| Esophagus Gastroesophageal Junction | 0 (0.00%) | - |
| Esophagus Mucosa | 0 (0.00%) | - |
| Esophagus Muscularis | 0 (0.00%) | - |
| Heart Atrial Appendage | 1 (0.03%) | 5.88 ($p$ = 0.156371) |
| Heart Left Ventricle | 2 (0.05%) | 7.14 ($p$ = 0.0325744) |
| Kidney Cortex | 0 (0.00%) | - |
| Liver | 0 (0.00%) | - |
| Lung | 2 (0.05%) | 2.82 ($p$ = 0.15927) |
| Minor Salivary Gland | 0 (0.00%) | - |
| Muscle Skeletal | 4 (0.10%) | 0.92 ($p$ = 0.63004) |
| Nerve Tibial | 0 (0.00%) | - |
| Ovary | 1 (0.03%) | 12.50 ($p$ = 0.0768929) |
| Pancreas | 0 (0.00%) | - |
| Pituitary | 0 (0.00%) | - |
| Prostate | 2 (0.05%) | 1.39 ($p$ = 0.421922) |
| Skin Not Sun Exposed Suprapubic | 3 (0.08%) | 3.12 ($p$ = 0.0730636) |
| Skin Sun Exposed Lower leg | 3 (0.08%) | 5.45 ($p$ = 0.0184417) |
| Small Intestine Terminal Ileum | 1 (0.03%) | 9.09 ($p$ = 0.104185) |
| Spleen | 1 (0.03%) | 0.47 ($p$ = 0.882392) |
| Stomach | 1 (0.03%) | 2.94 ($p$ = 0.288275) |
| Testis | 2 (0.05%) | 0.96 ($p$ = 0.617861) |
| Thyroid | 0 (0.00%) | - |
| Uterus | 3 (0.08%) | 0.93 ($p$ = 0.622227) |
| Vagina | 0 (0.00%) | - |
